# Single-cell mitochondrial lineage tracing decodes fate decision and spatial clonal architecture in human hematopoietic organoids

**DOI:** 10.1101/2023.09.18.558215

**Authors:** Yan Xue, Junhao Su, Yiming Chao, Lu Liu, Xinyi Lin, Yang Xiang, Mun Kay Ho, Zezhuo Su, Junyi Chen, Zhuojuan Luo, Chengqi Lin, Ruibang Luo, Theo Aurich, Jianfeng Wu, Kelvin Sin Chi Cheung, Yuanhua Huang, Joshua WK Ho, Ryohichi Sugimura

**Affiliations:** School of Biomedical Sciences, Li Ka Shing Faculty of Medicine, The University of Hong Kong, Pokfulam, HKSAR; Laboratory of Data Discovery for Health Limited (D24H), Hong Kong Science Park, HKSAR; Centre for Translational Stem Cell Biology, HKSAR; Department of Orthopaedics and Traumatology, School of Clinical Medicine, Li Ka Shing Faculty of Medicine, The University of Hong Kong, Pokfulam, Hong Kong SAR, China; Jiangsu Provincial Key Laboratory of Critical Care Medicine, Southeast University, Nanjing, China; Jiangsu Province Hi-Tech Key Laboratory for Biomedical Research, Southeast University, Nanjing, China; Department of Computer Science, School of Computing and Data Science, The University of Hong Kong; Heidelberg University Hospital, Germany; Department of Diagnostic Radiology, School of Clinical Medicine, Li Ka Shing Faculty of Medicine, The University of Hong Kong, Pokfulam, Hong Kong SAR, China; Department of Statistics and Actuarial Science, Faculty of Science, The University of Hong Kong, HKSAR

## Abstract

The ability to perform lineage tracing at the single-cell level is critical to reconstructing dynamic transitions during cell differentiation. However, prospective tracing approaches inevitably encounter outstanding challenges including barcoding precision, barcode diversity, and detection efficiency, which can skew inferred lineage relationships. Human pluripotent stem cells (hPSCs) even face risks of DNA-damage-induced toxicity-related cell death. We explored the use of naturally occurring somatic mutations in mitochondrial transcripts detected in single-cell RNA-seq as genetic lineage barcodes in hPSCs. In this study, we used an enrichment of scRNA-seq mitochondrial reads and a robust computational method to identify clonally relevant mitochondrial variants as endogenous genetic barcodes for clonal tracking of early embryonic hematopoiesis from hPSC. We modeled the development of embryonic tissues from hPSCs and delineated cell fate specification by integrating synthetic barcoding with mitochondrial lineage tracing. Using a biophysical model, we reconstructed the sequential transcriptional logic of fate specification and its underlying regulatory network. We further applied mitochondrial lineage tracing to spatial transcriptomics, which enabled us to identify the spatial clonal architecture of human embryonic organoids. Our analysis revealed that this spatial zonation was orchestrated by NOTCH-mediated crosstalk between stromal cells and hematopoietic progenitors. Our multi-modal framework links clonal dynamics with niche-specific fate decisions, providing a generalizable approach to dissecting tissue organization in development and disease. This study underscores the utility of mitochondrial variants as endogenous markers for high-resolution spatial clonal tracking in stem cell-derived organoid models of human development.

## Introduction

Tracing the progeny of individual cells is essential for understanding cell-division dynamics and fate decisions during embryonic development, stem cell differentiation, and disease progression. Prospective lineage tracing systems have been widely applied to study embryogenesis, hematopoiesis, neural development, and cancer biology (Bowling et al., 2020; Chan et al., 2019; Kalhor et al., 2018). Integrating genetic barcodes with single-cell transcriptomics enables simultaneous lineage reconstruction and state characterization (Baron & van Oudenaarden, 2019; Weinreb et al., 2020). However, technical limitations—including barcode homoplasy, restricted diversity, and detection sensitivity—compromise lineage resolution and accuracy. Current synthetic barcoding systems, such as LARRY, face inherent trade-offs: while enabling high-throughput clonal tracking through expressed DNA tags, they generate artifacts like spurious multi-progenitor labelling (Wang et al., 2022). Furthermore, dependency on multi-locus genetic modifications risks perturbing native cell behaviours— sustained CRISPR-Cas9 activity, for instance, can induce cytotoxicity in human pluripotent stem cells (Ihry et al., 2018). Such limitations underscore an unmet need for non-invasive, high-fidelity lineage recorders.

Retrospective lineage tracing, leveraging endogenous somatic mutations, offers a compelling alternative by eliminating artificial manipulation. Mitochondrial DNA (mtDNA) mutations are particularly advantageous due to their high mutation rate, heteroplasmic segregation, and compatibility with single-cell assays (Lareau et al., 2021; Ludwig et al., 2019). Recent advances in mitochondrial variant enrichment (MAESTER; Miller et al., 2022) and identification algorithms (MQuad; Kwok et al., 2022) now enable precise lineage tracing at scale. Yet, effectiveness of mitochondrial variants as lineage tracing marker to characterize the spatial arrangement of cell states and phenotypic transition haven’t been explored.

Spatial transcriptomics has significantly enhanced our understanding of developmental hierarchies, cellular plasticity and diverse tissue microenvironment in both healthy and diseased tissues (Gulati et al., 2025). The proper functioning of tissues relies on intricate interactions between cells, creating complex and dynamic cellular ecosystems (Palla, Fischer, et al., 2022). Since cells often have limited mobility within tissues, their physical proximity often indicates relatedness and their differentiation can be influenced by signals from neighbouring cells and morphogenetic gradients (Fischer et al., 2023). Tissues maintain stability during development through the coordinated efforts of various cell types, each with specialized roles and functions. Studying the spatial context of cell-fate decision-making can aid in distinguishing between intrinsic and extrinsic factors that impact cell differentiation and developmental processes (Lohoff et al., 2022; Tyser et al., 2021; Van Den Brink et al., 2020). Current spatial profiling can also capture substantial auxiliary information that influences cell fate, lineage, and differentiation (Erickson et al., 2022; Lomakin et al., 2022; Ru et al., 2023; Seferbekova et al., 2023).

Here, we leveraged naturally occurring mitochondrial somatic mutations from single-cell RNA-seq as endogenous genetic barcodes for lineage tracing in human pluripotent stem cell (hPSC)-derived embryonic organoids (HEMOs). By integrating this approach with synthetic barcoding, we reconstructed embryonic tissue development and delineated early cell fate choices. Furthermore, coupling mitochondrial clonal tracking with spatial transcriptomics revealed the coordinated emergence of hematopoietic cells and their supportive niche cells from common progenitors within yolk sac-like tissues. We further demonstrated that this spatially restricted clonal architecture is shaped by NOTCH-mediated stromal-progenitor crosstalk. Together, our findings establish mitochondrial variants as a powerful tool for resolving high-resolution spatial clonal dynamics in human developmental models.

## Results

### Distinct early fate choices and trajectories uncovered by lineage tracing in hematopoietic organoids

A significant challenge in stem cell haematopoiesis lies in linking molecular variations within progenitor cells to their ability to produce mature cell types. In this study, we employed expressed DNA barcodes to track transcriptomes clonally over time, applying this method to investigate hematopoietic clonal dynamics in HEMO. We infected M1-hPSCs with LARRY, which employs a barcoding system detectable by single-cell RNA sequencing (scRNA-seq) for clonal labelling, to investigate hematopoiesis within our HEMOs (Fig. 1a). Our aim is to track the dynamic process of blood regeneration in the human embryonic stem cell-derived organoid model, where hematopoietic stem and progenitor cells give rise to various blood cell lineages. Due to the 3D structure of the organoid, it is challenging for us to separate half of the cell population as the "early stage" and allow the remaining half to undergo further divisions to become "late stage" cells. Instead, we barcoded two batches of stem cells and harvested the D4 and D8 organoids respectively to generate early and late stage cells, enabling us to explore the lineage dynamics in HEMO hematopoiesis.

**Fig. 1.**
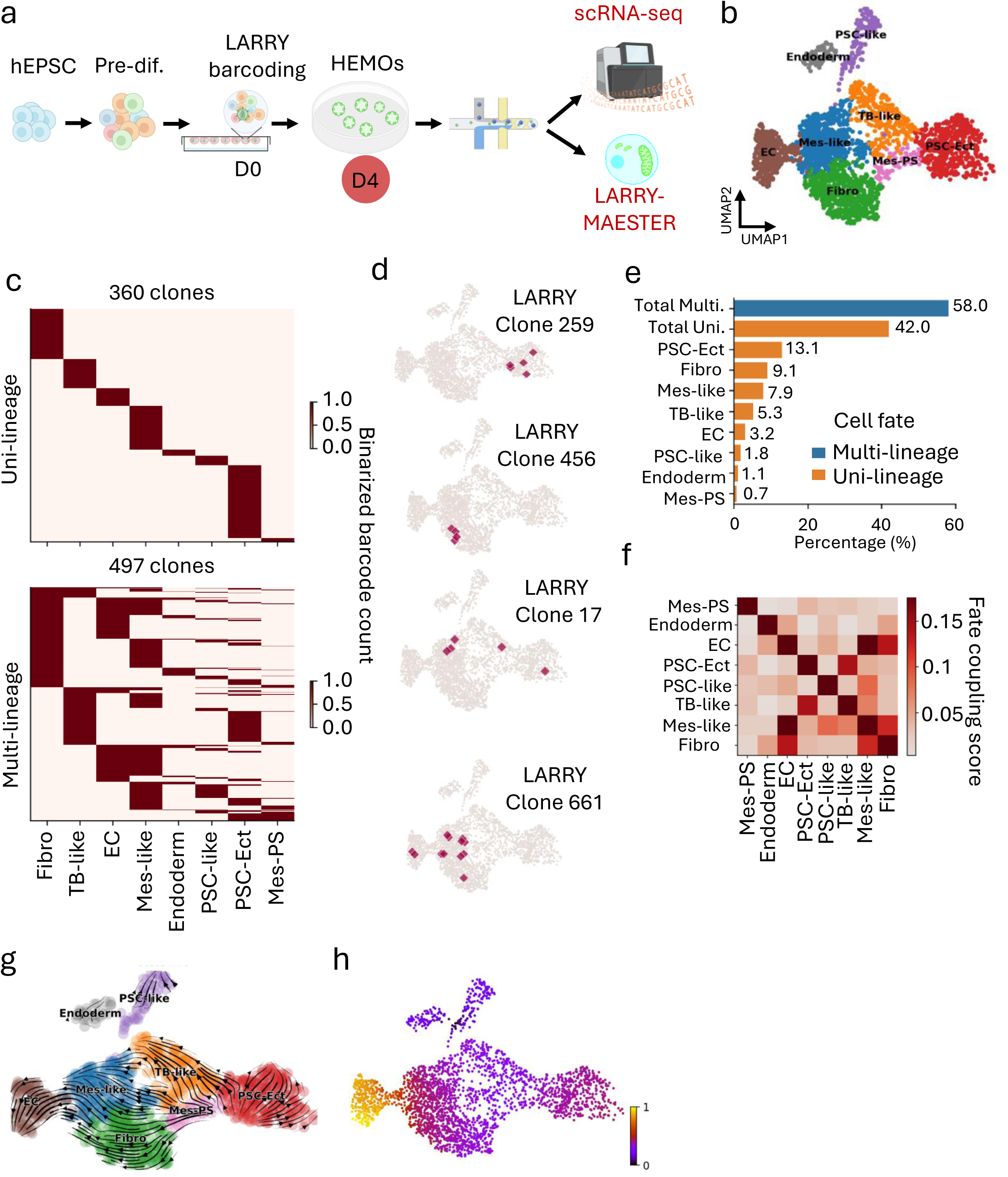
Lineage tracing reveals distinct cell fates during the early development of HEMOs (D4). a, Overall study design. Pre-differentiated human pluripotent stem cells were barcoded using LARRY on D0. Barcoded HEMOs harvested at D4 were subjected to single-cell RNA sequencing and mitochondrial variant enrichment to generate LARRY-MAESTER libraries. b, UMAP plot illustrates the transition of pluripotent cells into three distinct germ layers (ectoderm, mesoderm, endoderm) and specialized lineages. Key cell types include PSC-like, PSC-Ect, TB-like, Mes-PS (mesoderm primitive streak/cardiac mesoderm-like cells), Mes-like, EC, and Fibro. c, Heatmaps showing clonal fate predictions generated by CoSpar. Each row represents a LARRY clone, and each column corresponds to a specific lineage. d, UMAP plots showing the example clones with uni-(top two) or multi-lineage (bottom two) characteristics. e, Bar chart showing the proportion of clones in uni- and multi-lineages. f, Cell fate coupling revealed by CoSpar. The heatmap highlights coupling between Mes-like and EC as well as Fibro. g, RNA velocity field describes the fate decisions of major HEMO lineages in LARRY dataset. The velocity field is projected onto a PCA plot with arrows indicating the local average velocity evaluated on a regular grid. RNA velocity was estimated without cell or gene pooling. Mes-like derivatives demonstrated a strong bias for endothelial fate. Mes-PS cells were overwhelmingly likely to end up as Fibro. h, Pseudotime analysis reveals that Mes-like cells represent a transitional state, while EC cells correspond to a late developmental stage.

At the early stage of HEMO hematopoiesis (D4), we captured a diverse cell population, including endoderm, pluripotent stem cells-like (PSC-like), pluripotent stem cells with ectoderm specification (PSC-Ect), trophoblast-like cells (TB-like), mesoderm primitive streak/cardiac (Mes-PS), mesoderm-like cells (Mes-like), endothelial cells (EC) and fibroblasts (Fibro) (Fig. 1b and Fig. S1a). This suggests that our HEMO recapitulates key developmental processes, where pluripotent cells transition into three distinct germ layers (ectoderm, mesoderm, endoderm) and specialized lineages. The strong intra-cell-type connectedness (Weng et al., 2024) not only validated our clustering results but also provided additional evidence for the dynamic differentiation continuum within the HEMO system (Fig. S1b). LARRY barcoded a total of 4334 cells, grouping them into 2453 clones. We noted that 65% of the LARRY clones are one cell clones (Fig. S1c), aligning with the expected distribution of clone size (Weinreb et al., 2020). After filtering out the one cell clones, the remaining cells were assigned to 857 clones based on distinct LARRY barcodes (Fig. S1d). The clone sizes ranged from 2 to 15 cells. Notably, 58% of LARRY clones (481 clones) are two-cell clones. The number of large clones was relatively small, with only 15 clones (2%) containing 10 or more cells (Fig. S1d).

CoSpar analysis (Wang et al., 2022) revealed distinct cell fate in LARRY barcoded clones (Fig. 1c-g). Among the LARRY clones analysed, 42% (360 clones) were lineage specific, encompassing endoderm, PSC-like, Mes-PS, PSC-Ect, TB-like and Fibro lineages (Fig. 1c, d). These uni-lineage clones, such as LARRY Clone 259 and 456, spatially confined to a distinct region with the UMAP embedding (Fig. 1f, top panel), suggesting that they exhibit distinct lineage commitments and differentiation potentials. Notably, a significant proportion of LARRY clones spanned multiple lineages (Fig. 1c-e), displaying divergent spatial distributions in the UMAP embeddings (Fig. 1f, bottom panel). For instance, LARRY Clone 17 gave rise to PSC-Ect, Mes-like, and TB-like cells, and Clone 661 produced EC, Mes-like, and Fibro cells (Fig. 1f, bottom), demonstrating the multipotency of their progenitors.

Our lineage coupling analysis identified non-random interactions. Mes-like exhibited the strongest coupling with EC and Fibro lineages (Fig. 1g). Consistently, the dynamics of gene expression as captured by RNA velocity revealed pronounced directional flows converging toward each lineage in early HEMOs (Fig. 1h). The reconstructed velocity field mapped fate decisions across the principal lineages. Mes-like derivatives exhibited a strong primary bias toward EC specification (Fig. 1h). These dynamic patterns aligned with the clonal relationships in Fig. 1g. Furthermore, Mes-PS populations showed a dominant differentiation trend toward the Fibro lineage. Pseudotime ordering positions the Mes-like lineage as a progenitor state within a differentiation continuum that culminates in the acquisition of a mature EC identity (Fig. 1i). Collectively, these multi-faceted analytical approaches provide a coherent and dynamic portrait of early cell fate decisions, underscoring the central orchestrating role of mesenchymal progenitors in HEMO early development.

### A biophysical model reconstructs the sequential transcriptional logic of fate spacification

To reconstruct the dynamic trajectory of cell fate specification within our human embryonic hematopoietic organoids, we employed cell2fate (Aivazidis et al., 2025), a Bayesian framework that infers transcriptional dynamics through interpretable RNA velocity modules. This analysis identified a series of sequentially activated gene expression modules that delineate a complete developmental hierarchy, from a pluripotent progenitor state to the emergence of mature hematopoietic lineages and their requisite supportive niche (Fig. 2).

**Fig. 2.**
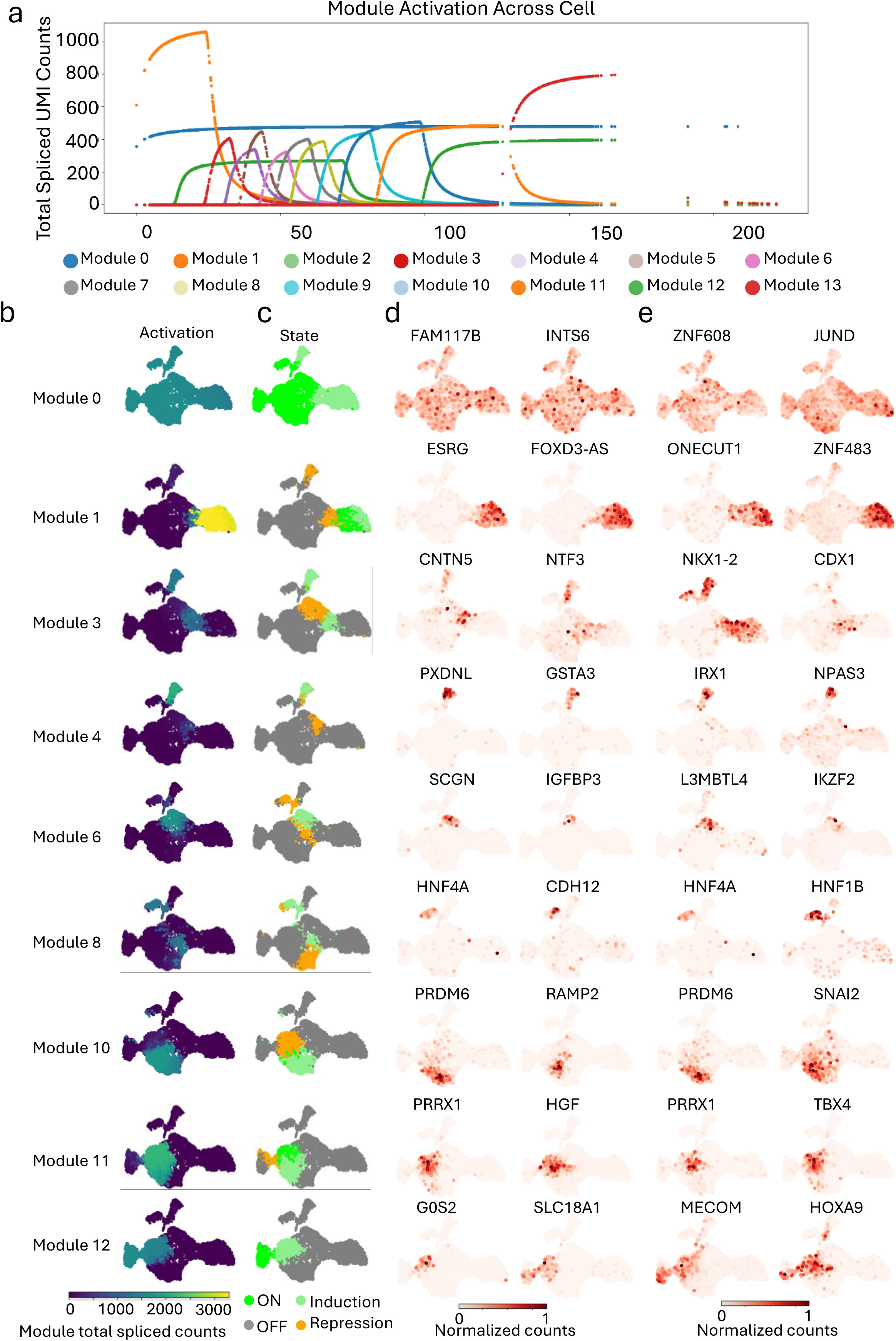
Module decomposition of cell2fate uncover coupled lineage specification in a human hematopoietic organoid. a, Total spliced mRNA abundance attributable to each cell2fate module across pseudotime. b, Module activity scores per cell, quantified as the fraction of spliced counts explained by each module. c, Expression patterns of key lineage-specific modules during late hemogenic endothelium development. d, Top two marker genes for each module, identified based on the proportion of their transcription rate attributed to the module. e, Key regulatory transcription factors (TFs) within each module, defined as module marker genes with known TF activity.

The initial state of the system was defined by Module 0, which was ubiquitously active across all early cell types (Fig. 2b,c). This module was characterized by high expression of marker genes FAM117B and INTS6, and driven by transcription factors ZNF608 and JUND, indicative of a primordial, proliferative, and metabolically active foundation from which all subsequent lineages emerge (Fig. 2d,e).

Following the initial ectodermal commitment (Module 1), Module 3 became specifically active in both Mes-PS and a distinct pluripotent (PSC-like) population. This module was characterized by marker genes CNTN5 and NTF3, and regulated by transcription factors NKX1-2 and CDX1, suggesting a role in priming mesodermal potential while maintaining pluripotency.

Subsequently, Module 4 expression became restricted solely to the PSC-like population, with loss of Mes-PS activity. This shift was marked by the expression of genes PXDNL and GSTA3, under the regulatory control of IRX1 and NPAS3, indicating a refinement of the pluripotent state, potentially toward a more specialized or transient progenitor identity.

Later stages of differentiation were driven by the activation of lineage-specific modules. Module 6 was uniquely associated with TB-like cells, defined by markers SCGN and IGFBP3, and controlled by the factors L3MBTL4 and IKZF2. Module 8 activation specifically heralded endoderm specification, evidenced by the expression of the classic marker HNF4A alongside CDH12, and directed by the key endoderm-regulatory transcription factors HNF4A and HNF1B.

Further lineage diversification was marked by the activation of Module 10, which drove the emergence of a Fibro identity. This state was characterized by genes PRDM6 and RAMP2, and transcription factors PRDM6 and SNAI2, the latter indicative of potential roles in epithelial-to-mesenchymal transition. Concurrently, Module 11 activation specified Mes-like cells, marked by PRRX1 and HGF expression and governed by the regulators PRRX1 and TBX4. Finally, the acquisition of an EC fate was propelled by Module 12, defined by the expression of G0S2 and SLC18A1 and orchestrated by the transcription factors MECOM and HOXA9.

This modular series of gene expression activations and restrictions outlines a hierarchical roadmap of cell fate decisions, from a multipotent progenitor state to the establishment of distinct endodermal, mesenchymal, and endothelial lineages.

To directly link the molecular mechanisms identified by cell2fate to our empirical clonal observations (Fig.1c-f), we investigated whether the inferred gene expression modules could provide a mechanistic basis for the non-random lineage couplings uncovered earlier. Remarkably, the model offered a clear explanatory framework for these key findings:

Our lineage coupling analysis identified non-random interactions, with the Mes-like lineage exhibiting the strongest coupling to both EC and Fibro lineages (Fig. 1g). This bifurcated potential was further elucidated by distinct gene module activities: Module 10 was specifically activated in Fibro and repressed in Mes-like, driving stromal specification (Fig. 2). Conversely, Module 11 showed strong activation in Mes-like with parallel induction in Fibro, supporting a shared progenitor–stromal relationship (Fig. 2). Importantly, Module 12 was specifically activated in EC with clear induction in Mes-like but absent in Fibro, highlighting a privileged differentiation path from mesenchymal progenitors to endothelium (Fig. 2). Finally, Module 13 was exclusively highly expressed in mature EC, marking terminal endothelial differentiation.

These modular patterns were consistent with RNA velocity trajectories, which revealed pronounced directional flows from Mes-like toward EC specification (Fig. 1h), corroborating the role of Modules 12 and 13 in endothelial commitment (Fig. 2). The strong bias of Mes-PS toward Fibro was also reinforced by Module 10-specific activation. Pseudotime analysis positioned the Mes-like population as a central progenitor within a continuous lineage continuum culminating in EC identity (Fig. 1i), aligning with the transient co-expression patterns in Modules 11 and 12 during fate restriction (Fig. 2).

Collectively, these multi-modal data integrate regulatory modules with fate dynamics, providing a coherent model of early lineage diversification in which modular gene activation and repression direct bifurcated trajectories from multipotent mesenchymal progenitors to stromal and endothelial fates.

### Clonal expansion and restricted fate potential characterize late-stage organoid maturation

At D8, HEMOs transitioned into a more developmentally advanced and stable state, characterized by clearly segregated lineage compartments and the emergence of novel progenitor populations, including erythro-myeloid progenitors (EMP) and cardiac mesoderm-like cells (CM-like) (Fig. S2a,b). Analysis of the D8 LARRY dataset (2,423 cells assigned to 1,706 clones) revealed that 50% of clones (845/1,706) were one cell clones (Fig. S2c,d). After excluding these singletons, the remaining 1,578 cells clustered into 347 clones (Fig. S2e), indicating reduced clonal diversity as hematopoiesis progressed. Notably, late stage HEMO exhibited significant clonal expansion: the largest clone contained 29 cells—double the maximum observed in early stage—and 10% of clones (38/347) exceeded 10 cells, a marked increase compared to earlier timepoints (Fig. S2e). This shift suggests the stabilization and dominance of specific clones during later developmental stages. Additionally, we identified lineage specific LARRY clones at D8, including PSC-like, CM-like, PSC-Ect, ectoderm-like cells (Ect-like), endoderm, and EMP lineage specific clones (Fig. S3a). We also observed 60% LARRY clones were multi-lineages (Fig. S3c). Similarly, these uni-lineage clones, such as LARRY Clone 185 and 188, spatially confined to a distinct region with the UMAP embedding (Fig. S3c, top panel), suggesting that they exhibit distinct lineage commitments and differentiation potentials. The multi-lineage LARRY clones displayed divergent spatial distributions in the UMAP embeddings (Fig. S3c, bottom panel). For instance, LARRY Clone 71 gave rise to Fibro and EC cells (Fig. S3c, bottom), demonstrating the multipotency of their progenitors.

Lineage coupling analysis revealed a significant reduction in broad non-random interactions compared to D4, reflecting enhanced lineage restriction and specialization (Fig. S3d). Despite this overall segregation, strong coupling between EC and Fibro lineages persisted (Fig. S3d), reaffirming a conserved bifurcated differentiation path toward vascular and stromal fates—a hallmark of multipotent mesenchymal progenitor activity. Notably, a moderate but distinct clonal coupling was detected between EMP and EC lineages (Fig. S3d). This interaction suggests a developmental kinship or shared progenitor trajectory between endothelial and erythro-myeloid lineages, supporting the notion of hemogenic endothelial-like activity or a vascular-affiliated origin for a subset of EMPs within the HEMO model.

RNA velocity analysis further corroborated these findings showing refined and pronounced directional flows toward well-defined terminal lineages (Fig. S3e). The reconstructed velocity fields illustrated a maturation of fate decisions: the EC lineage demonstrated heightened maturity and structural formation (Fig. S3e). In contrast to its strong EC-specification bias at D4, the Mes-like population had largely shifted its differentiation trajectory by D8, with only a minor subset of cells retaining potential to contribute to the Fibro lineage (Fig. S3E). Concurrently, PSC-Ect lineages showed committed flow toward ectodermal fates (Fig. S3e). This supports the interpretation that earlier pluripotent or bipotent states have resolved into more restricted identities by D8. The pseudotime trajectory positions EC cells as a late-stage population, aligning with their phenotypic maturation into functional endothelial cells (Fig. S3f). Together, these results indicate that D8 represents a stage of increased lineage resolution and structural maturity within HEMOs. The persistence of EC–Fibro coupling— alongside the emerging EMP–EC relationship and the clear separation of other lineages— highlights the sustained role of mesenchymal and endothelial progenitors in supporting hematopoietic and stromal crosstalk even amid overall developmental progression toward compartmentalization.

### Benchmarking of mtDNA variant calling pipelines establishes MQuad for reliable clonal inference

Recent advancements have made it feasible to delve deeply into mtDNA for resolving clonal substructures and lineage tracing by implementing the MAESTER. To conclusively distinguish true clonal multilineage potential from technical artifacts and to verify the reconstructed differentiation trajectories, we integrated mitochondrial DNA (mtDNA) mutations as naturally occurring, endogenous somatic markers for lineage tracing. mtDNA mutations accumulate stochastically during cell division and are clonally inherited, making them ideal orthogonal markers to validate synthetic barcode-based clonal groupings. Unlike engineered barcodes, mtDNA variants are not subject to homoplasy from viral integration bias or limited barcode diversity, and their detection via single-cell RNA-seq allows simultaneous assessment of transcriptome and lineage history without additional experimental burden.

Bulk RNA-seq of expanded pluripotent stem cells (EPSCs, D0) and D18 HEMOs demonstrated substantial overlap in mtDNA variants between stages (Fig. 3a), supporting their utility as stable lineage markers. By tracking stem cell subsets marked by unique mtDNA variants, we aim to resolve multi-progenitor labeling events and refine clonal assignments (Fig. 3b). To operationalize this strategy, we designed a targeted mitochondrial variant enrichment assay in LARRY barcoded cells to generate LARRY-MAESTER (Fig. 1a), enabling high-confidence reconstruction of true clonal relationships independent of LARRY technical artifacts.

**Fig. 3.**
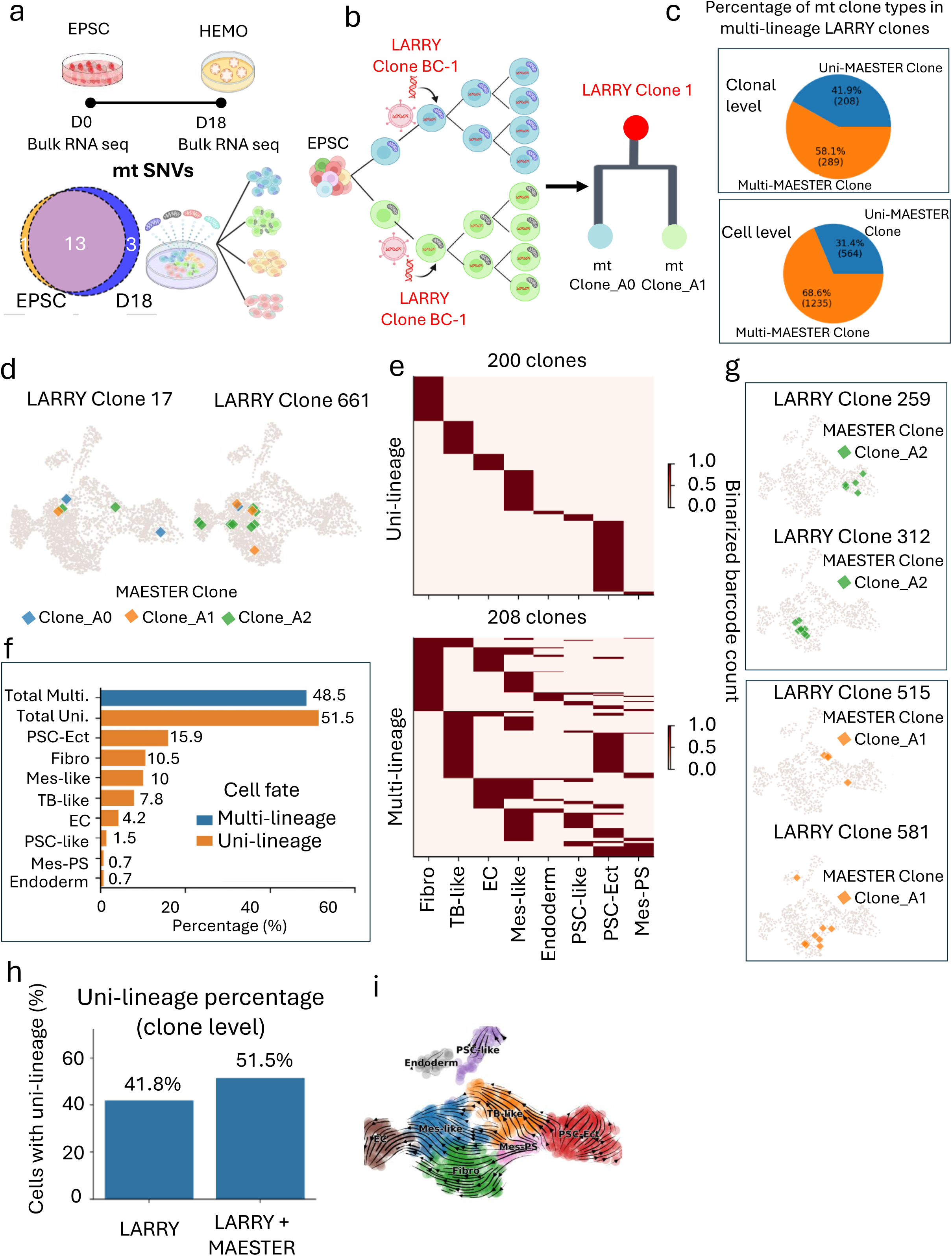
Mitochondrial variant barcoding resolves LARRY noise and uncovers cell fate decisions during early development of HEMOs (D4). a, Stable lineage markers: mtDNA variants. Bulk RNA-seq analysis reveals substantial overlap in mtDNA variants between EPSCs (D0) and D18 HEMOs, supporting their utility as stable lineage markers. b, Diagram shows integration of mtDNA variant barcoding can address LARRY barcode homoplasy, enhancing clonal resolution and lineage tracing accuracy. c, Barcode homoplasy in LARRY clones. The UMAP plot shows that barcode homoplasy results in LARRY clones, such as Clone 17 and Clone 661, consisting of multiple subclones defined by distinct mitochondrial variants. d, Heatmaps showing refined clonal fates after correction with mitochondrial variants. Each row represents a LARRY clone, and each column corresponds to a specific lineage. e, Bar chart showing the proportion of clones in uni- and multi-lineages after mitochondrial variant refinement. f, Chord diagram depicting the multi-lineage couplings after mitochondrial variant refinement. g, UMAP plots showing the example mitochondrial variant refined clones with uni-(top two) or multi-lineage (bottom two) characteristics. h, The bar chart demonstrates that mitochondrial variants significantly increase the proportion of cells committed to specific fates. i, The RNA velocity field elucidates the fate decisions of major hematopoietic lineages in the LARRY-MAESTER dataset, which exhibits similar, yet distinct cell state transitions compared to the original LARRY dataset.

Informative mitochondrial variants are essential for establishing clonality. However, the absence of a definitive gold standard computational pipeline for mtDNA variant calling and clonal reconstruction presents a significant obstacle to leveraging mtDNA for lineage tracing inference. Presently, two pipelines stand out for mtDNA analysis in single cell sequencing data: MQuad and maegatk. Prior to applying these techniques to actual biological datasets, it is essential to conduct a foundational comparison of two available analysis pipelines. This comparative analysis will facilitate an informed decision on the optimal pipeline for analysing real data.

To facilitate the benchmarking work, we generated three Chromium-MAESTER datasets of HEMOs at D8, D15 and D18. We also included a hematopoietic dataset from the original MAESTER study. To ensure accurate clonality assessment, we initially benchmarked MQuad and maegatk computational pipelines in four HEMO datasets (Fig. S4). Subsequent variants identified and clonal assignments derived from processing using both pipelines were used to evaluate the strengths and weaknesses of each pipeline (Fig. S5-S22).

In the MQuad pipeline, the number of variants identified in each dataset fell between a similar range of 41 to 44 (Fig. S4b and S5) comparing the three organoid datasets. We observed sharp increases in the cumulative distribution of ΔBIC in all three stages (Fig. S 6) which further justified the rationale behind determining the cutoff based on a knee point and confirmed the informativeness of the identified mitochondrial variants with high sensitivity and specificity. There were significantly fewer variants identified in the hematopoietic dataset (19 variants) (Fig. S4b and S7); however, this may be attributed to the size of the dataset being significantly smaller than the other three. We defined 3, 5, 4 and 4 clones by unique combinations of barcodes and reconstructed lineage in D8, D15, and D18 HEMOs, as well as the hematopoietic dataset respectively (Fig.S5 and S7).

For maegatk pipeline, with the suggested minimum 3 UMI reads parameter, cells with more than 1% VAF for any identified variant were selected and subset for the subsequent clonal assignment step (Fig. S8-S12). The variant allele frequency of the selected cells and the variants identified were applied to the R package, clValid (Brock et al., 2008). Using the internal validation method, metrics such as connectivity, Dunn index and silhouette width were applied for validation (Fig. S13a, S14a, S15a and S16a). In all four datasets, the hierarchical clustering method using Euclidean distance metrics was suggested as the best method, this was achieved using the eclust function from the factoextra R package (Kassambara & Mundt). The ward. D clustering method was applied to accomplish minimal variance between clusters. To aid visualization, the branches of each cluster in the dendrogram were highlighted in a different colour and can be easily interpreted in the simplified dendrogram (Fig. S13b, S14b, S15b and S16b). The number of clonal clusters identified in each of the four datasets is highly varied.

The primary question in evaluating both pipelines remains in their respective capabilities in identifying informative mtDNA variants which would be useful in downstream analysis of clonal reconstruction for meaningful lineage inference and biological interpretation. When the list of mtDNA variants identified by the two pipelines for each dataset was compared, there was a significantly higher number of variants identified by the maegatk pipeline. However, few overlapping or intersecting variants were identified by both pipelines (Fig. S17). This is interesting in understanding that the primary sequencing data used as input for each pipeline is completely identical and yet the results produced are highly varied.

Heatmaps of scaled VAF values were constructed to visualize the relationship between the variants identified and the clonal structure (Fig. S18-S21). In clones assigned using the MQuad pipelines, there was a small but significant number of variants seen to be highly specific to certain clonal clusters (Fig. S18a, S19a, S20a and S21a). This was not observed in the comparatively scattered heatmaps for clones identified using the maegatk pipeline (Fig. S18b, S19b, S20b and S21b). This preliminary result was highly corroborative with the heatmaps produced using the seriation R package which visually represents the Euclidean distance matrix calculated from the scaled VAF matrix from each pipeline. This suggests that the clones identified by maegatk do not show a clear distinction between clonal clusters using the information supplied by the VAF values of identified variants.

To quantitatively assess the clonal clusters identified using mtDNA informative variants from both pipelines, the Davies-Bouldin’s (DB) index was applied to analyse the clusters identified using the informative variants for each pipeline (Fig. S22). A minimal value of DB index value is indicative of strong clustering, and this was observed in MQuad inferred clonal clusters (Fig. S22). The DB index value derived from the maegatk variant inferred clonal clusters were often more than double that in the MQuad clones. The outcome from cluster validation of clonal clusters identified through mtDNA single nucleotide variation demonstrated that the variants discovered using the MQuad pipeline are more informative than maegatk. As a result, the MQuad pipeline was used in our study for variant calling and clonal inference.

### Orthogonal mitochondrial tracing validates and refines clonal dynamics in HEMOs

To resolve potential technical artifacts in LARRY-based clonal tracing, we combined LARRY barcoding with mitochondrial variants profiling using MAESTER enrichment to generate LARRY-MAESTER libraries (Fig. 1a). From D4 and D8 HEMO LARRY-MAESTER libraries, MQuad identified 35 and 29 high-confidence mtDNA variants, respectively. The uneven distribution of mtDNA variants across various cell types suggests the potential to distinguish clonality (Fig. S23a). To comprehensively capture cellular correlations at the early stage (D4), we utilized all mtDNA variants for clone assignment, identifying three clones based on unique mitochondrial barcode combinations and lineage reconstruction (Fig. S23b). Notably, the mtDNA variants 6321G>A, 6322G>C, 6233A>C, and 6234G>A exhibited high VAF and distinctly marked Clone A0, while another mtDNA variant, 7402C>T, with high VAF, specifically identified Clone A2 (Fig. S23b). Interestingly, most cells in Clone_A0 were characterized by mtDNA variant 6321G>A, Clone_A1 predominantly exhibited mtDNA variant 6699G>A, and Clone_A2 was primarily distinguished by mtDNA variant 7402C>A (Fig. S23c, d). Neighbourhood analysis revealed strong intra-clone connectedness, highlighting the ability of mitochondrial variants to accurately delineate robust clonal identities (Fig. S23e).

Comparative analysis revealed a striking discordance (52%) between clonal identities defined by LARRY barcodes and those resolved by endogenous mitochondrial variants (Fig. S23f). Among multi-lineage LARRY clones, 208 clones (41.9%) consisted of cells sharing the same mitochondrial variant, providing strong evidence for a pluripotent progenitor origin (Fig. 3c). Conversely, 289 multi-lineage LARRY clones (58.1%) contained more than one mitochondrial clone, indicating they were false positives resulting from LARRY barcode homoplasy (Fig. 3c). At the cellular level, only 31.4% of cells could be confidently assigned to pluripotent progenitors based on congruent mitochondrial evidence; the remainder were deemed technical artifacts (Fig. 3c). Illustrating this discrepancy, LARRY clones 17 (containing PSC-Ect, TB-like, and Mes-like cells) and 661 (with Mes-like, EC, and Fibro cells) each comprised three distinct mitochondrial subclones (Fig. 3d). This recurrent "one-to-many" relationship underscores two critical limitations: pervasive genetic heterogeneity within synthetically defined clones and the high frequency of barcode homoplasy in the LARRY system. The presence of distinct mtDNA variant profiles among cells within a single LARRY clone confirms that such groups are not monoclonal but represent aggregates of biologically independent clones. Given the high conservation of mitochondrial variants between stem cell progenitors and late-stage (D18) HEMOs in our bulk RNA analysis, we conclude that the observed discordance predominantly reflects intrinsic technical noise of the LARRY system rather than biological variation.

Notably, cells within the 48% concordant clones exhibited more distinct cell type boundaries in the UMAP plot, highlighting the enhanced resolution achieved by integrating endogenous mtDNA markers with exogenous barcoding (Fig. S24a). CoSpar analysis further revealed well-defined cell fates within these concordant clones (Fig. 3e). When mitochondrial variants were included, the proportion of uni-lineage clones increased, while the diversity of multi-lineage clones decreased, compared with LARRY only clones (Fig. 1d,e, 3e,f). For uni-lineage clones, consistent spatial distributions in UMAP embeddings were observed regardless of whether mitochondrial variants or LARRY barcodes were used. For instance, LARRY Clone 259, belonging to the PSC-Ect lineage, aligns specifically with mitochondrial Clone_A2, and LARRY Clone 312, representing the Fibro lineage, also maps exclusively to mitochondrial Clone_A2 (Fig. 3g).

By contrast, LARRY Clone 515, which spans both PSC-Ect and TB-like lineages, is consistently associated with mitochondrial Clone_A1, revealing its bi-potential nature (Fig. 3g, bottom panel). Similarly, LARRY Clone 581, encompassing PSC-Ect, Fibro, and endoderm lineages, is consistently linked to mitochondrial Clone_A1, highlighting its multi-potency during HEMO development (Fig. 3g, bottom panel). These results suggest that mitochondrial variants not only refine clonal classification but also provide insights into the underlying mechanisms governing lineage commitment and multi-lineage potential within organoid development.

Interestingly, in the concordant clones, we consistently observed the coupling between the cell fates of Mes-like and Fibro, as well as Mes-like and EC cells, further proved the conserved bifurcated differentiation path toward vascular and stromal fates—a hallmark of multipotent mesenchymal progenitor activity (Fig. S24b). Statistically, mitochondrial variants significantly increased the proportion of cells with individual cell fate when compared with LARRY barcoded cell population (Fig. 3h).

Comparing to the LARRY only clones, similar the reconstructed RNA velocity field was observed of mitochondrial refined LARRY clones delineating the clearer fate-determination patterns across principal HEMO lineages especially in Fibro lineage (Fig. 4i). Through likelihood-based computational screening, we identified putative driver genes governing lineage transitions (Fig. S24c). Phase-plane analysis integrating transcriptional bursting kinetics, velocity vectors, and expression dynamics revealed specific regulatory programs: ZEB2 mediated the terminal differentiation of Mes-like progenitors (blue) into mature EC (grey) via progressive activation (Fig. S24c). This bifurcated architecture indicates early stage lineage segregation during hematopoietic specification.

**Fig. 4.**
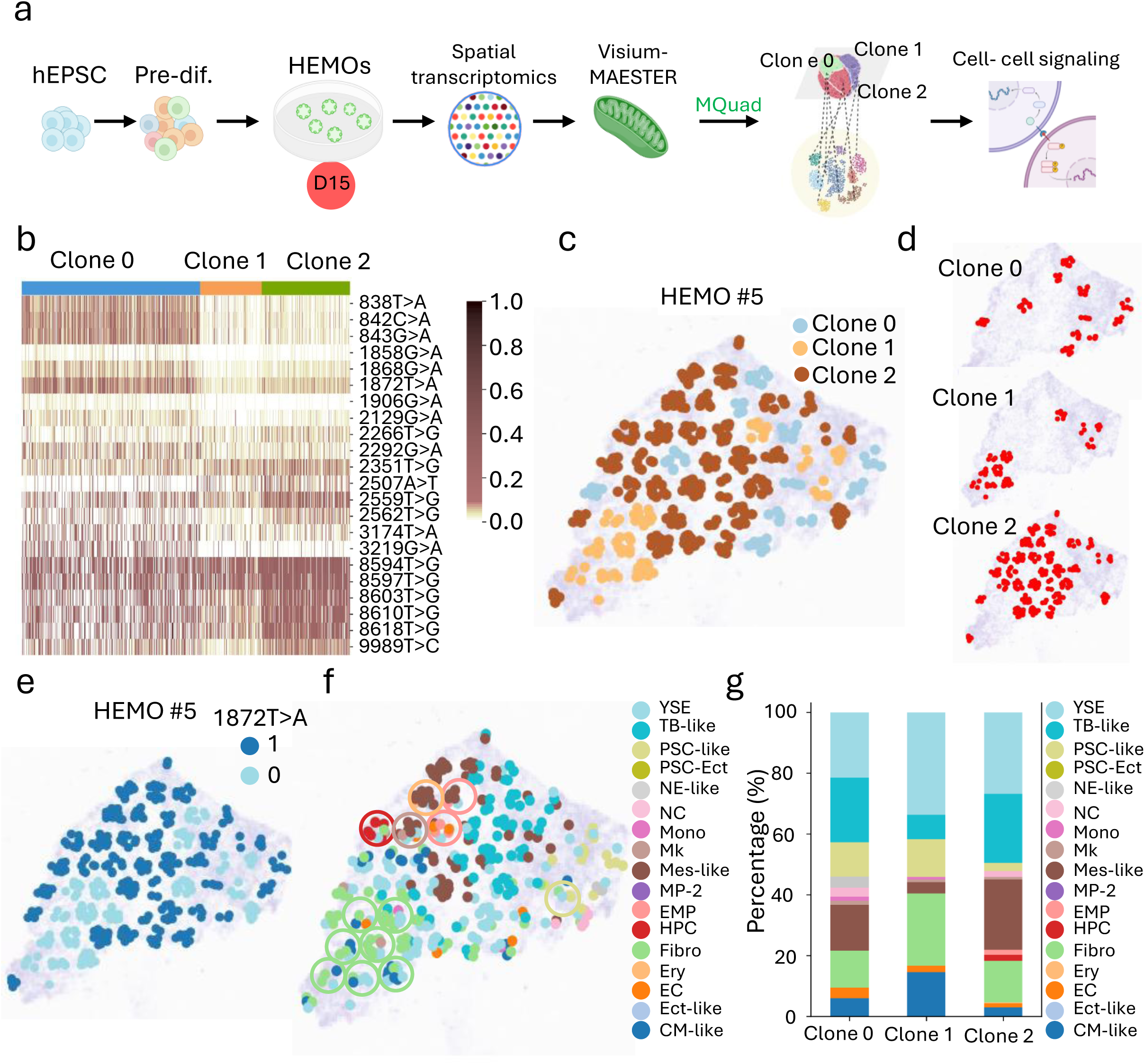
Mitochondrial variants based spatial clonal tracking in D15 HEMOs. a, Overall study design for mitochondrial variants based spatial lineage tracing. Pre-differentiated (Pre-diff.) human expanded pluripotent stem cells (hEPSC) were induced to form human embryonic hematopoietic organoids (HEMOs). HEMOs harvested at Day 15 (D15) were subjected to spatial transcriptomics and mitochondrial variant enrichment to generate Visium-MAESTER, enabling mitochondrial variants based spatial lineage tracing. b, Allele frequency heatmap shows the 22 informative mtDNA SNVs detected by MQuad in each clone. Each row is a variant, each column is a barcode. Heatmap color indicates the value of the allele frequency. c, Spatial architecture of 3 clones identified by mtDNA SNVs from MQuad in Visium-MAESTER library in HEMO#5. d, Spatial distribution of individual clones in HEMO #5, highlighting the distinct localization patterns of each clone. e, Spatial allelic frequency of mitochondrial SNV 1872T>A, as identified by SpatialDE, aligns strongly with the spatial distribution of Clone 0 and Clone 2 in HEMO #5. This demonstrates the correspondence between mitochondrial variation and clonal spatial patterning. f, Spatial transcriptomics map of HEMO #5, colored by clusters. The spatial organization of clonal niches is highlighted, with each niche circled and color-coded to indicate the dominant cell state present within that region. g, Bar plot shows multiple cell populations in HEMO #5 illustrating the increased formation of HPC and EMP from Clone 1 to Clone 2, as well as increased formation of Mk from Clone 2 to Clone 0 at late stage of hematopoiesis. YSE, yolk sac endoderm; TB-like, trophoblast-like cells; PSC-like, pluripotent stem cells-like; PSC-Ect, pluripotent stem cells with ectoderm specification; NE-like, neural ectoderm-like cells; NC, neural crest; mono, monocytes; Mk, megakaryocytes; Mes-like, mesoderm-like cells; MP-2, Myeloid progenitor 2; EMP, erythro-myeloid progenitor; HPC, hematopoietic progenitor cells; Fibro, fibroblasts; Ery, erythroid cells; EC, endothelial cells; Ect-like, ectoderm-like cells; CM-like, cardiac mesoderm-like cells.

In D8 HEMOs, clonal resolution with mitochondrial variants provides strong support for our conclusion that the system has reached a stable stage of lineage differentiation and structural maturity (Fig. S25, 26). The persistence of EC–Fibro coupling—alongside the emerging EMP–EC interaction and clear segregation among other lineages—underscores the continued contribution of mesenchymal and endothelial progenitors in facilitating hematopoietic and stromal crosstalk, even as the system progresses toward compartmentalization (Fig. S26).

### Spatial mtDNA clonal mapping unveils niche-driven fate patterning in organoids

To validate the ability of MAESTER-MQuad algorithm for accurate and reliable clonal tracking in spatial context, we first conducted spatial MAESTER experiments using chondrosarcoma samples. After enriching the mitochondrial transcriptome in the Visium library, MQuad identified 24 informative mitochondrial variants in the Visium-MAESTER dataset. Utilizing these informative variants, we assigned 1344 chondrosarcoma cells into 16 clones (Fig. S27). Notably, we observed that the mitochondrial variant 10310A>G effectively distinguished two major clones: Clone 5 and Clone 11. Clone 11 exhibited a high allelic frequency of 10310A>G, whereas Clone 5 did not show any variation at this position (Fig. S27).

When we examined the histoclonal relationship, we observed that Clone 5 was predominantly distributed on the left side of the histological slide, corresponding to the tumor-adjacent normal area (Fig. S28). In contrast, Clone 11 was primarily distributed on the right side of the histological slide, representing the neoplastic tumor area (Fig. S28). This suggests that mitochondrial variant 10310A>G is accumulated during the cancer progression process. Our Visium-MAESTER pipeline is a powerful approach for spatially distinguishing normal clones from neoplastic tumor clones, which provide valuable insights for cell states transition in cancer progression.

Furthermore, we utilized spatialDE (Svensson et al., 2018) to identify gene expressions that significantly depended on spatial coordinates in a non-linear and non-parametric manner. Consistent with our MQuad results, spatialDE identified the mitochondrial variant 10310A>G as a significant feature, indicating that the expression of 10310A>G was significantly dependent on its spatial location (Fig. S29). Based on these findings, we conclude that the mitochondrial variants identified by MQuad after MAESTER are informative for clonal tracking, and their expression is significantly correlated with their spatial location. Therefore, the clonality established using mitochondrial variants in our Visium-MAESTER pipeline effectively resolves the spatial architecture of clones.

In light of this validation, we applied spatial transcriptomics in combination with mitochondrial variants to dissect the spatial clonal architecture of hematopoietic and niche cells in D15 HEMOs (Fig. 4 and 5). We enriched mitochondrial transcripts from 10x Visium full-length cDNA and dissected their clonal structure using informative mtDNA variants identified by MQuad (Fig. 5a). Across all five HEMOs, we identified a total of 22 informative mtDNA variants (Fig. 4a). Among these variants, we found that most of the informative variants were C>T or T>G transitions (Fig. S30a). Using vireoSNP, we consistently identified three distinct clones based on these informative variants (Fig. 4b). Notably, there is a tendency for the numbers of variants to increase from Clone 1 to clone 2 and further to Clone 0, suggesting a developmental trajectory among the three clones (Fig. 4b). The signature variant 1868G>A, 1906G>A and 3219G>A are predominantly associated with Clone 0, whereas 2507A>T is primarily observed in Clone 1 and 2 (Fig. 4b). By contrast, Clone 1 exhibits a significantly lower number of variants compared to Clones 0 and 2 (Fig. 4b). Upon mapping the mtDNA variants to spatial locations, we observed that Clone 0 was primarily located at the edges of organoids, whereas Clone 2 was surrounded by Clone 1 across all five HEMO samples (Fig. 4c, d and Fig. S30b). Using SpatialDE, we found that the expression of 1872T>A was significantly dependent on spatial coordinates. Interestingly, the integrative spatial mapping of both clonality and mitochondrial variants revealed that the high-expression spatial regions of 1872T>A aligned well with Clone 0 and Clone 2, suggesting the informativeness of mitochondrial variants for dissecting spatial architecture in our organoids (Fig. 5e and Fig. S30c).

**Fig. 5.**
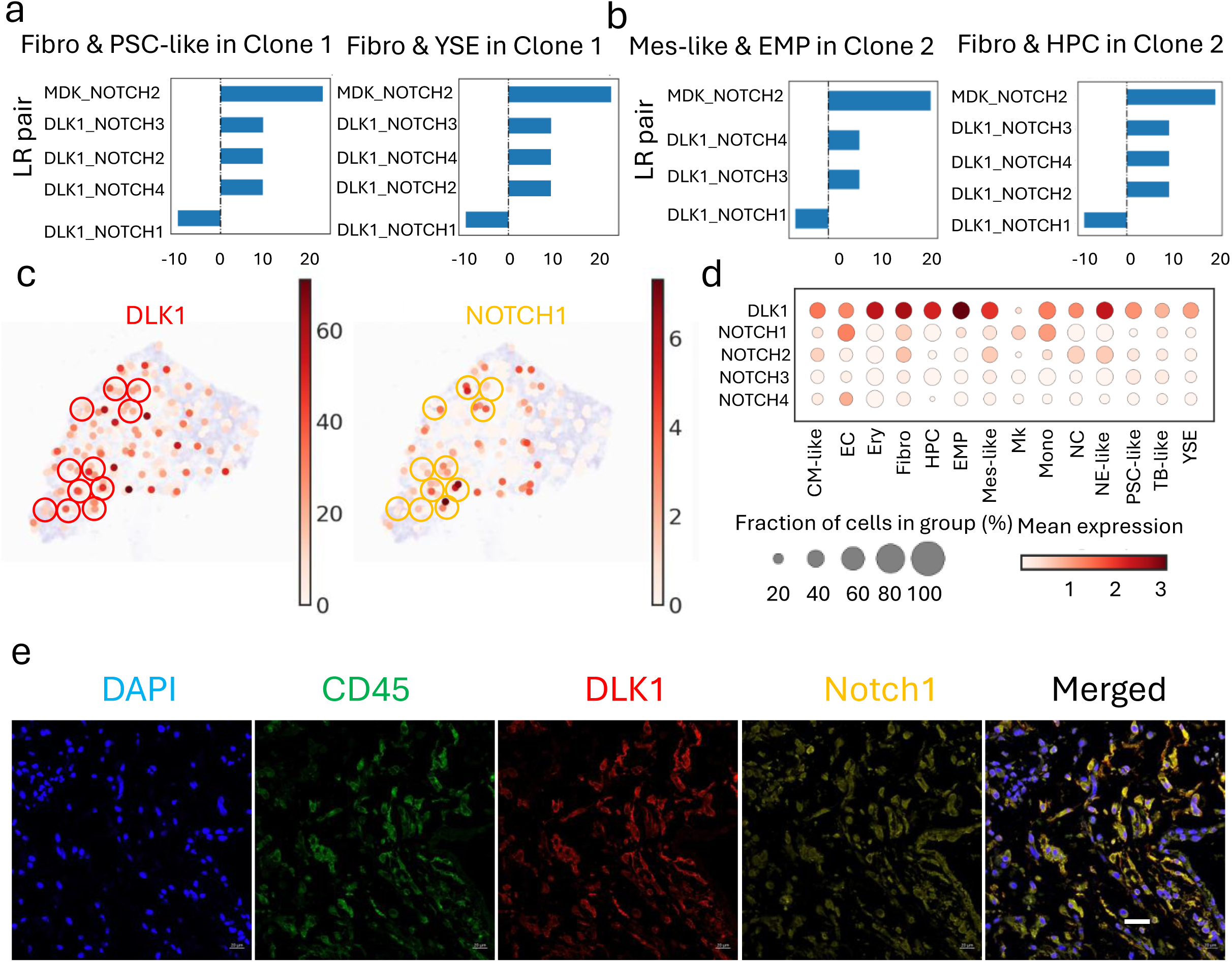
Intercellular signaling drives spatial phenotypic shifts in multicellular neighborhoods of HEMO clones at D15. a, Spatial cell-cell interaction analysis conducted by CellPhoneDB within Clone 1 between Fibro and PSC-like, as well as Fibro and YSE. Pairs with a mean value above 0 indicate the activation of the ligand-receptor pairs. Pairs with a mean value below 0 indicate the inhibition of the pairs. Among these, *DLK1*_*NOTCH1* exhibits high enriched score between Fibro and PSC-like, as well as Fibro and YSE in Clone 1. b, Spatial cell-cell interaction analysis conducted by CellPhoneDB within Clone 2 between Mes-like and EMP, as well as Fibro and HPC. *DLK1*_*NOTCH1* exhibits high enriched score between Mes-like and EMP, as well as Fibro and HPC in Clone 2. c, Gene expression pattern of *DLK1* and *NOTCH1* within Clone 1 and Clone 2 in HEMO #5 in a spatial slice. *DLK1* highly expressed in PSC-like and YSE within Clone 1 and in EMP and HPC within Clone 2. *NOTCH1* highly expressed in Fibro within both Clone 1 and Clone 2. *NOTCH1* also highly expressed in Mes-like in Clone 2. Circles indicate niches where *DLK1* (red) and *NOTCH1* (yellow) were co-expressed. d, Dot plot shows the expression of *DLK1* and *NOTCH1* across various cell types. e, Immunofluorescence staining of DLK1^+^ hematopoietic cells and Notch1^+^ non-hematopoietic cells. Scale bar = 20 µm.

We further aggregated and analysed the cell type composition for each spatial spot based on mtDNA information, revealing that Clones 0 and 2 exhibited greater cellular diversity than Clone 1 across all HEMOs (Fig. 4f, g and Fig. S30d, e). Due to the presence of large vacuolated regions in the tissue section of HEMO #2, and the significant reduction in cellular diversity observed in HEMOs #3 and #4, these samples were deemed insufficient to accurately represent the cellular differentiation characteristics of late stage HEMOs. Therefore, we selected HEMOs #1 and #5 to study the lineage differentiation and underlying mechanisms of late stage HEMOs. In HEMO #5, Clone 1 was characterized by dominance of yolk sac endoderm (YSE) and fibroblasts (Fibro), with minimal mesoderm-like cells (Mes-like) progenitor activity (Fig. 4f, g). This suggests an early bias toward stromal and extraembryonic differentiation, potentially reflecting a niche-supporting role. Predominantly surrounding Clone 2 of HEMO #5 (Fig. 4c, d), Clone 1, with its high Fibro composition, was positioned in a supportive role within the stromal niche. Fibro likely provided structural support or secrete signals regulating neighbouring clones. The presence of pluripotent stem cells-like (PSC-like) cells hinted at retained pluripotency or self-renewal capacity, while low trophoblast-like cells (TB-like) and absent hematopoietic erythroid cells (Ery) and erythro-myeloid progenitor (EMP) lineages indicated limited trophoblast or blood-lineage priming of Clone 1 in HEMO #5.

Clone 2 marked a transitional phase, with a resurgence of Mes-like progenitors alongside a sharp rise in TB-like cells (Fig. 4f, g). Residing in the HEMO core, Clone 2 balanced Mes-like (23.1%) progenitors with TB-like (22.7%) and YSE (26.8%) cells, alongside rare hematopoietic progenitor cells (HPC) (1.95%) (Fig. 4c, d, f, g). This central niche may act as a developmental crossroads, where mesenchymal progenitors (Mes-like) transiently reactivate hematopoietic potential (HPC) while seeding trophoblast (TB-like) and yolk sac (YSE) lineages toward peripheral zones. The central position aligned with a "signaling hub" role, coordinating differentiation cues for surrounding clones in HEMO #5.

Significantly localized to the right bottom edge of HEMO #5, Clone 0’s dominance of TB-like (21.1%) and YSE (21.5%) cells, alongside residual Mes-like (15.0%) and PSC-like (11.4%) populations, suggests a role in extraembryonic boundary formation. Its peripheral position aligned with TB-like and YSE lineages, which often demarcate tissue interfaces or nutrient-exchange zones (Fig. 4c, d, f, g). The persistence of PSC-like cells at the edge hints at a "stemness reservoir" for tissue maintenance or repair at dynamic boundaries.

Similarly, HEMO #1 central clone (Clone 2) consistently occupied dynamic, transitional niches, marked by mixed Mes-like and lineage-primed populations (EMP) (Fig. S30d, e). This positions it as a differentiation hub directing lineage-restricted outputs (megakaryocyte/erythroid lineages) rather than retaining multipotent hematopoietic potential. Conversely, peripheral clones (Clone 0) were enriched in stromal (Fibro), extraembryonic (YSE and TB-like), or niche-supporting lineages (Fig. S30d, e). It lacked PSC-like cells, suggesting its niche role focused on structural support rather than pluripotency retention. In both HEMO #1 and #5, a shared hierarchy emerges: stromal/extraembryonic-biased clones (Clone 1) localize to lower/edge zones, while transitional clones (central Clone 2 in both) bridge progenitor states and lineage outputs. Notably, TB-like and YSE lineages showed spatial polarization, dominating peripheral niches in both HEMOs, likely reflecting conserved roles in boundary formation or nutrient exchange. The persistence of PSC-like cells at edges and reactivation of HPC or TB-like potential in cores further suggested a universal tension between stemness maintenance and lineage commitment across spatially partitioned niches. These patterns highlighted a recurring logic: spatial segregation of clones into stemness-permissive edges and differentiation-active cores, orchestrated by microenvironmental cues.

### NOTCH signaling orchestrates spatial lineage transitions via stromal-progenitor crosstalk in HEMOs

To elucidate the molecular mechanismsIG underlying this spatial lineage transition during organoid development, we conducted ligand-receptor (LP) interaction analysis in the aforementioned targeted lineages in both HEMO #1 and #5 (Fig. 5, and Fig. S31). CellPhoneDB revealed a high interaction score for NOTCH ligands and receptors between Fibro and PSC-like, as well as Fibro and YSE in Clone 1 of HEMO #5 (Fig. 5a). The high interaction score was also discovered between Mes-like and EMP, as well as Fibro and HPC in Clone 2 of HEMO #5 (Fig. 5b). Intrigued by the critical role of NOTCH signaling in developmental hematopoiesis (Ditadi et al., 2015; Hadland et al., 2015; Koch et al., 2013), we quantified both the activation and inhibition of NOTCH signaling based on the combination of ligand-receptor interactions (Fig. 5c, d). In HEMO #5, PSC-like and YSE expressed Delta-Like Non-Canonical Notch Ligand 1 (DLK1), while Fibro expressed Notch Receptor 1 (NOTCH1) in the same spatial area within Clone 1 (Fig. 5c, d). Similarly, EMP and HPC expressed DLK1, while Fibro and Mes-lie expressed NOTCH1 in the same spatial region of Clone 2 (Fig. 5c, d). DLK1 regulates the hematopoietic progenitor pool by suppressing proliferation and differentiation (Li et al., 2005; Mirshekar-Syahkal et al., 2013; Qian et al., 2016). Immunofluorescence staining confirmed the spatial organization of this niche (Fig. 5e), highlighting the pivotal role of NOTCH signaling in both clonal architecture and niche regulation within human embryonic organoid models.

## Discussion

Our study establishes mitochondrial DNA (mtDNA) variants as endogenous genetic recorders that enable high-resolution reconstruction of spatiotemporal clonal dynamics during hematopoietic organoid (HEMO) development. By integrating mtDNA-based lineage tracing with single-cell and spatial transcriptomic profiling, we uncover fundamental principles of cell fate decisions and tissue self-organization that reflect conserved developmental processes reminiscent of early human embryogenesis. These findings significantly advance our understanding of in vitro hematopoiesis and provide a conceptual and technical framework for investigating lineage segregation and clonal architecture in complex cellular ecosystems.

A major insight arising from our analysis is the striking spatial zonation of clonal units within HEMOs. We found that clones are not randomly distributed but occupy distinct microniches: peripheral regions were enriched in pluripotent (PSC-like) and extraembryonic (TB-like, YSE) lineages, whereas central areas harbored more mesenchymal (Mes-like) and hematopoietic (EMP, HPC) progenitors. This structural organization—which we term “stemness-permissive edges and differentiation-active cores”—parallels spatial patterning phenomena observed in vertebrate embryos and mirrors mechanisms observed in early embryogenesis (Tyser et al., 2021) suggesting the influence of conserved biophysical or biochemical gradients on cell fate specification. The clear correlation between clonal identity and spatial context underscores that organoid development is not solely cell-intrinsic but is profoundly shaped by positional cues, potentially mediated through morphogen gradients, cell-cell contact, and metabolic microenvironments.

Notably, ligand-receptor analysis provided mechanistic insight into this spatial patterning by revealing enriched NOTCH signaling between stromal (Fibro, Mes-like) and progenitor (PSC-like, HPC, EMP) cell populations within specific spatial niches. This finding resonates with the essential role of NOTCH signaling during in vivo embryonic hematopoiesis, particularly in the aorta-gonad-mesonephros (AGM) region and fetal liver. It aligns with in vivo studies showing that NOTCH ligands in the bone marrow niche regulate hematopoietic stem cell maintenance (Espinosa & Bigas, 2011), and the observed NOTCH1 signaling suggest a model in which fine-tuned stromal-progenitor crosstalk regulates hematopoietic differentiation in a spatially restricted manner, further validating HEMOs as physiologically relevant models of human hematopoietic development.

Our comparative analysis indicates that HEMOs recapitulate many features of yolk sac and early AGM-stage hematopoiesis, including the emergence of erythro-myeloid progenitors and endothelial commitment. That said, the absence of robust lymphoid differentiation and inter-organoid variability highlight remaining limitations. Nevertheless, the faithful reconstruction of NOTCH-mediated signaling and the spatial organization of clonal structure reinforce the physiological relevance of this model for studying human hematopoietic development.

An important methodological contribution of our work is the demonstration that mtDNA variants can significantly improve confidence in lineage tracing. Recent studies have pioneered the use of mtDNA variants to track clonal dynamics in contexts such as aging hematopoiesis (Lareau et al., 2021) and cancer evolution (Ludwig et al., 2019). However, our work is the first to apply and validate this approach for high-resolution lineage tracing in a developing human organoid model. We found that nearly 60% of putative multi-lineage clones identified by the LARRY barcoding system were actually polyclonal aggregates—a result of barcode homoplasy rather than true multipotency. The use of mtDNA variants enabled us to distinguish such technical artifacts from legitimate multipotent progenitors, including clones capable of producing both ectodermal and trophoblastic lineages, a fate combination seldom reported in conventional differentiation systems. This orthogonal approach greatly enhances the accuracy of lineage reconstruction in densely populated organoid systems.

However, clonal fate assignments remain inferential, based on transcriptomic states; functional validation through differentiation assays or transplantation experiments will be necessary to firmly establish potency. Looking forward, the multimodal framework developed here offers multiple promising avenues. Perturbation experiments targeting NOTCH signaling—via small molecules or genetic editing—could functionally validate its role in spatial fate patterning. Applying this platform to disease-specific iPSCs could elucidate clonal dynamics in genetic blood disorders or leukemias. Technologically, integration of mtDNA lineage tracing with CRISPR-based recording systems could further enhance resolution for building comprehensive lineage maps.

In conclusion, our work establishes mtDNA variants as powerful natural barcodes for reconstructing lineage relationships and spatial clonal dynamics in human organoids. Beyond providing technical corrections to synthetic barcoding systems, we demonstrate that clonal architecture is intimately linked to spatial niche identity, unveiling patterning principles that underlie early blood development. This integrated approach offers a new foundation for studying human development, disease modeling, and regenerative medicine with unprecedented spatial and clonal resolution.

## Methods

### HEMO differentiation

HEMO differentiation was done based on the previous established protocol (Chao et al., 2023). In brief, we formed EB from hPSCs and differentiated to mesoderm and hematoendothelial lineage by morphogens and cytokines.

#### hPSCs line

Human M1-hPSCs were gifted by Pentao Liu and maintained in the media described before (Gao et al., 2019).

#### hPSCs maintenance and pre-differentiation

The hPSCs were cultured and maintained in PSC medium, with medium changes occurring every other day. The composition of the medium was previously reported (Gao et al., 2019). Prior to the formation of embryoid bodies (EBs), cells were pre-differentiated in KSR medium (DMEM/F12 + 10% KSR) (Thermo Scientific, catalog no. 11320033; Thermo Scientific, catalog no. 10828028) for a period of three days.

#### EB formation in hanging drop

To begin the process, the KSR medium was removed and the cells were washed with PBS (Thermo Scientific, catalog no. 10010023). The cells were then digested with 500 mL of 0.05% Trypsin (Thermo Scientific, catalog no. 25300054) at 37°C for three minutes, but no longer than seven minutes. The Trypsin was removed and 2 mL of KSR medium was added to harvest the cells. Careful handling of the cells was necessary, and harsh pipetting was to be avoided. The cells were then centrifuged at 300g for three minutes, and the supernatant was carefully removed. To resuspend the cells, 1 mL of KSR medium and 1 μL of Y27632 (10 ug/ml) (Tocris, catalog no. 1254/10) were added. For each 25 μL hanging drop on the cap of a 10 cm petri dish, 4,000 cells were added. Thirty to forty drops were made for each cap, and the dish was filled with PBS to keep it moist. The cap was then gently and slightly inverted to cover the dish, and the dishes were labeled and kept at 37°C for three days.

#### EB collection and differentiation

After the formation of embryoid bodies (EBs), all EBs were collected and washed with PBS. The EBs were then centrifuged at 100g for one minute, and the supernatant was carefully removed. 1 mL of medium A (STEMdiff Hematopoietic Kit, catalog no. 05310) was added, and the EBs were transferred to a non-adherent 24-well plate. The date was labeled as Day 0, and the EBs were cultured with 1 mL of STEMdiff medium A for three days. On Day 2, the medium was half-changed, and the EBs were subsequently cultured with 1 mL of STEMdiff medium B for the following days. This process involved the maintenance of hPSCs, pre-differentiation, EB formation, collection, and differentiation.

### Chondrosarcoma sample

The current study was approved by the Institutional Review Board of the University of Hong Kong/Hospital Authority Hong Kong West Cluster (IRB reference number: UW 16-2036). The patient sample was obtained from Queen Mary Hospital, Hong Kong, following informed consent from the patient. The fresh specimen was collected during surgical resection and immediately transported on ice to maintain tissue integrity. Diagnosis and grading were performed by a multidisciplinary team of pathologists and orthopaedic surgeons. The specimen was stored at -80°C prior to processing.

### 10x Genomics Chromium scRNA-seq

To prepare single-cell suspensions, organoids were harvested at D8, D15, D18, D25 and D32 since EB formation in hanging drop. After washing with PBS, organoids were mechanical chopped with scissors for 20-30 times, followed by digestion with 500 μL Accumax at 37 °C for 10-15 minutes. Digestion was terminated by 500 μL PBS containing 2% FBS. Cells were filtered with 40 μm cell filter and centrifuged at 500 × g for 5 min. The collected cells were resuspended in FACS sorting buffer (1 × PBS with 2% FBS) and their concentration was adjusted to around 3k cells per μL by counting with a hemocytometer.

Subsequently, cells were stained with DAPI (BD Biosciences, catalog no. 564907) in 1:100 for 5 min at 4 °C and were sorted by BD Influx flow cytometry in CPOS at HKUMed. Cells were gated to exclude dead cells and doublets and collected in a chilled PBS with 0.04% BSA for scRNA-seq library construction. Cell concentration was adjusted to around 500-1000 cells per μL.

10x Genomics 3′ Single Cell Gene Expression v3 reagents were used for scRNA-seq library preparation. Single cell encapsulation and reverse transcription were performed at the Centre for PanorOmic Sciences, the University of Hong Kong. Briefly, 8-16k live single cells of size 30 μm or smaller and of good viability were encapsulated in gel bead-in emulsions. Captured mRNAs were transcribed into cDNAs and amplified to generate whole-transcriptome amplifications (WTAs). Libraries were constructed by fragmentation, adapter ligation and a sample index PCR. Libraries were sequenced using Illumina Novaseq 6000 for Pair-End 151bp sequencing.

### 10x Genomics Visium library construction and sequencing

The Visium Spatial Tissue Optimization Slide & Reagent kit (10x Genomics, catalog no. PN-1000193) was used to optimize permeabilization conditions for the tissue sections. The Visium Spatial Gene Expression Slide & Reagent kit (10X Genomics, catalog no. PN-1000184) was used to generate spatially barcoded cDNA from every 10-µm sections of D15 HEMO samples. Library construction kit (10 x Genomics, catalog no. PN-1000190) was used for library construction. Frozen HEMO sample preparation, RNA integrity detection, tissue optimization, staining and imaging, cDNA synthesis and second strand synthesis, cDNA amplification and library construction were carried out according to Chao et al. (Chao et al., 2023). Libraries were sequenced on Illumina sequencing instruments using TruSeq Read 1 and TruSeq Read 2 primers.

### Mitochondrial alteration enrichment for Chromium-MAESTER and Visium-MAESTER

MAESTER was performed according to Miller et al. (Miller et al., 2022). Briefly, WTAs yield in high-throughput 10x Genomics scRNA-seq 3’ and Visium platforms served as initial templates for the enrichment of 15 mitochondrial transcripts by two rounds of PCR. Sequences of the amplified mitochondrial transcripts were obtained by standard next-generation sequencing.

In PCR1, twelve primer mixes were created using the designed primers tiled across the entire mitochondrial transcriptome (Table 1). A barcoded sample indexing i5 primer was included in each mix in which 20 ng WTAs were included. PCR was performed using the following conditions: denaturation at 95 ℃ for 3 minutes, followed by 6 cycles of 98 ℃ for 20 seconds, 65 ℃ for 15 seconds, and 72 ℃ for 3 minutes, ending with a final extension at 72 ℃ for 5 minutes. Following amplification, the PCR products were pooled in a certain ratio and purified with 1x AMPure XP beads (Beckman Coulter A63881).

**Table 1.**
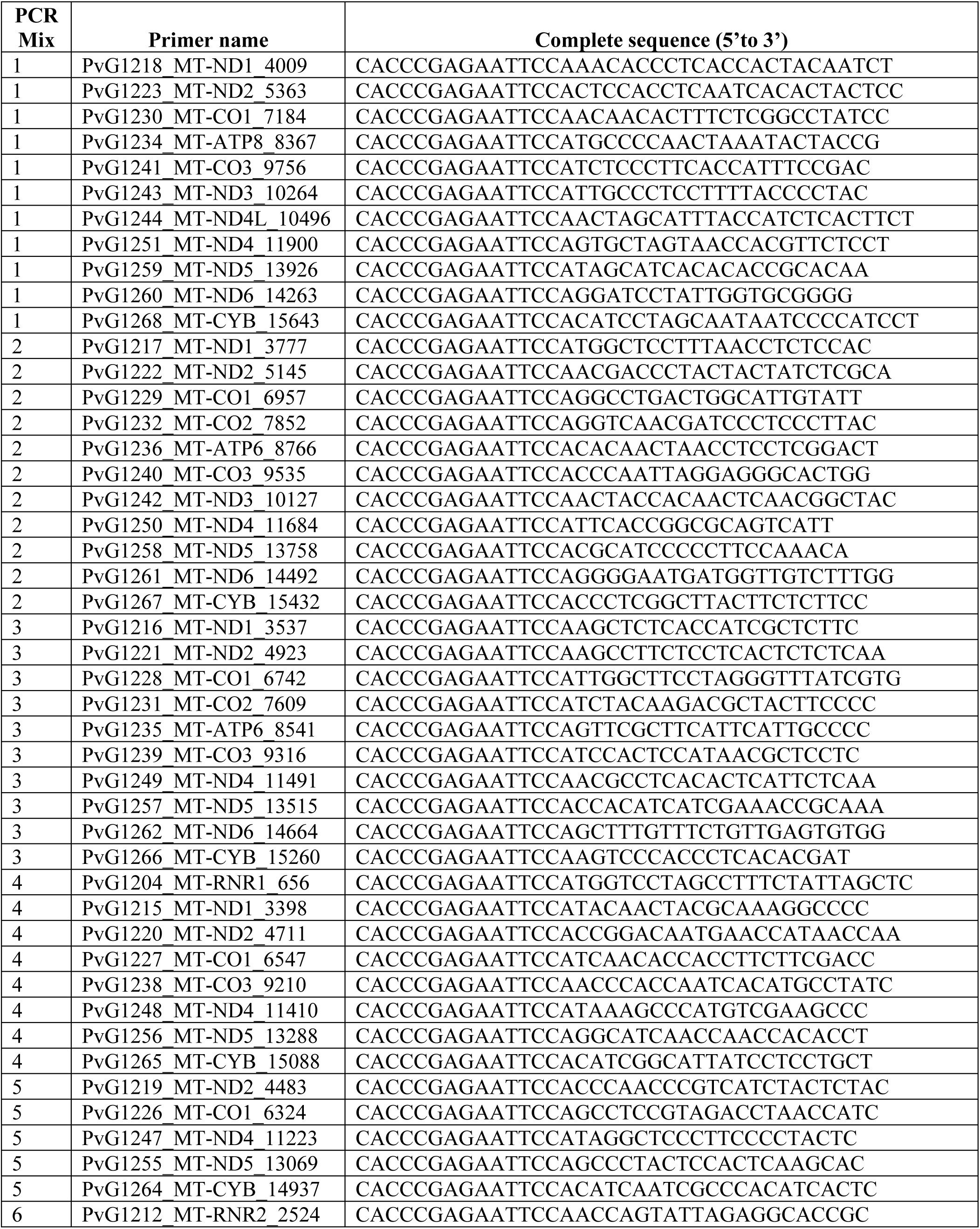
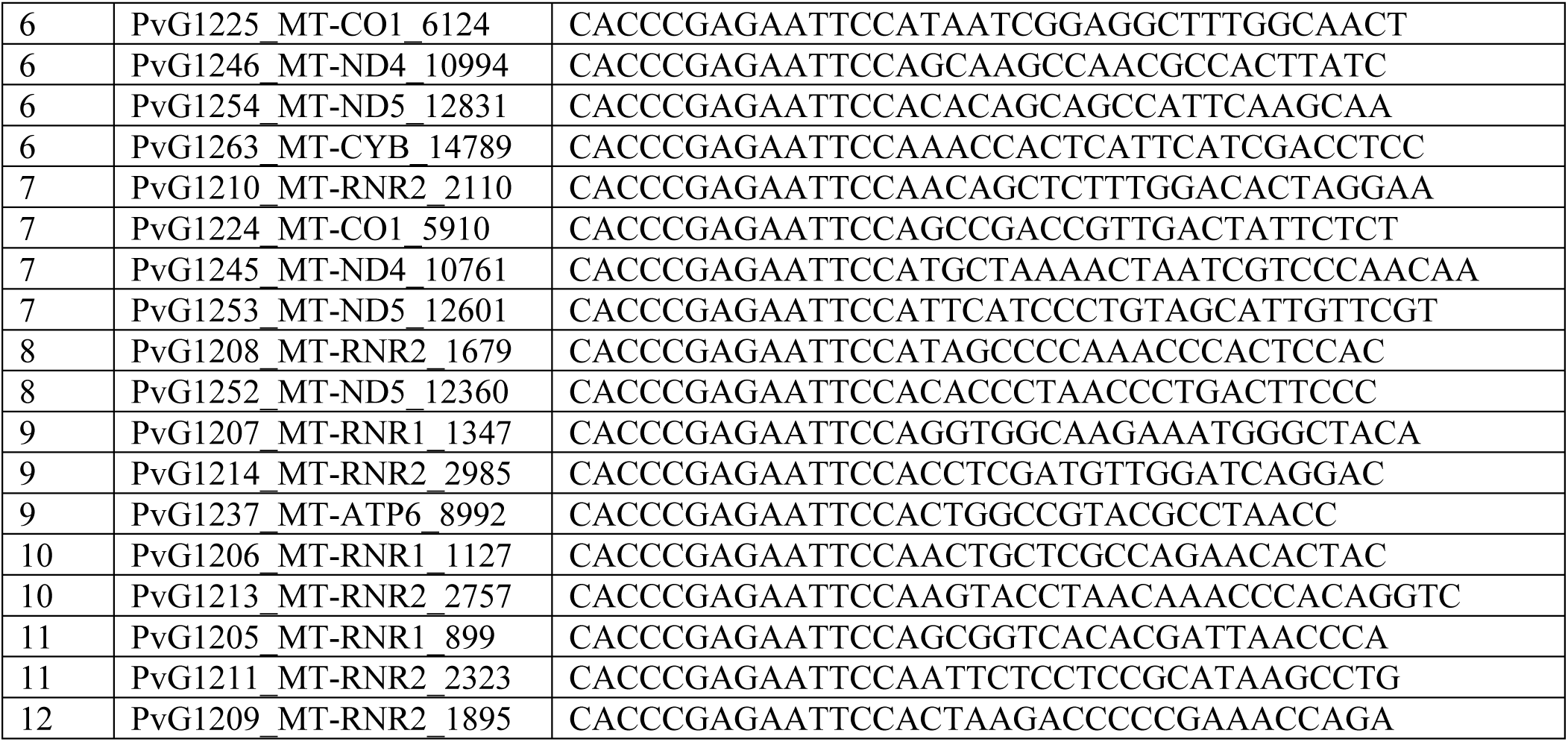
Primers used for mitochondrial variants enrichment.

The purified products were used as the template for PCR2 in which the Illumina adapters (P5, P7), dual index barcodes to identify the sample (i5, i7), and sequencing primer binding sites to the fragments were added. The programme of PCR2 was an initial denaturation at 95 ℃ for 3 minutes, then 6 cycles of 98 ℃ for 20 seconds, 60 ℃ for 30 seconds, and 72 ℃ for 3 minutes, and then a final extension at 72 ℃ for 5 minutes. After PCR2, the DNA was purified with 0.8x AMPure XP beads. The library concentration of three Chromium- and Visium-MAESTER samples were 50, 59, 71 and 21 ng/mL respectively. Resulting MAESTER and Visium-MAESTER libraries were sequenced on the Illumina NovaSeq SP PE150 kit with run cycle 28, 8, 8, 256 for Read 1, i7, i5 and Read 2 respectively. The sequences of Nove seq primers are listed in Table 2.

**Table 2:**
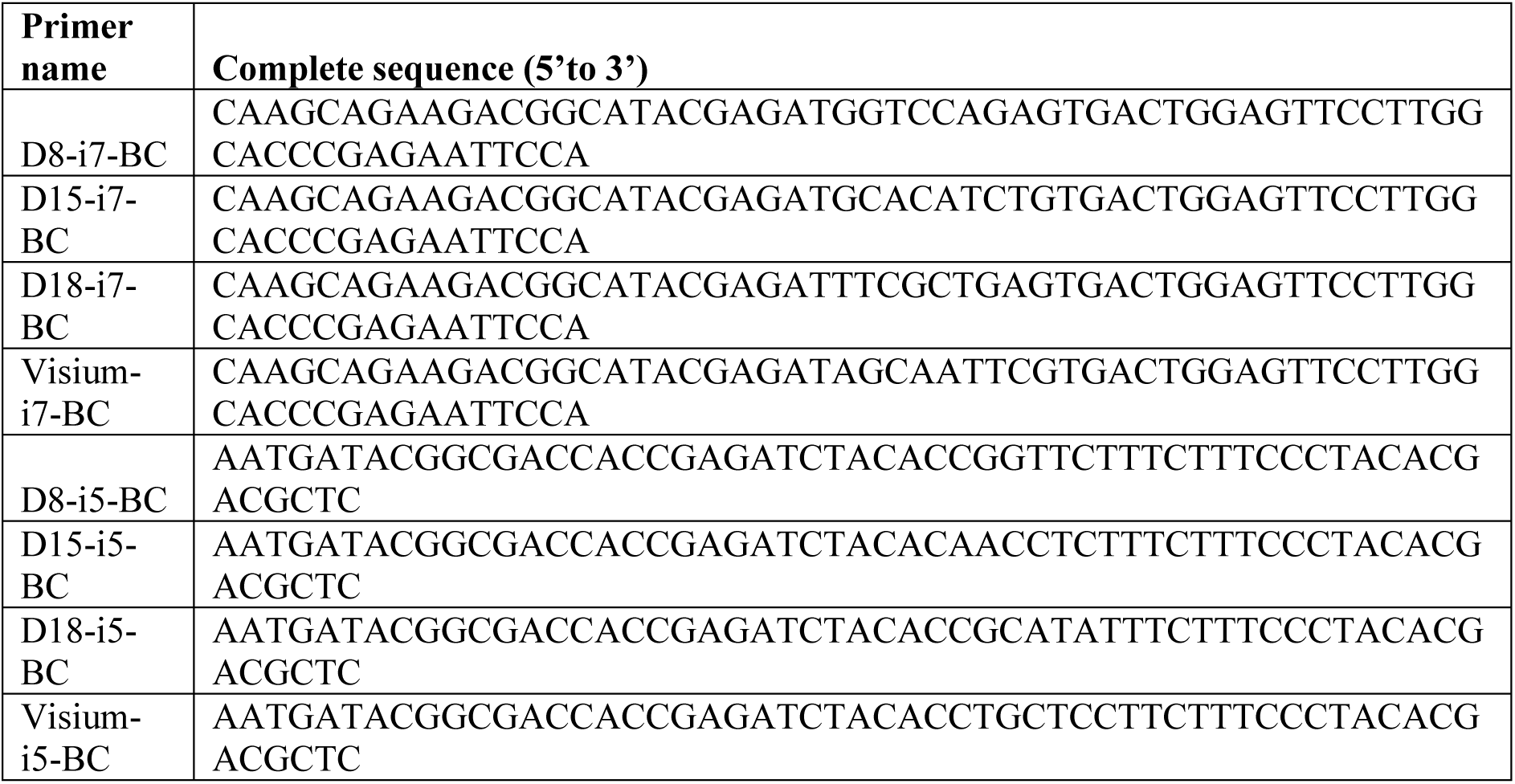
Primers used for Illumina NovaSeq.

### Immunofluorescence staining and imaging

D15 HEMO samples were paraffin embedded. The subsequent fixation, sectioning, dehydration, permeabilization and antibody incubation were performed according to Chao et al., 2023 (Chao et al., 2023). Following the washing step with PBS twice, the slides were maintained in a wet state. Subsequently, images were captured using a Nikon Ti2E Fluorescent microscope and merged using ImageJ software. Antibodies for hematopoietic cells were anti-CD45 antibody (Invitrogen catalog no. MA5-17687) and Goat Anti Rabbit IgG H&L (Abcam catalog no. ab150157). Antibodies for DLK1 were anti-DLK-1 antibody (Abcam catalog no. ab89908) and Goat Anti-Mouse IgG H&L (Abcam catalog no. ab175473). Antibodies for Notch1 were anti-Notch1 antibody (Abcam catalog no. ab245686) and Goat Anti-Rabbit IgG H&L (Abcam catalog no. ab150079).

### Data analysis and statistics

#### scRNA-seq data processing and analysis

We annotated the transcriptomic data using the Scanpy package (v1.10.2). Quality control (QC) was applied to filter out cells expressing fewer than 100 genes and genes expressed in fewer than 3 cells. Doublet detection was performed using Scrublet (Wolock et al., 2019), and identified doublets were removed. Next, we selected the top 2000 highly variable genes for dimensionality reduction. These genes were used to compute the neighborhood graph of cells, which was further visualized using UMAP (McInnes et al., 2020). Finally, we identified cell types such as ’Mes-like’, ’TB-like’, ’Fibro’, ’PSC-Ect’, ’PSC-like’, ’EC’, ’Mes-PS’, and ’Endoderm’ based on known marker genes. The identified cell types were visualized in UMAP.

#### RNA velocity analysis

To investigate dynamic transcriptional states and infer potential cell fate transitions, we performed RNA velocity analysis using the scVelo (v0.2.4) and cell2fate (v0.1a0) (Aivazidis et al., 2025)packages. Spliced and unspliced count matrices were obtained from loom files processed by velocyto (v0.17.15) and integrated with cell-type annotated transcriptomic data using scv.utils.merge(). We applied the Cell2fate_DynamicalModel to reconstruct transcriptional dynamics, automatically determining the number of modules based on data structure. The model was trained using a GPU-accelerated PyTorch backend with default parameters over multiple iterations until convergence. Velocity-inferred latent time and module activation states were projected onto the UMAP embedding to visualize developmental trajectories.

We further identified gene modules associated with specific transcriptional programs and computed module-specific velocity vectors. Genes within each module were ranked by their contribution weights, defined as the proportion of total inferred expression attributable to the module. Transcription factors (TFs) were specifically identified by overlapping the ranked genes with a curated list of human TFs provided in the Cell2fate package. Top features, including these identified TFs, were selected based on this ranking. Gene expression values visualized in UMAP plots were min-max normalized to a [0, 1] scale to ensure consistent interpretation. Visualizations of velocity streamlines, module activation states, and key gene expression patterns were generated using customized plotting functions based on scVelo and Scanpy.

#### 10x Visium spatial transcriptomics processing

The sequencing data and imaging tiff files of individual 10x Visium slides were processed using SpaceRanger software (version 1.3.1) with the GRCh38 human reference genome. Count matrices were then loaded into the Scanpy package (version 4.1.0) in Python. Spots were selected based on RNA counts and mitochondrial gene percentage. After spot selection, high variable genes (HVGs) were identified and used for principal component analysis (PCA) to reduce dimensionality and perform clustering. The Scanpy adata objects were utilized for downstream analysis individually.

#### Spatial transcriptomics spot deconvolution

Two packages were utilized for spatial transcriptomics spot deconvolution. The first method applied was SpatialScope, a newly developed statistical method that integrates scRNA-seq and spatial transcriptomics data to obtain the spatial distribution of the whole transcriptome at the single-cell resolution. SpatialScope initially performs nuclei segmentation to count and locate nuclei within each spatial spot. It then assigns a cell type to each located nucleus. Finally, by leveraging the paired scRNA-seq reference data and a learned deep generative model, it decomposes gene expression at each spot into gene expression of the individual cells located within the spot. The RCTD package (version 2.0.0) was also utilized to validate the cell proportion (Cable et al., 2022). It first takes the single-cell reference and predicts the proportion of each cell type in an individual 10x Visium spot.

#### Cell interaction within spatial transcriptomics data

Two different packages were utilized for spatial cell-cell interaction analysis. After obtaining single-cell resolution spatial data, CellPhoneDB within the Squidpy toolbox was utilized to predict cellular ligand-receptor interactions (Efremova et al., 2020; Palla, Spitzer, et al., 2022). All predicted pairs were pre-selected based on a p-value of 0.001. The enriched score and metadata of individual pairs were then examined, and the final results were presented in a bar plot.

#### Mitochondrial variant calling

In the MQuad pipeline, data pre-processing was performed before mitochondrial variant calling. After getting the fastq files of MAESTER and Visium-MAESTER data, umitools were first used to generate a whitelist of valid barcodes. The number of cells was estimated by the data. Based on the whitelist of valid barcodes, umitools extracted reads coming from real cells and moved UMIs and barcodes from sequence lines to the headlines. In this step, background reads and barcodes with low number of reads were filtered. Reads were aligned to the reference genome by STAR using hg38 as the reference genome. UMIs and barcodes were added as tags in the bam file by using pysam. Mitochondrial variants were called by cellsnp-lite mode 2a. We used minimum minor allele frequency 0.1 and minimum aggregated count as filtering criterion. MQuad (Kwok et al., 2022) was applied to call informative mitochondrial variants for the following analysis.

For maegatk pipeline, pre-processing of the raw sequence files is required before maegatk processing. The maegatk pipeline was promoted as being superior to the previous mgatk tool due to its capability in performing UMI consensus using the additional ‘-mr’ parameter flag in filtering out reads with less than the minimally specified amount of reads. This is speculated as an important aspect in pre-filtering mtDNA reads to improve the subsequent variant calling results. However, when comparing the output generated using UMI consensus and without, the distinction is unclear (Fig. S5a). The number of variants identified using the same threshold varies for each dataset and does not suggest higher sensitivity after applying a UMI consensus limit of 3 when running maegatk. Yet, there is insufficient data from this study to conclusively dismiss the advantages of the added UMI parameter in maegatk processing, thus further analysis is required to determine the importance of this added parameter when considering using maegatk for MAESTER mtDNA sequencing data processing. Using the maegatk output generated with the suggested minimum 3 UMI reads parameter, cells with more than 1% VAF for any identified variant are selected and subset for the subsequent clonal assignment step.

#### Clonality reconstruction

In the MQuad pipeline, clone assignment was performed using vireoSNP (Huang et al., 2019). First, we determined several candidate optimal clone numbers by analyzing an elbow plot, following the guidelines from the ‘Mito Clones’ tutorial. Based on this tutorial, we then generated assignment probability plots and mean allele ratio plots, starting with the lowest number of candidate clones. The optimal clone number was selected to maximize clonal resolution while ensuring high-confidence assignments. Specifically, we retained as many clones as possible, provided that each clone exhibited unique patterns of allele frequency in terms of allele ratios. Additionally, we ensured that each clone had a high assignment probability for its constituent cells. Finally, we used the anno_heat function from vireoSNP to visualize the allele frequency heatmap for each cell.

In the maegatk pipeline, the clValid R package (Brock et al., 2008) was employed to determine the optimal number of clusters and clustering method based on the variant allele frequency matrix. This package provides statistical validation for clustering and was used here to evaluate various combinations of cluster numbers and clustering methods. Internal validation metrics— such as silhouette width, Dunn index, and connectivity scores—were used to assess and visualize the quality of clonal clustering iterations. For all datasets, hierarchical clustering was identified as the optimal method, with varying numbers of clusters selected based on the validation results. In the next step, hierarchical clustering was performed using the eclust function in R, with Euclidean distance as the metric and the Ward.D method to minimize inter-cluster variance.

#### LARRY lineage tracing analysis

LARRY barcode-based lineage tracing was performed to find the clonal relationships that underpin cellular fate determination. The analytical workflow comprised the following steps: (1) Preprocessing of raw data to eliminate cells with invalid LARRY barcodes, by flowing the quality control framework established by Weinreb et al. (Weinreb et al., 2020); (2) Assignment of clones through the aggregation of barcodes utilizing similarity matrices; (3) Further filtration of LARRY clones containing only a single cell for downstream analysis; (4) Integration with single-cell RNA sequencing (scRNA-seq) annotations via the Cospar package (v0.3.1) to establish correlations in lineage fate (Wang et al., 2022). A statistical evaluation of clonal biases across cell types was conducted using Fisher’s exact test with Benjamini-Hochberg correction, implemented within the Cospar computational framework.

#### MAESTER dataset processing pipeline

The MAESTER dataset analysis involved a comprehensive four-module processing pipeline: (1) Read alignment was performed using Cell Ranger (10x Genomics) with the GRCh38 human reference genome; (2) Variant calling was conducted through cellSNP-lite (v1.2.0) utilizing parameters (minimum allele frequency [MAF] = 0, minimum count = 1) to enhance variant retention; (3) Informative variant selection was executed via MQuad (v0.1.3) with default thresholds; (4) Clone assignment was implemented using vireoSNP (v2.1.4), which determined optimal clone numbers through elbow plot analysis of clustering stability metrics. Cells lacking variants were filtered out, specifically those without any variants with coverage >=2 and an alternative allele. Candidate variants are further filtered for variants with allele frequency >0.05 across 95% of cells and less than 5% of cells.

#### Mapping mtDNA variants into spatial space

The integration of mtDNA variants with spatial information enables the identification of spatial distribution of mtDNA within the sample. Initially, we map mtDNA variant information into spatial contexts by identifying spatial coordinates using cell barcodes. Following integration, the mtDNA clone information and single nucleotide polymorphism (SNP) allele frequencies for each cell are visualized within the spatial framework.

To further analyze mtDNA data, we employ SpatialDE (version 1.1.3), a tool originally designed to identify genes significantly dependent on spatial coordinates. The allele frequencies of mtDNA are normalized to a scale of 100, after which SpatialDE is applied to identify spatially variable mtDNA features. Features with an adjusted Q-value of less than 0.05 are considered significant.

#### Integration of mtDNA variants and spatial transcriptomics data

Spatially resolved transcriptomic data were obtained from our previous study (Chao et al., 2023). We selected five HEMOs representing diverse cell types within each orgnoid. For each hemo cell, we gathered single-cell resolution annotations generated through SpatialScope’s deconvolution algorithm. The mtDNA variants for each spot were processed using our MAESTER analysis pipeline. We integrated the mtDNA information for each spot with the cell type percentages to investigate (1) the differences in cell types associated with various mtDNA SNPs and (2) the variations in cell types among different mtDNA clones. Upon identifying distinct cell type distributions for different clones, we further analyzed the cell-cell interactions within a selected clone. The cell-cell interaction analysis was performed using CellPhoneDB’s permutation testing methodology (Efremova et al., 2020), involving 1,000 permutations and retaining interactions that met dual thresholds: statistical significance (adjusted p-value < 0.01) and biological relevance (mean expression score > 6.5). Directionality assignment for inhibitory interactions was derived from Squidpy’s module (v1.6.2), where inhibitory relationships were annotated with negative interaction weights.

## Code availability

The code for analysis is available at https://github.com/sujunhao/HEMO_maester_spatial

## Data availability

The data for analysis is available at https://www.ncbi.nlm.nih.gov/bioproject/PRJNA855311

## Acknowledgement

We appreciate Dr Peter van Galen at Harvard Medical School for generous sharing of bioanalyzer positive library controls, critical discussion on this project, and comments to the manuscript. We thank CPOS at HKUMed for technical assistance in 10x Chromium, 10x Visium, and Novaseq. This study is supported in part by ECS 27109921 (R.S.) and GRF 17109022 (R.S.) by the Hong Kong Research Grant Council, and the Shenzhen-Hong Kong-Macau Technology Research Programme (Type C; SGDX2021082310356025 to J.W.K.H.). It is also supported in part by the InnoHK initiative of the Innovation and Technology Commission of the Hong Kong Special Administrative Region Government (AIR@InnoHK, Laboratory of Data Discovery for Health for J.W.K.H) and Health@InnoHK, Centre for Translational Stem Cell Biology for R.S.).

## Author contributions

Y. X., J. S., Y. C., J. W.K. H., and R. S. conceived the study. Y. X., L. L., Y. X., T. A., and J. W. conducted experiments and generated the data. J. S. and Y. C. conducted most informatics analysis with the guidance of Y. H. and J. W.K. H.. Y. X., X. L., M. K. H., Z. S., J. C., Z. L., C. L., R. L., and K. S. C.C. participated in informatics analysis. Y. X., J. S., Y. C., J. W.K. H., and R. S. wrote the manuscript. All authors edited the manuscript.

