## Supplementary figures for "Single-cell mitochondrial lineage tracing decodes fate decision and spatial clonal architecture in human hematopoietic organoids"

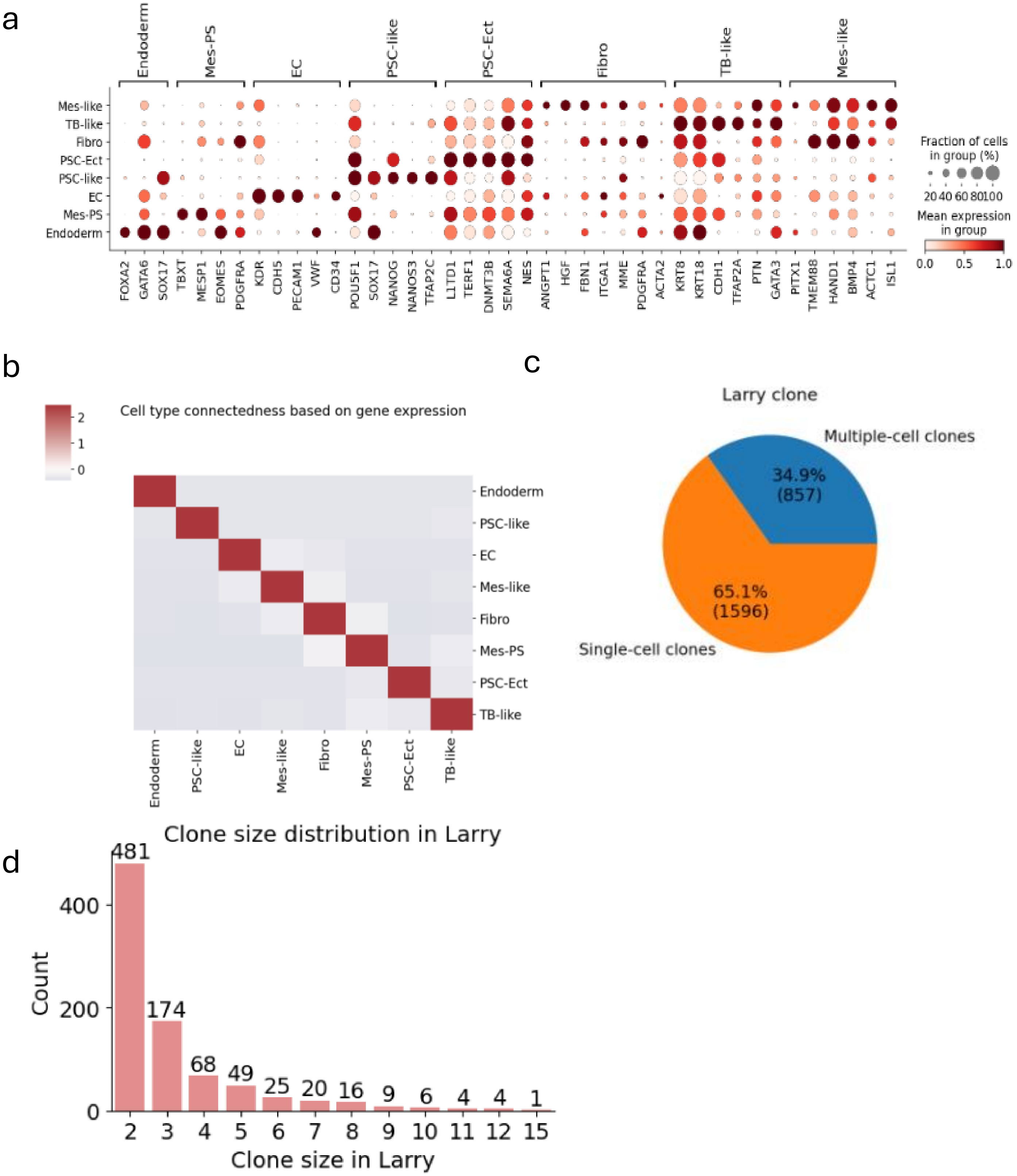


**Fig. S1 LARRY clones in early stage HEMOs (D4).**

a, Marker gene expression of D4 HEMO cell populations.

b, Heatmap showing strong intra-cell-type connectedness for each cell type.

c, Pie chart illustrating that 65% of LARRY clones are single-cell clones.

d, Bar chart showing the size distribution of non-singleton clones.

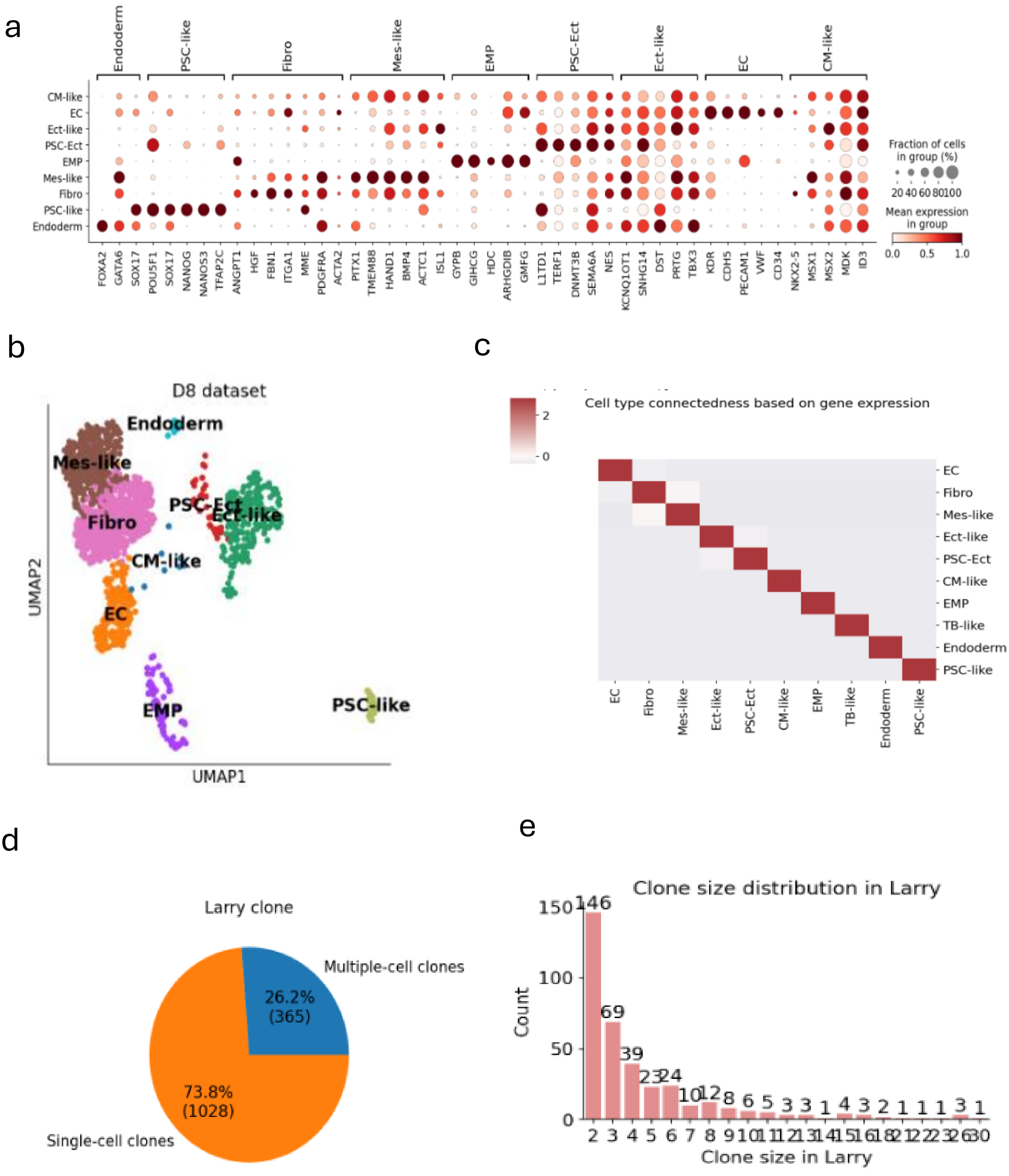


**Fig. S2 LARRY clones in late stage HEMOs (D8).**

a, Marker gene expression of D8 HEMO cell populations.

b, UMAP visualization reveals the emergence of erythro-myeloid progenitor (EMP) and cardiac mesoderm-like cells (CM-like) during late-stage HEMO hematopoiesis, highlighting lineage diversification and the dynamic progression of hematopoietic development.

c, Heatmap showing strong intra-cell-type connectedness for each cell type.

d, Pie chart illustrating that 74% of LARRY clones are single-cell clones.

e, Bar chart showing the size distribution of non-singleton clones, revealing a subset of larger clones with expanded populations.

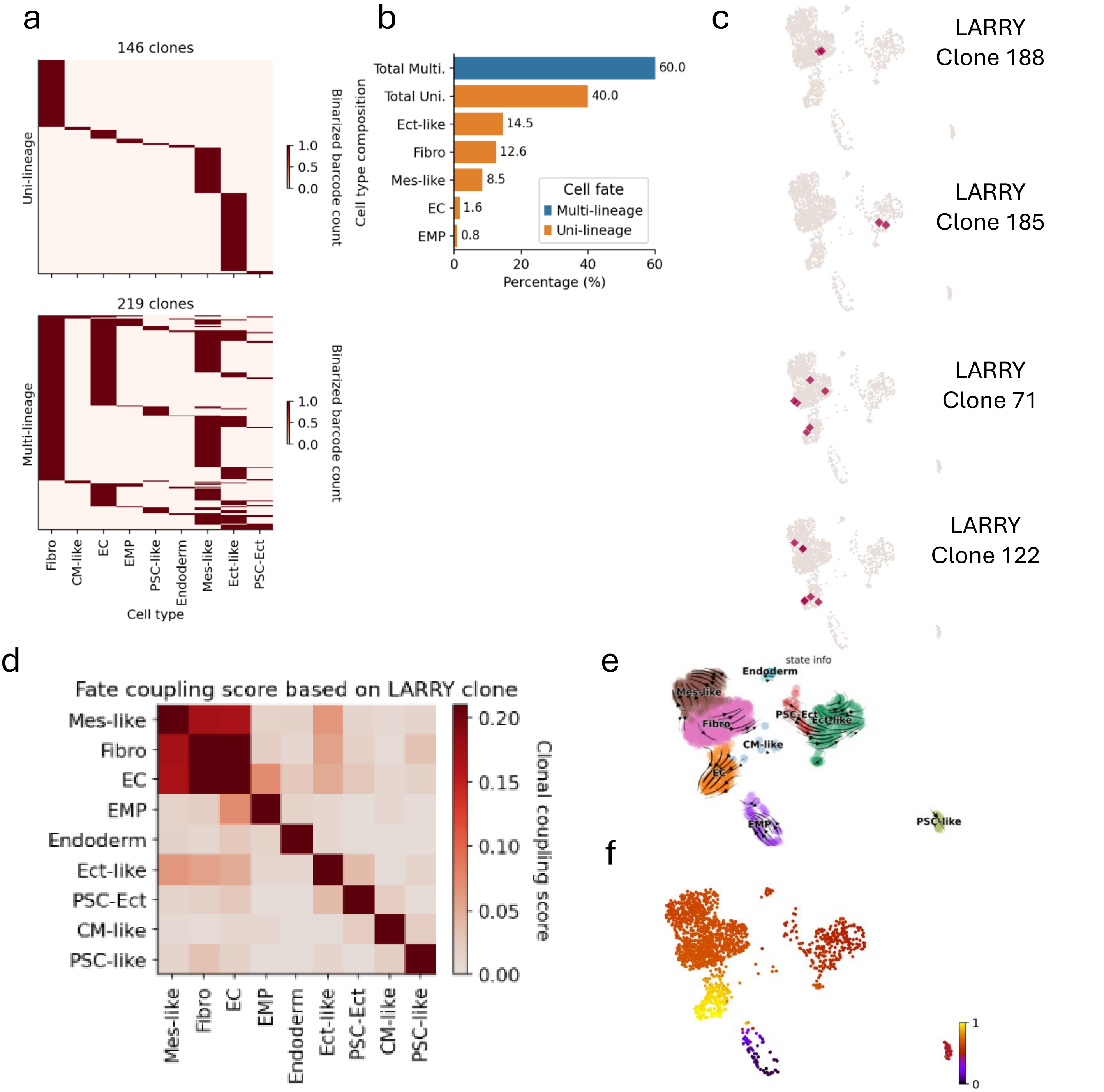


**Fig. S3 Lineage tracing reveals distinct cell fates during the late development of HEMOs and barcode homoplasy of LARRY (D8).**

a, Heatmaps showing clonal fate predictions generated by CoSpar. Each row represents a LARRY clone, and each column corresponds to a specific lineage.

b, Bar chart showing the proportion of clones in each lineage.

c, UMAP plots showing the example clones with uni- (top two) or multi-lineage (bottom two) characteristics.

d, Cell fate coupling revealed by CoSpar. The heatmap highlights significant coupling between Fibro and EC, as well as between TB-like and Mes-like cells, as identified by CoSpar analysis.

e, RNA velocity field describes the fate decisions of major HEMO lineages in LARRY dataset. The velocity field is projected onto a PCA plot with arrows indicating the local average velocity evaluated on a regular grid. RNA velocity was estimated without cell or gene pooling. the Mes-like population had largely shifted away from its early EC-biased differentiation, with only a small subset retaining fibroblast potential. PSC-Ect lineages exhibited a clear trajectory toward ectodermal differentiation.

f, Pseudotime analysis reveals that EC cells emerge at the latest pseudotime stages, reflecting their role as terminally differentiated endothelial cells.

**a**

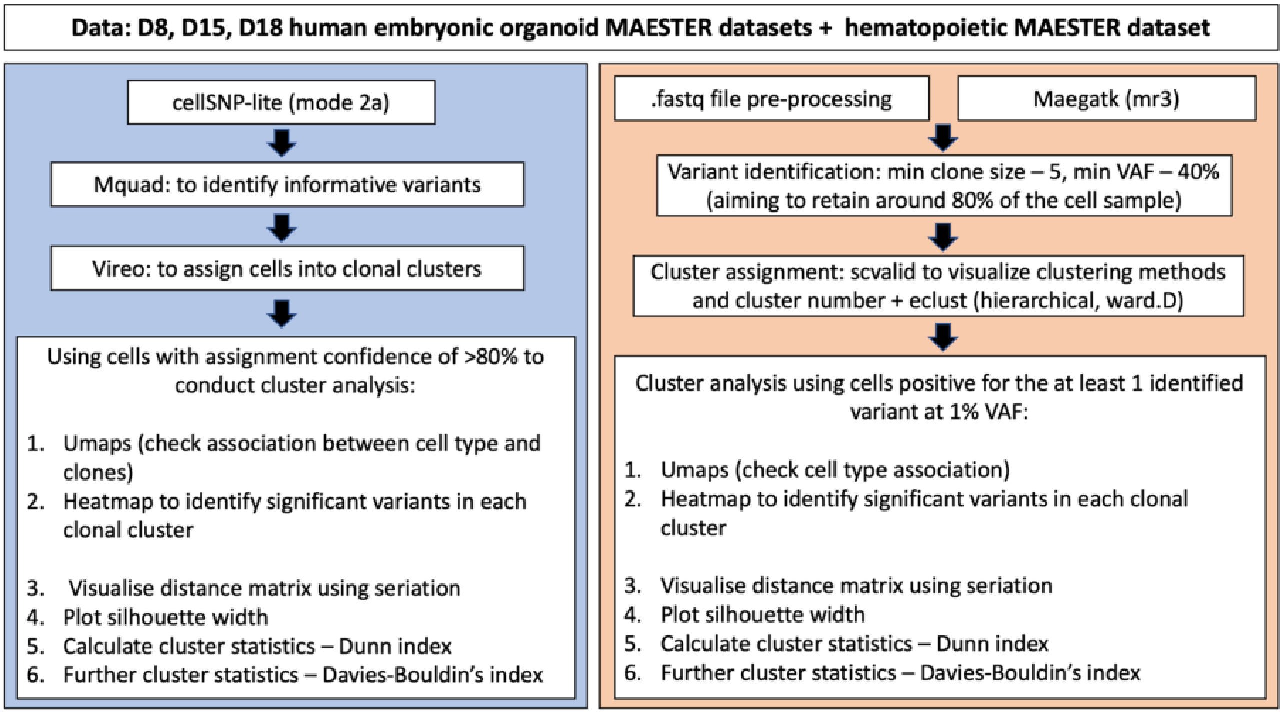


**b**

| | Human embryonic organoid datasets | | | Hematopoietic dataset |
|---|---|---|---|---|
| | **D8** | **D15** | **D18** | |
| **cellSNP** | | | | |
| **Runtime (Minutes)** | 1108 | 1293 | 1388 | 1041 |
| **MQuad** | | | | |
| **Runtime (Minutes)** | 44 | 177 | 213 | 16 |
| **Number of identified informative variants** | 44 | 41 | 41 | 19 |
| **vireoSNP** | | | | |
| **Number of clones** | 3 | 5 | 4 | 4 |
| **Number of confidently assigned cells (>0.8)** | 3188 | 8509 | 10462 | 1397 |
| **Percentage of cells confidently assigned (%)** | 81.72 | 89.46 | 86.57 | 99.50 |

**Fig. S4 Benchmarking workflow (a) and result summary (b) of MQuad pipeline in three HEMO datasets and the hematopoietic dataset.**

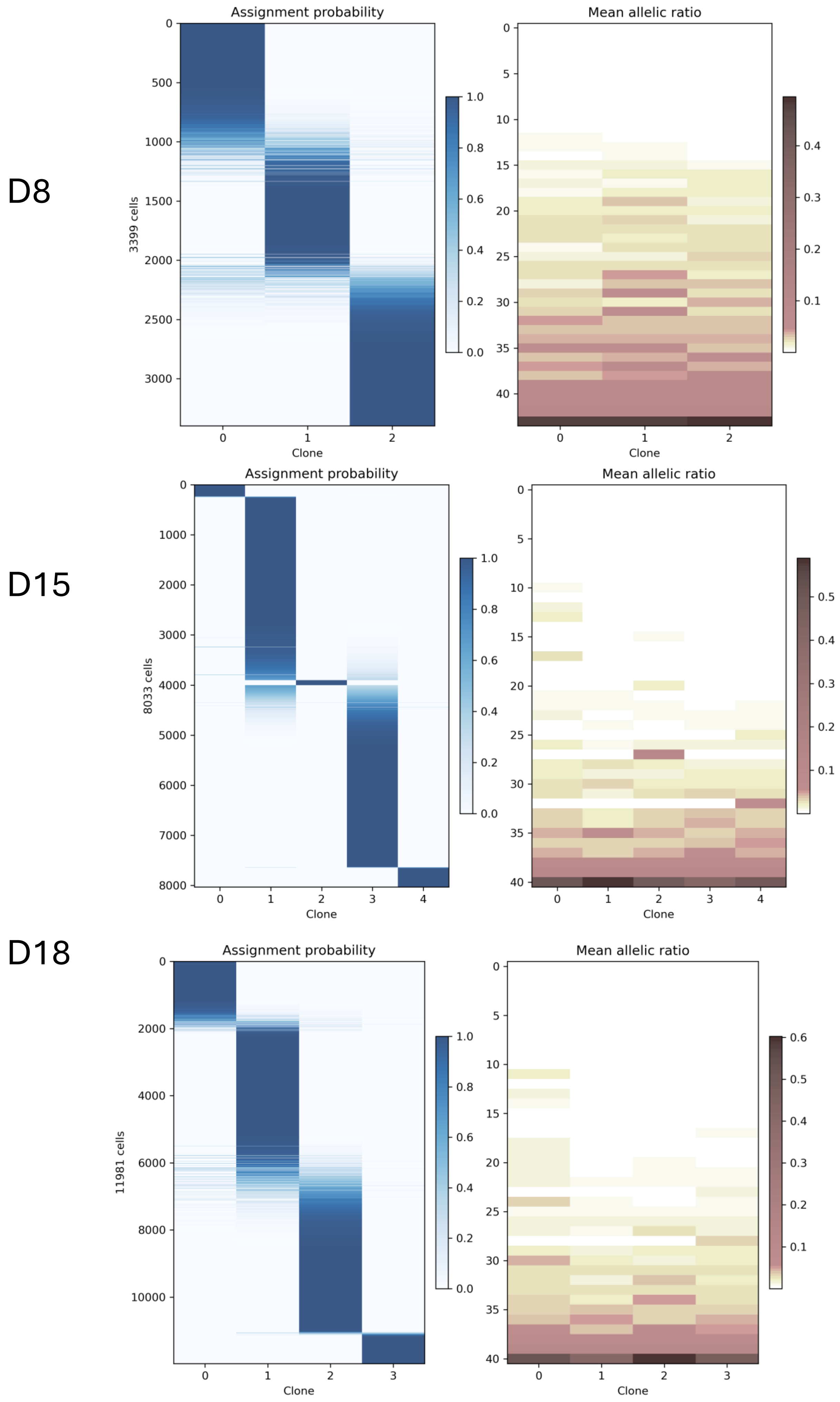
D8
Assignment probability
3399 cells
Clone
Mean allelic ratio
Clone
D15
Assignment probability
8033 cells
Clone
Mean allelic ratio
Clone
D18
Assignment probability
11981 cells
Clone
Mean allelic ratio
Clone

**Fig. S5 Benchmarking results of MQuad pipeline in three HEMO datasets.**

(Left) Heatmap shows barcode assignment with informative mtDNA variants detected with MQuad. Each row is a cell, each column is a barcode, heatmap color indicates assignment probability of cells in D8, D15 and D18 HEMOs.

(Right) Allele frequency heatmap shows the 43, 41, and 41 informative mtDNA SNVs detected by MQuad ranked from lowest to highest score of difference in Bayesian Information Criterion (ΔBIC) in each clone. Each row is a variant, each column is a barcode. Heatmap color indicates the value of the ΔBIC which is an indicator for clonal informativeness of each SNP with higher ΔBIC being more informative.

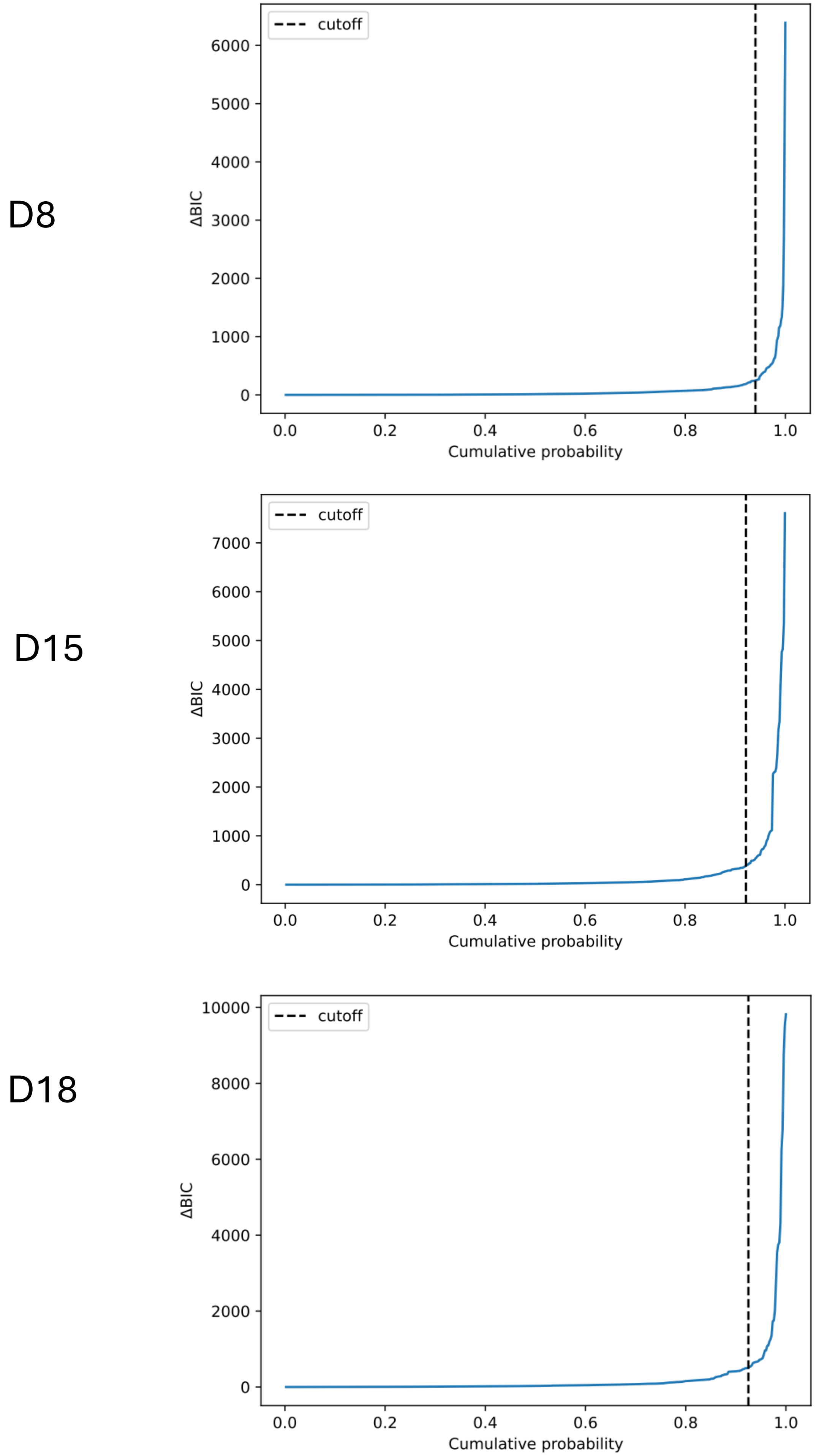


**Fig. S6 Cumulative distribution function of ΔBIC with cutoff shown generated by MQuad in three HEMO datasets.**

a

b

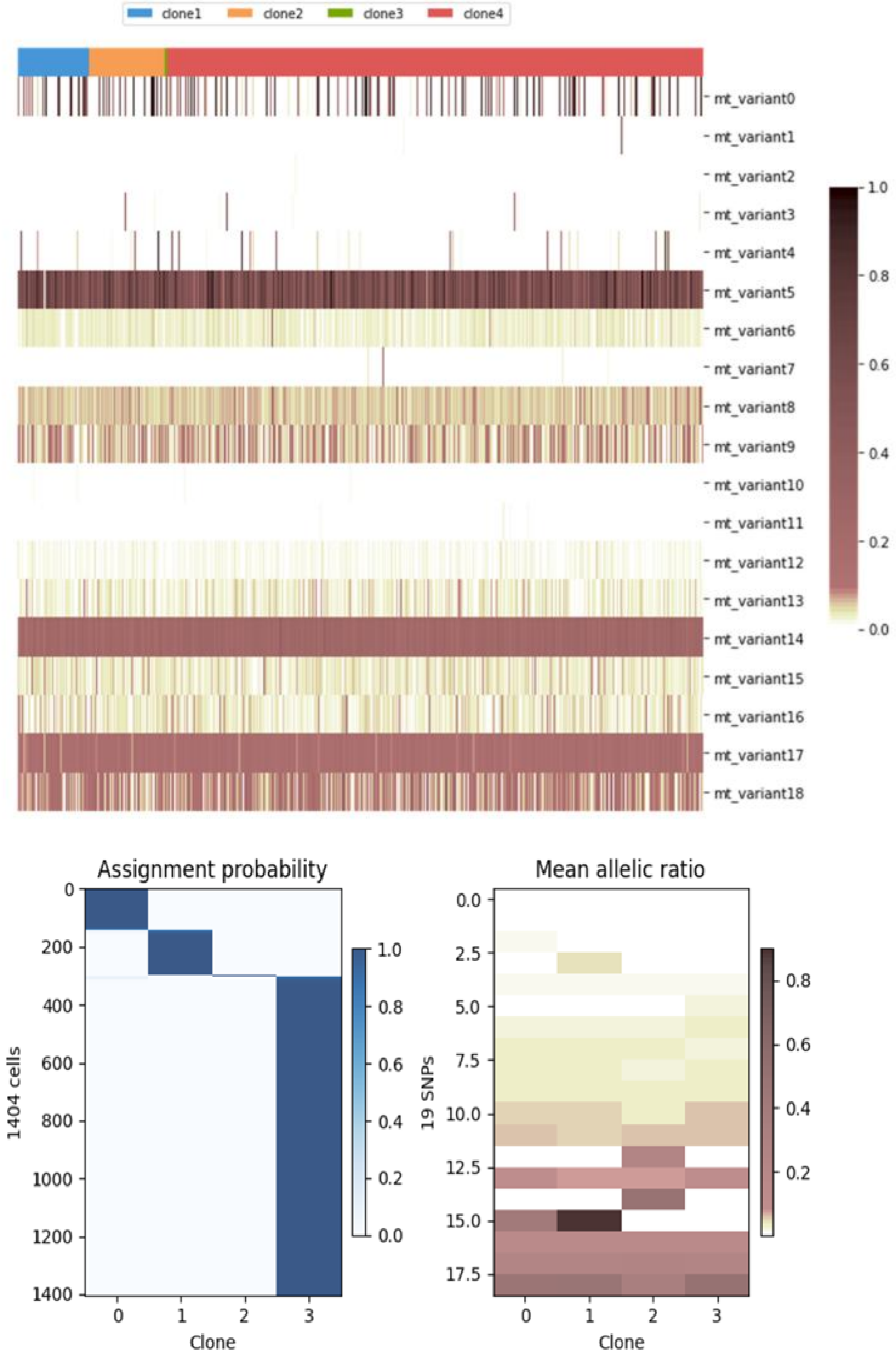

clone1
clone2
clone3
clone4
mt_variant0
mt_variant1
mt_variant2
mt_variant3
mt_variant4
mt_variant5
mt_variant6
mt_variant7
mt_variant8
mt_variant9
mt_variant10
mt_variant11
mt_variant12
mt_variant13
mt_variant14
mt_variant15
mt_variant16
mt_variant17
mt_variant18
1.0
0.8
0.6
0.4
0.2
0.0
Assignment probability
1404 cells
Clone
Mean allelic ratio
19 SNPs
Clone

**Fig. S7 Benchmarking results of MQuad pipeline in the hematopoietic dataset.**

a, Allele frequency heatmap shows the 19 informative mtDNA SNVs detected by MQuad in each clone. Each row is a variant, each column is a barcode. Heatmap color indicates the value of the allele frequency.

b, (Left) Heatmap shows barcode assignment with informative mtDNA variants detected with MQuad. Each row is a cell, each column is a barcode, heatmap color indicates assignment probability of cells.

(Right) Allele frequency heatmap shows the 19 informative mtDNA SNVs detected by MQuad ranked from lowest to highest score of difference in Bayesian Information Criterion (ΔBIC). Each row is a variant, each column is a barcode. Heatmap color indicates the value of the ΔBIC which is an indicator for clonal informativeness of each SNP with higher ΔBIC being more informative.

**a**

| | Human embryonic organoid datasets | | | Hematopoietic dataset |
|---|---|---|---|---|
| | D8 | D15 | D18 | |
| **Resulting number of variants after filtering for mean quality > 15 and mean coverage > 5** | | | | |
| **With UMI 3 consensus** | 21844 | 17531 | 15911 | 15860 |
| **Without UMI consensus** | 13838 | 14387 | 13887 | 11402 |
| **Number of variants identified with minimum VAF of 40 and minimal clone size of 5** | | | | |
| **With UMI 3 consensus** | 73 | 62 | 80 | 79 |
| **Without UMI consensus** | 19 | 45 | 62 | *No variants* |
| **Number of variants identified with minimum VAF of 40 and minimal clone size of 10** | | | | |
| **With UMI 3 consensus** | 16 | 14 | 29 | *No variants* |
| **Without UMI consensus** | 7 | 23 | 29 | *No variants* |

**b**

| | Human embryonic organoid datasets | | | Hematopoietic dataset |
|---|---|---|---|---|
| | D8 | D15 | D18 | |
| **Maegatk (with UMI consensus parameter)** | | | | |
| **Runtime (Minutes)** | 364 | 689 | 1082 | 358 |
| **Variant identification** | | | | |
| **Threshold** | *Minimum VAF – 40; Minimum clone size – 5* | | | *Min VAF – 30; min clone – 5* |
| **Number of identified informative variants** | 73 | 62 | 80 | 79 |
| **Clonal cluster assignment** | | | | |
| **Number of clones** | 6 | 7 | 4 | 3 |
| **Number of confidently assigned cells (>0.8)** | 3175 | 8009 | 10050 | 1224 |
| **Percentage of cells confidently assigned (%)** | 81.39 | 84.21 | 83.16 | 87.18 |

**Fig. S8 Benchmarking summary of meagatk pipeline in three HEMO datasets and the hematopoietic dataset.**
a, Summarizing results in comparison of the application of UMI consensus parameter in maegatk pipeline processing.
b, Summary of maegatk pipeline results on four benchmarking datasets.

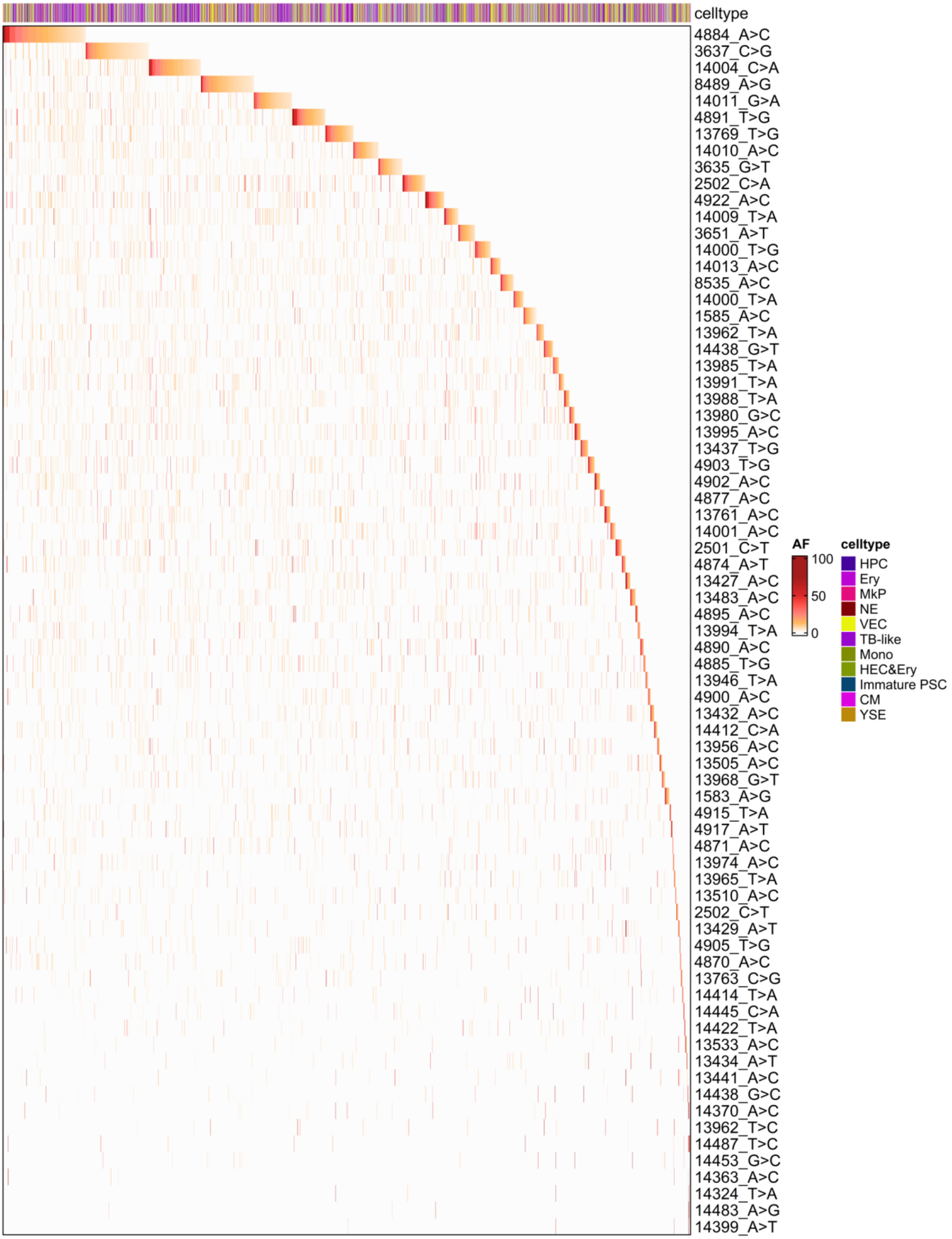


**Fig. S9 Benchmarking results of meagatk pipeline in D8 HEMO dataset.** Heatmap showing VAF value of 73 informative mtDNA variants dataset detected by maegatk organized by clones and sorted by clone size. Each row is a variant, each column is a cell. Heatmap color indicates the value of the allele frequency.

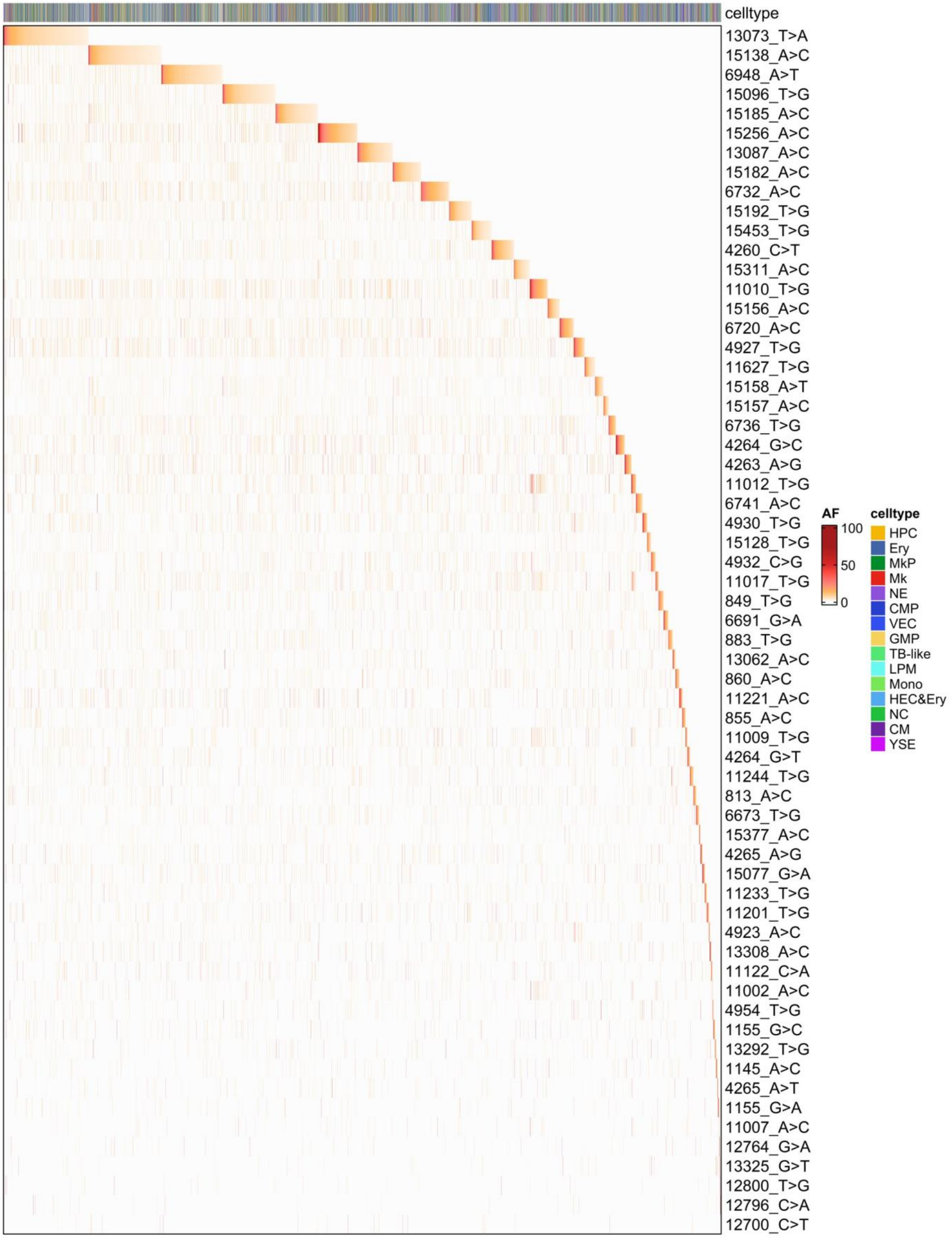


**Fig. S10 Benchmarking results of meagatk pipeline in D15 HEMO dataset.** Heatmap showing VAF value of 62 informative mtDNA variants dataset detected by maegatk organized by clones and sorted by clone size. Each row is a variant, each column is a cell. Heatmap color indicates the value of the allele frequency.

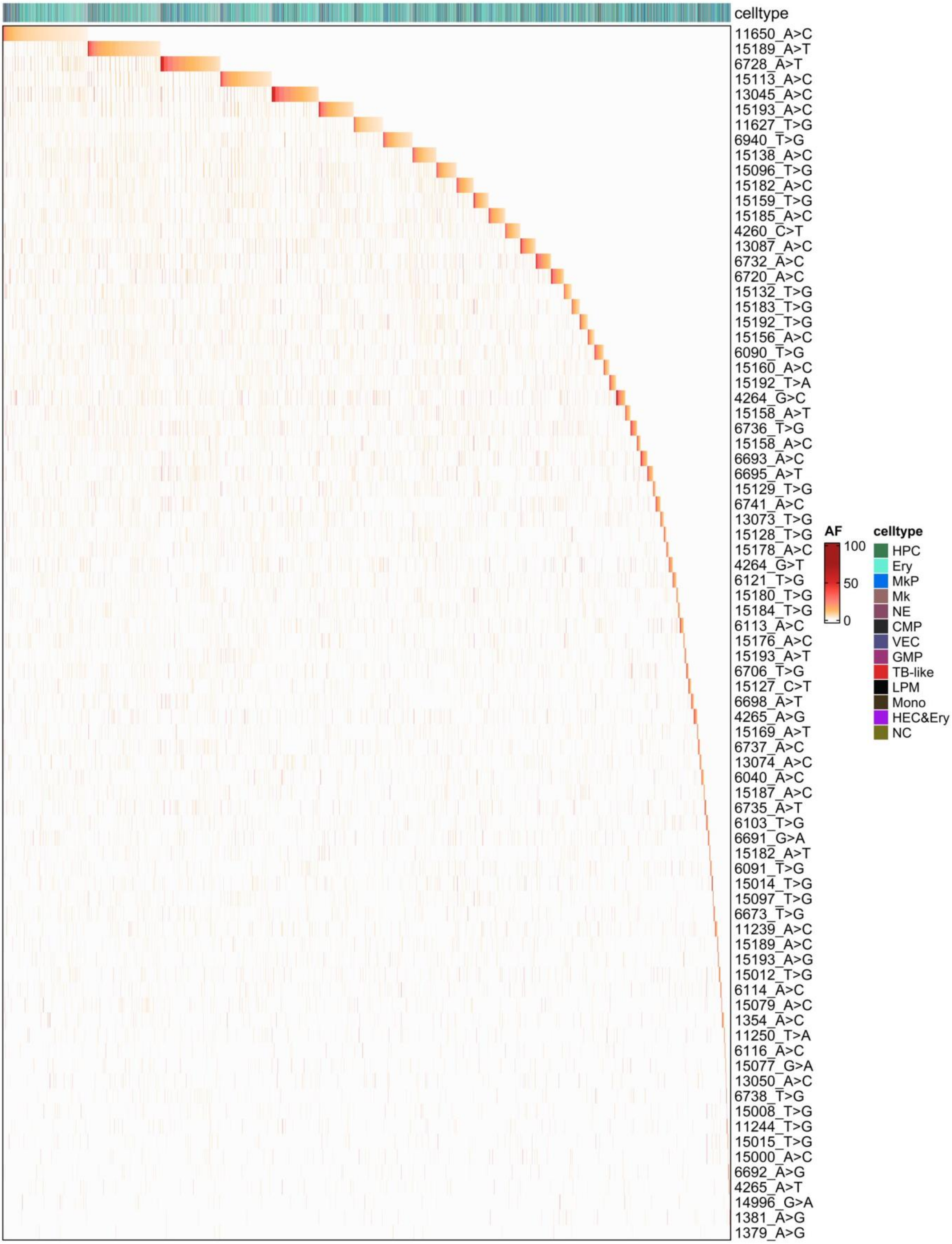


**Fig. S11 Benchmarking results of meagatk pipeline in D18 HEMO dataset.** Heatmap showing VAF value of 80 informative mtDNA variants dataset detected by maegatk organized by clones and sorted by clone size. Each row is a variant, each column is a cell. Heatmap color indicates the value of the allele frequency.

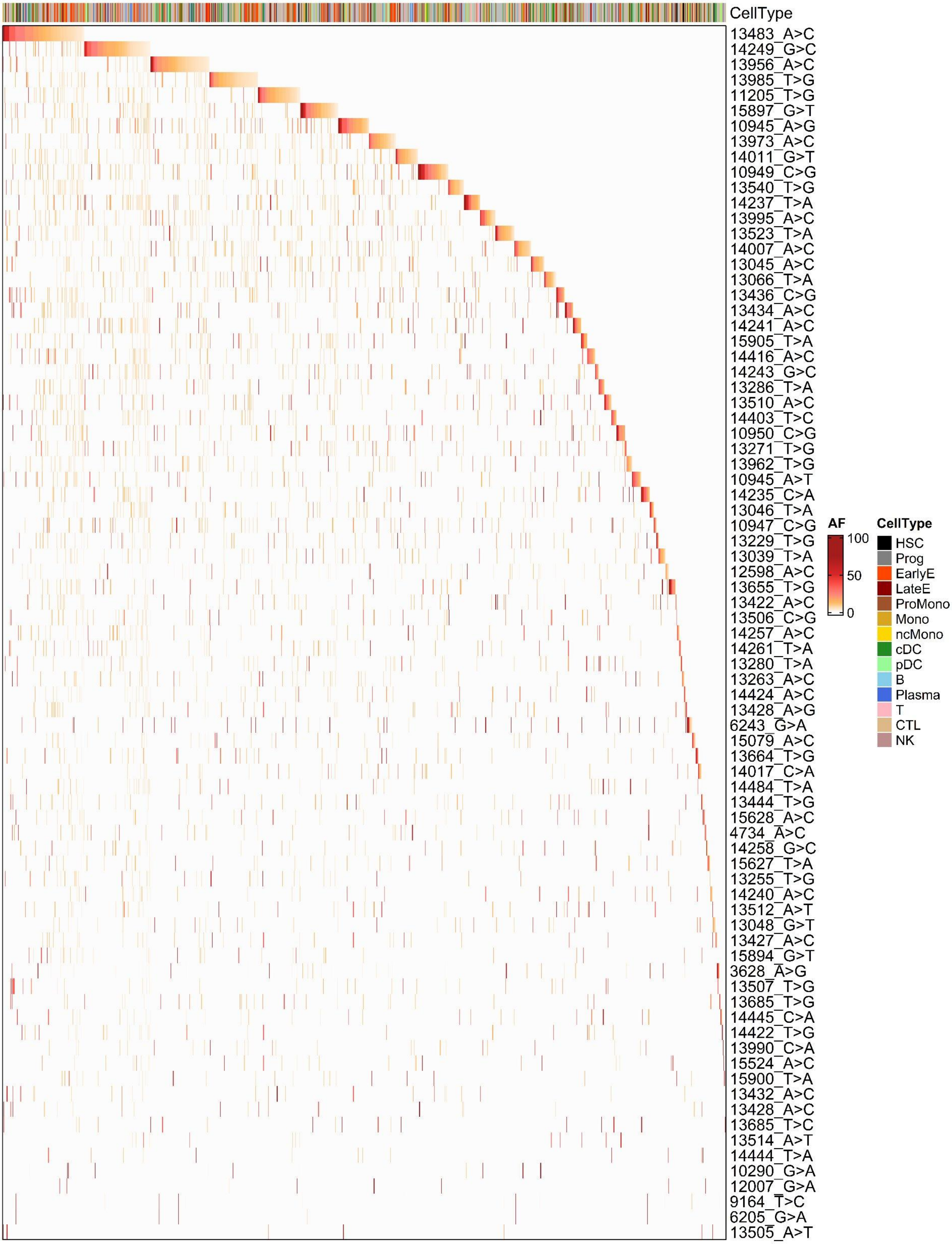


**Fig. S12 Benchmarking results of meagatk pipeline in the hematopoietic dataset.**
Heatmap showing VAF value of 79 informative mtDNA variants dataset detected by maegatk organized by clones and sorted by clone size. Each row is a variant, each column is a cell. Heatmap color indicates the value of the allele frequency.

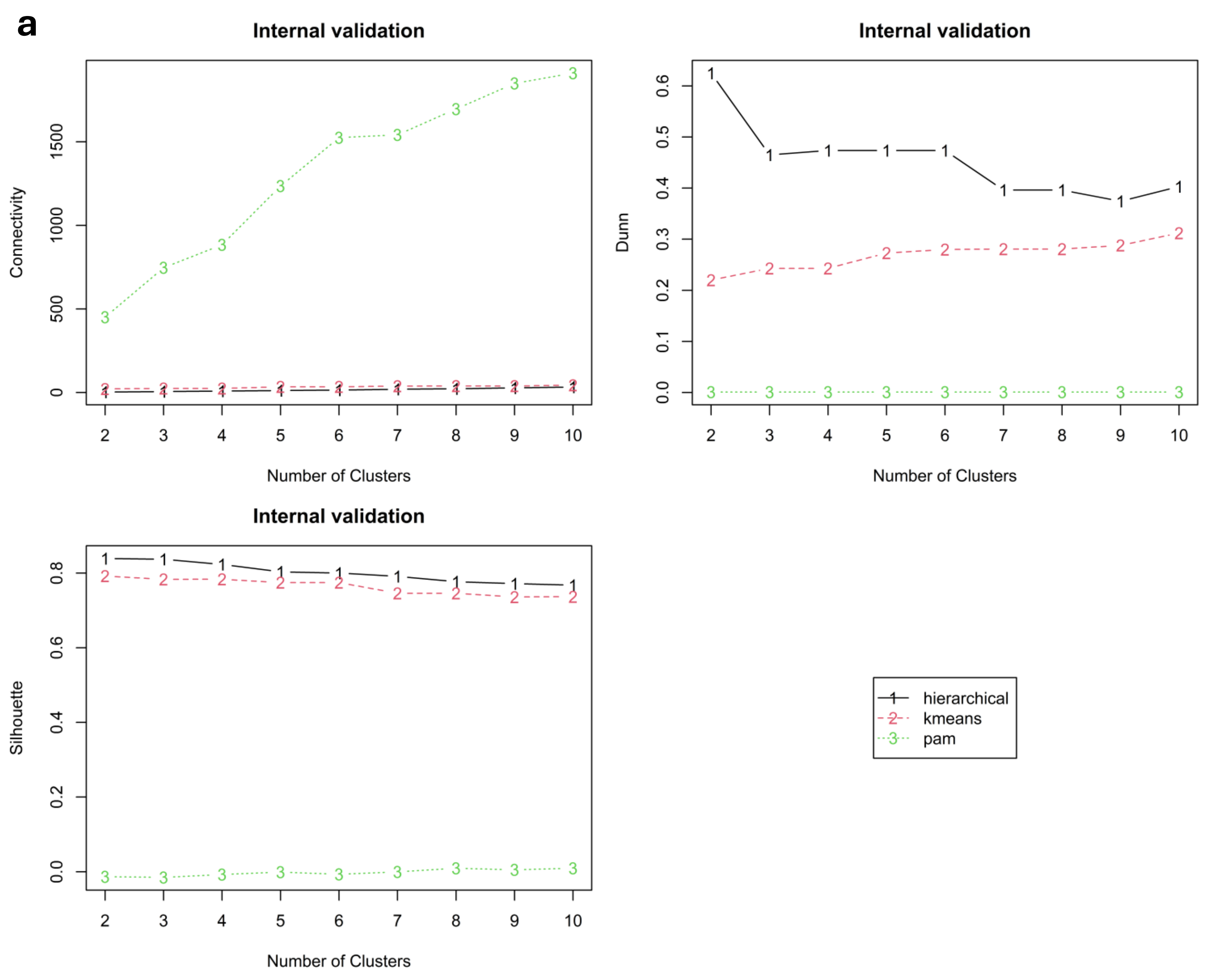


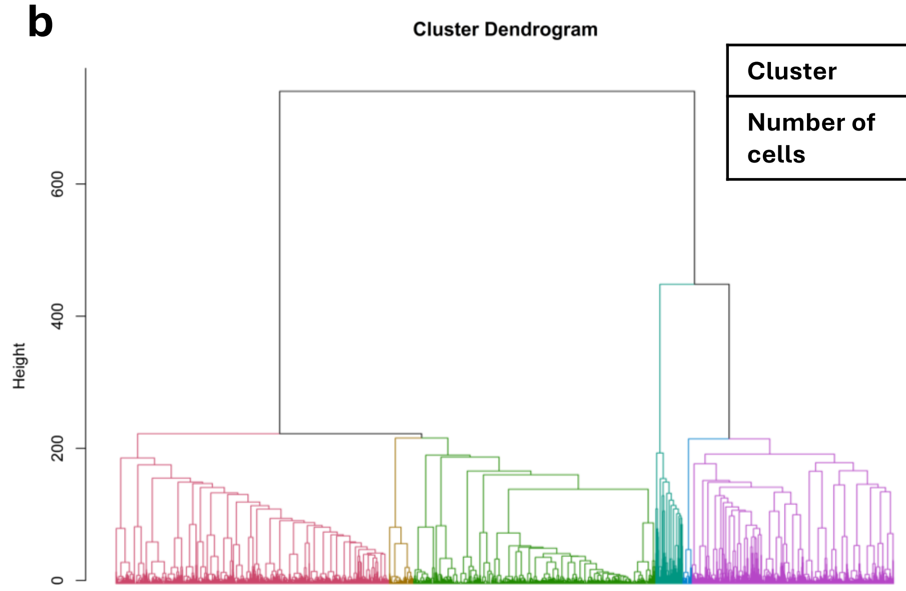


| Cluster | 1 | 2 | 3 | 4 | 5 | 6 |
|---|---|---|---|---|---|---|
| Number of cells | 1116 | 99 | 988 | 11<br>1 | 38 | 823 |

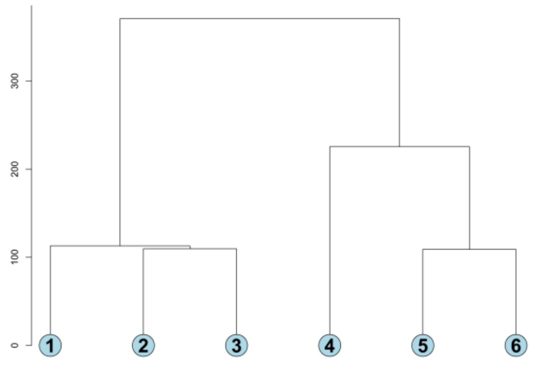


**Fig. S13 Clonal assignment in maegatk pipeline results on D8 HEMO dataset.**
a, Line plot visualizing the internal clustering results generated using clValid to determine the optimal clustering method and the number of clusters. Optimal conditions are identified with low connectivity scores and high Dunn and silhouette scores.
b, Hierarchical clustering results are visualised as dendrograms. (*Left*) Dendrogram branches are colour-segregated by clusters. (*Right*) Simplified dendrogram to visualise clustering branches.

**a**

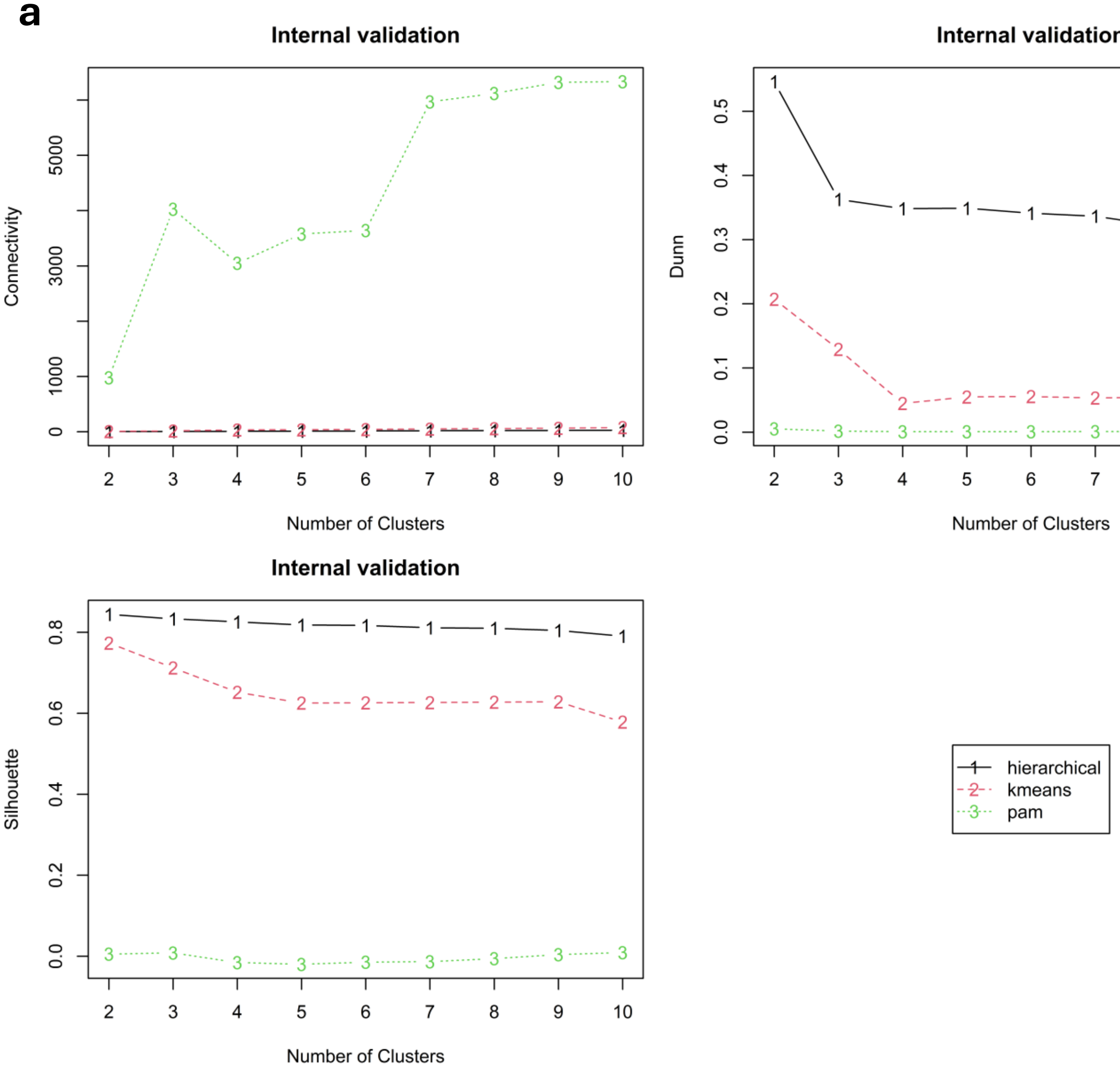


**b**

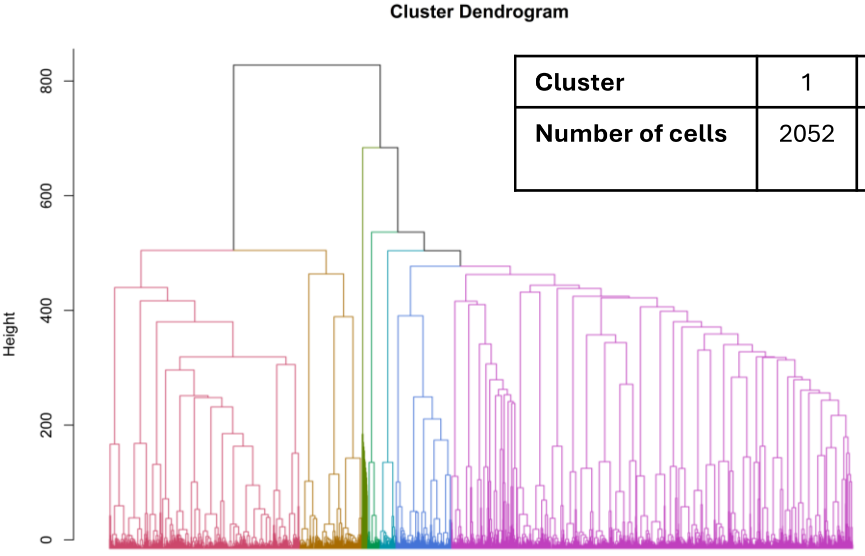


| Cluster | 1 | 2 | 3 | 4 | 5 | 6 | 7 |
|---|---|---|---|---|---|---|---|
| **Number of cells** | 2052 | 668 | 61 | 130 | 175 | 599 | 4324 |

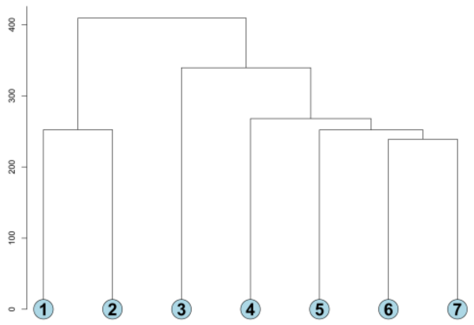


**Fig. S14 Clonal assignment in maegatk pipeline results on D15 HEMO dataset.**
a, Line plot visualizing the internal clustering results generated using clValid to determine the optimal clustering method and the number of clusters. Optimal conditions are identified with low connectivity scores and high Dunn and silhouette scores.
b, Hierarchical clustering results are visualised as dendrograms. (*Left*) Dendrogram branches are colour-segregated by clusters. (*Right*) Simplified dendrogram to visualise clustering branches.

**a**

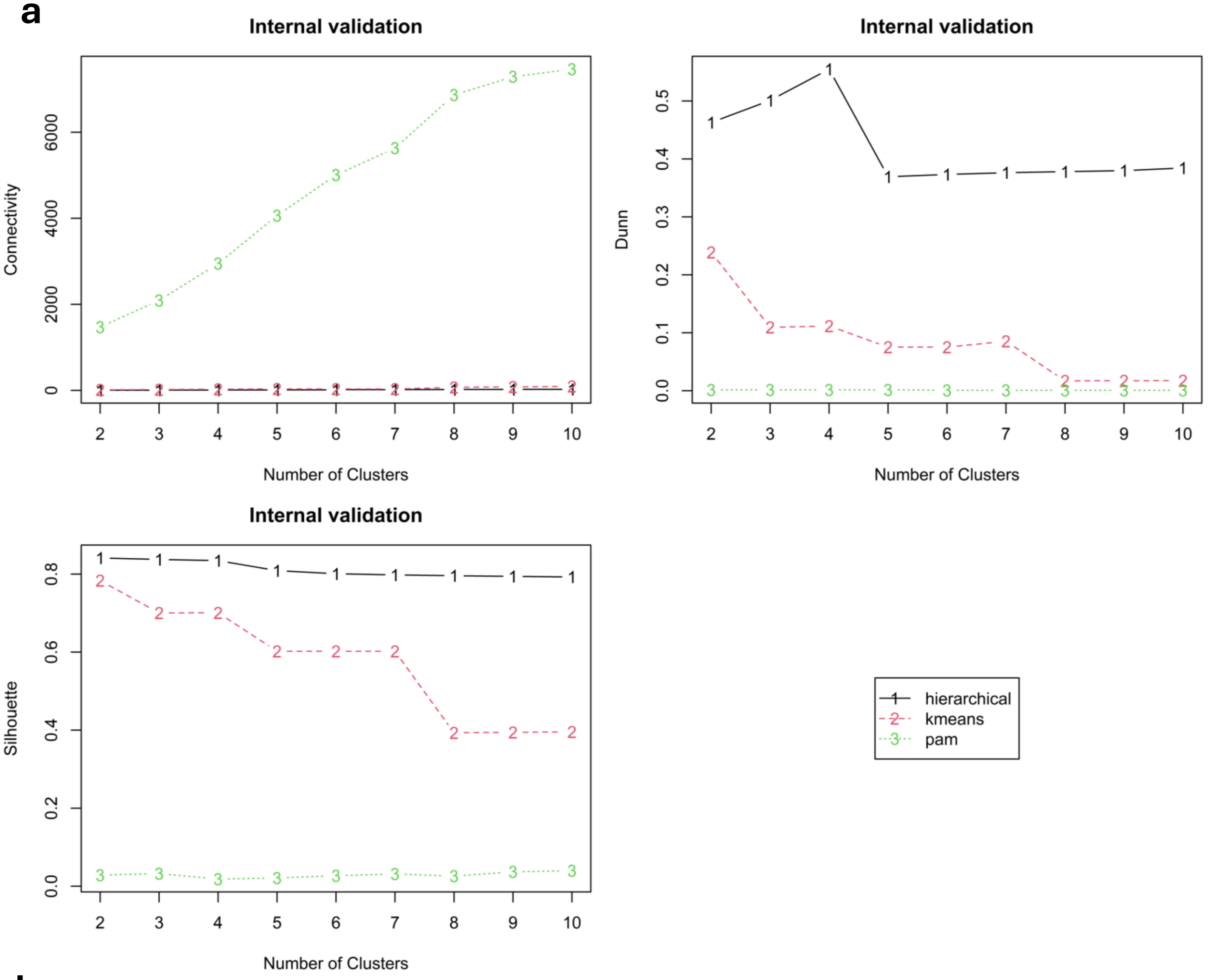


**b**

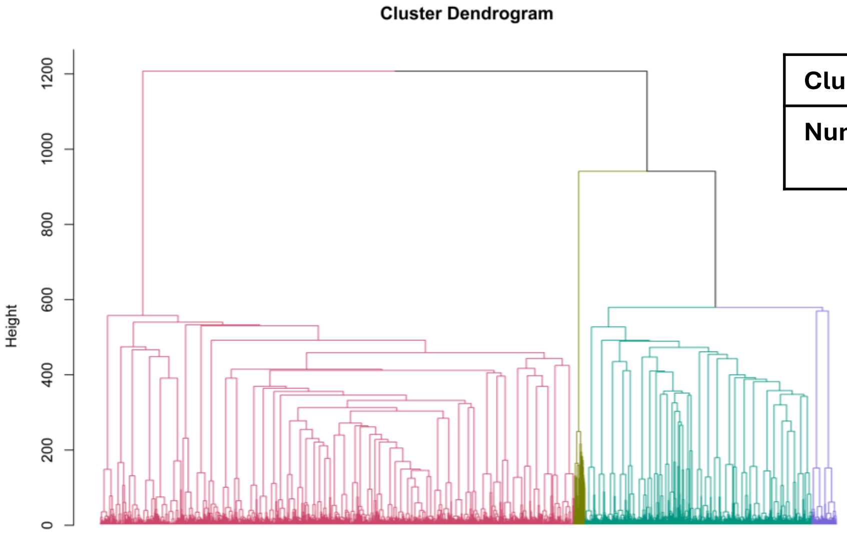


| Cluster | 1 | 2 | 3 | 4 |
|---|---|---|---|---|
| **Number of cells** | 6470 | 149 | 3097 | 334 |

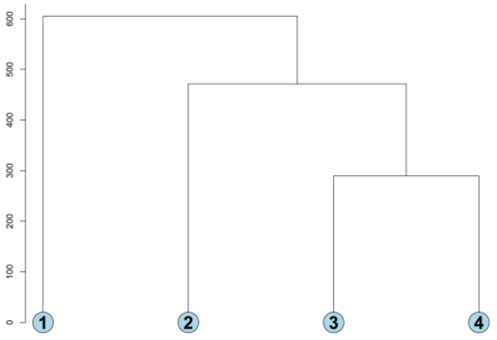

**Fig. S15 Clonal assignment in maegatk pipeline results on D18 HEMO dataset.**
a, Line plot visualizing the internal clustering results generated using clValid to determine the optimal clustering method and the number of clusters. Optimal conditions are identified with low connectivity scores and high Dunn and silhouette scores.
b, Hierarchical clustering results are visualised as dendrograms. (*Left*) Dendrogram branches are colour-segregated by clusters. (*Right*) Simplified dendrogram to visualise clustering branches.

a

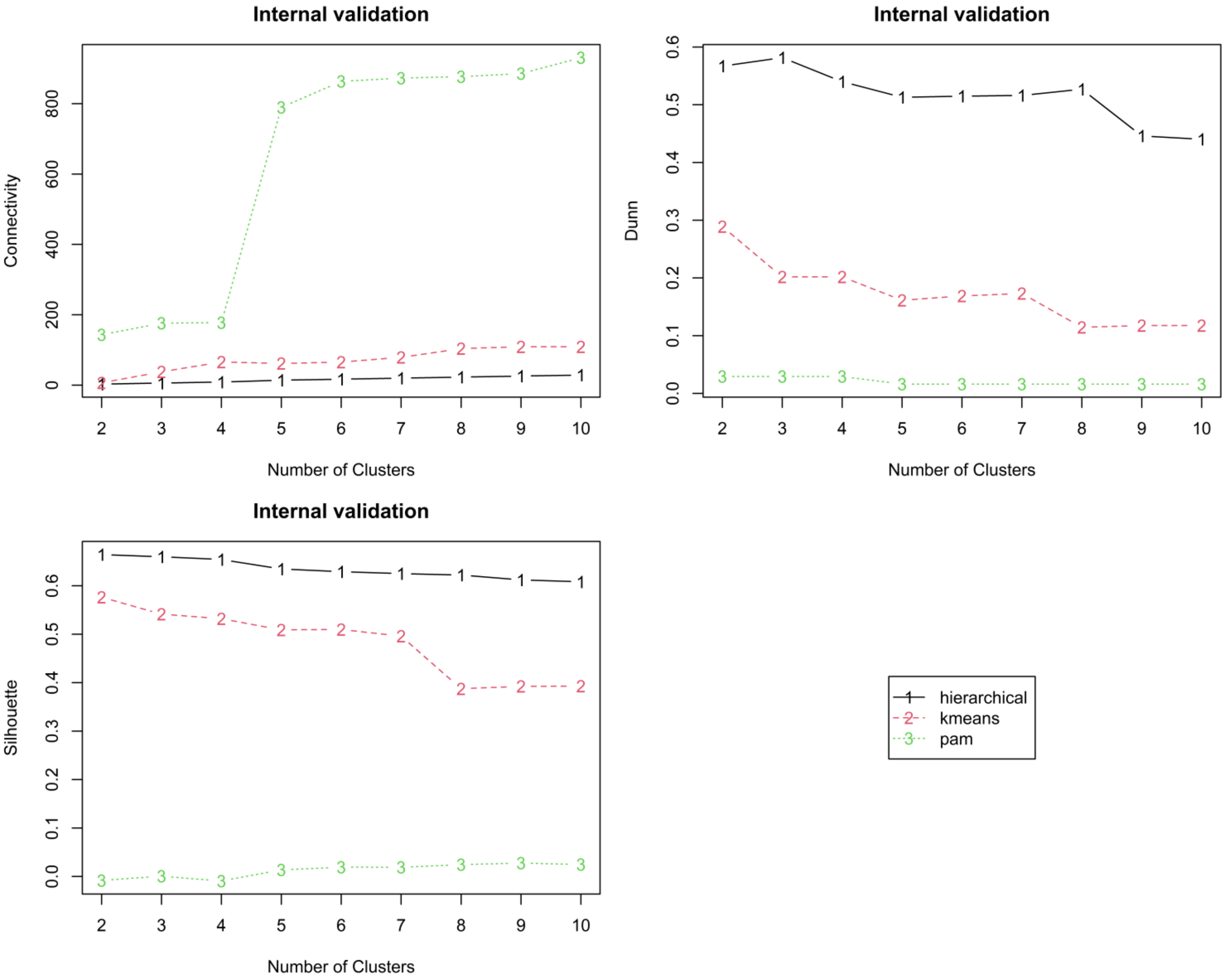


b

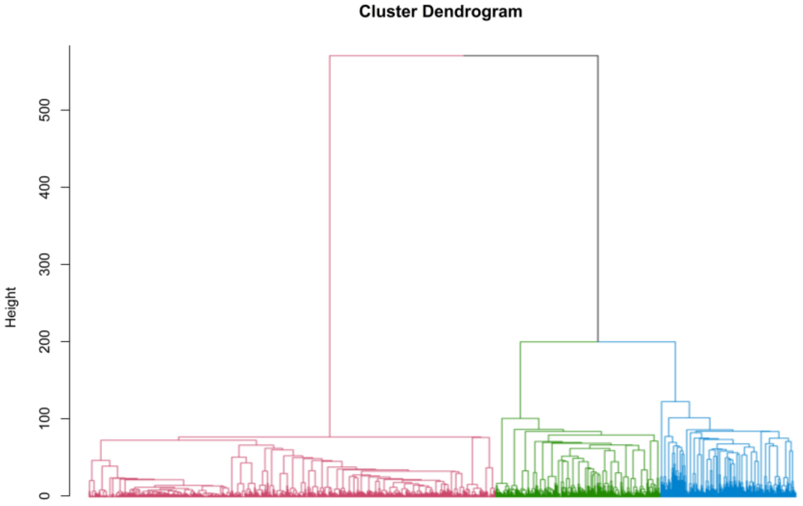


| Cluster | 1 | 2 | 3 |
|---|---|---|---|
| Number of cells | 704 | 285 | 235 |

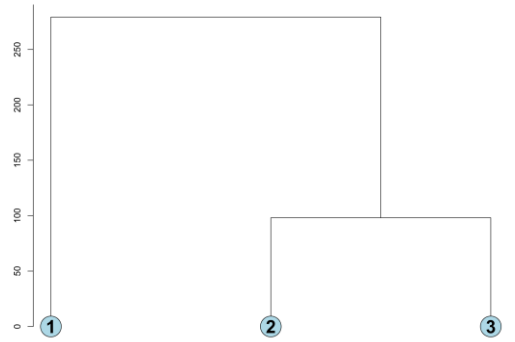

**Fig. S16 Clonal assignment in maegatk pipeline results on the hematopoietic dataset.**
a, Line plot visualizing the internal clustering results generated using clValid to determine the optimal clustering method and the number of clusters. Optimal conditions are identified with low connectivity scores and high Dunn and silhouette scores.
b, Hierarchical clustering results are visualised as dendrograms. (*Left*) Dendrogram branches are colour-segregated by clusters. (*Right*) Simplified dendrogram to visualise clustering branches.

| | Human embryonic organoid datasets | | | Hematopoietic dataset |
|---|---|---|---|---|
| | D8 | D15 | D18 | |
| **Variant identification** | | | | |
| **Number of variants from MQuad pipeline** | 44 | 41 | 41 | 19 |
| **Number of variants from maegatk pipeline** | 73 | 62 | 80 | 79 |
| **Number of overlapping variants** | **1** | **1** | **0** | **2** |
| **Clonal assignment** | | | | |
| **Number of clones identified from MQuad pipeline** | 3 | 5 | 4 | 4 |
| **Number of clones identified from maegatk pipeline** | 6 | 7 | 4 | 3 |

**Fig. S17 Summary of the mtDNA SNV variants and clones identified by the MQuad pipeline and maegatk pipeline respectively.**

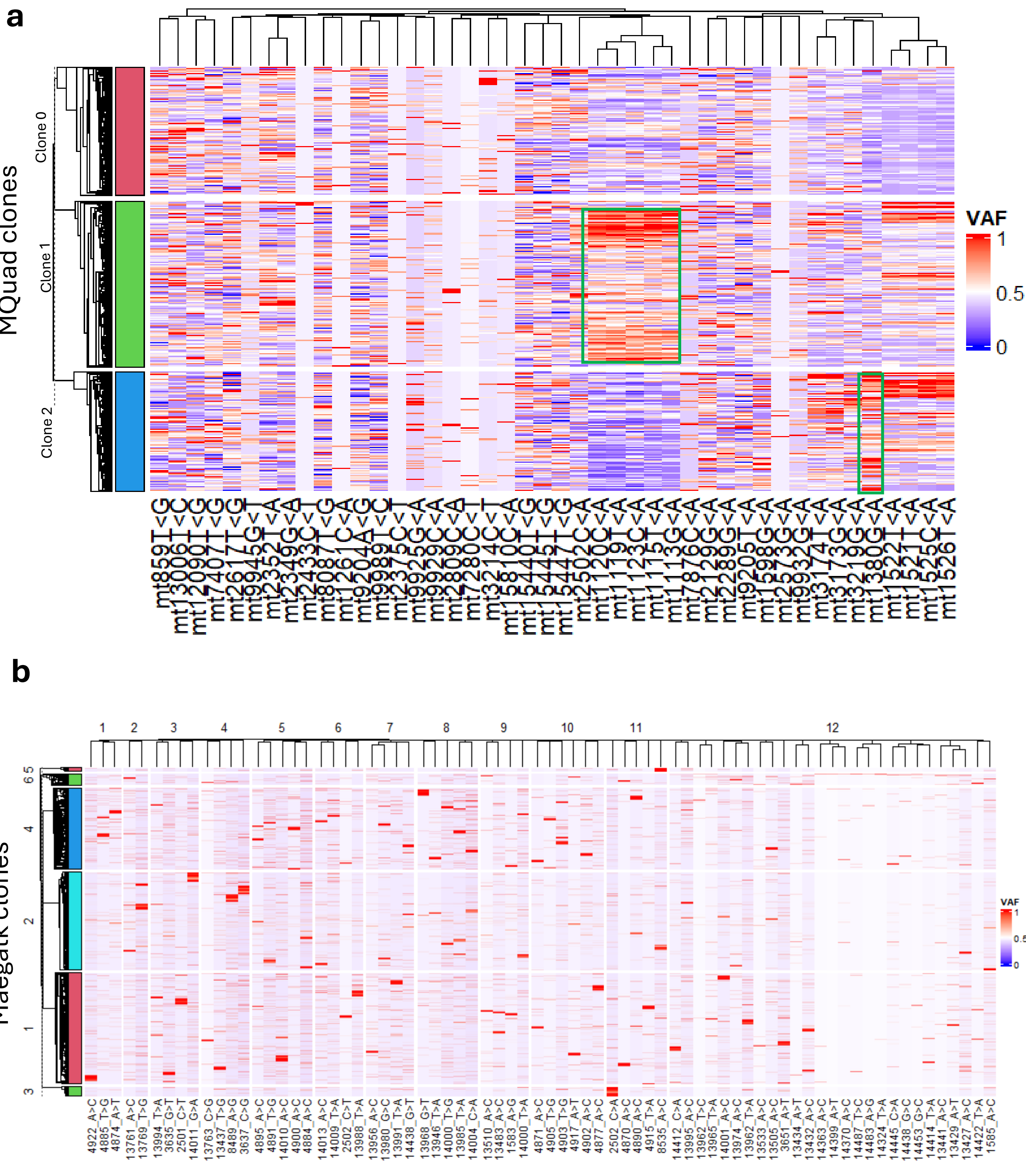


**Fig. S18 Validation of clonal clusters identified by both pipelines in D8 HEMO dataset.**

a, Heatmap constructed using scaled VAF values from MQuad pipeline, green boxes highlight clone-specific variants identified visually.

b, Similar heatmap using variants identified using the maegatk pipeline.

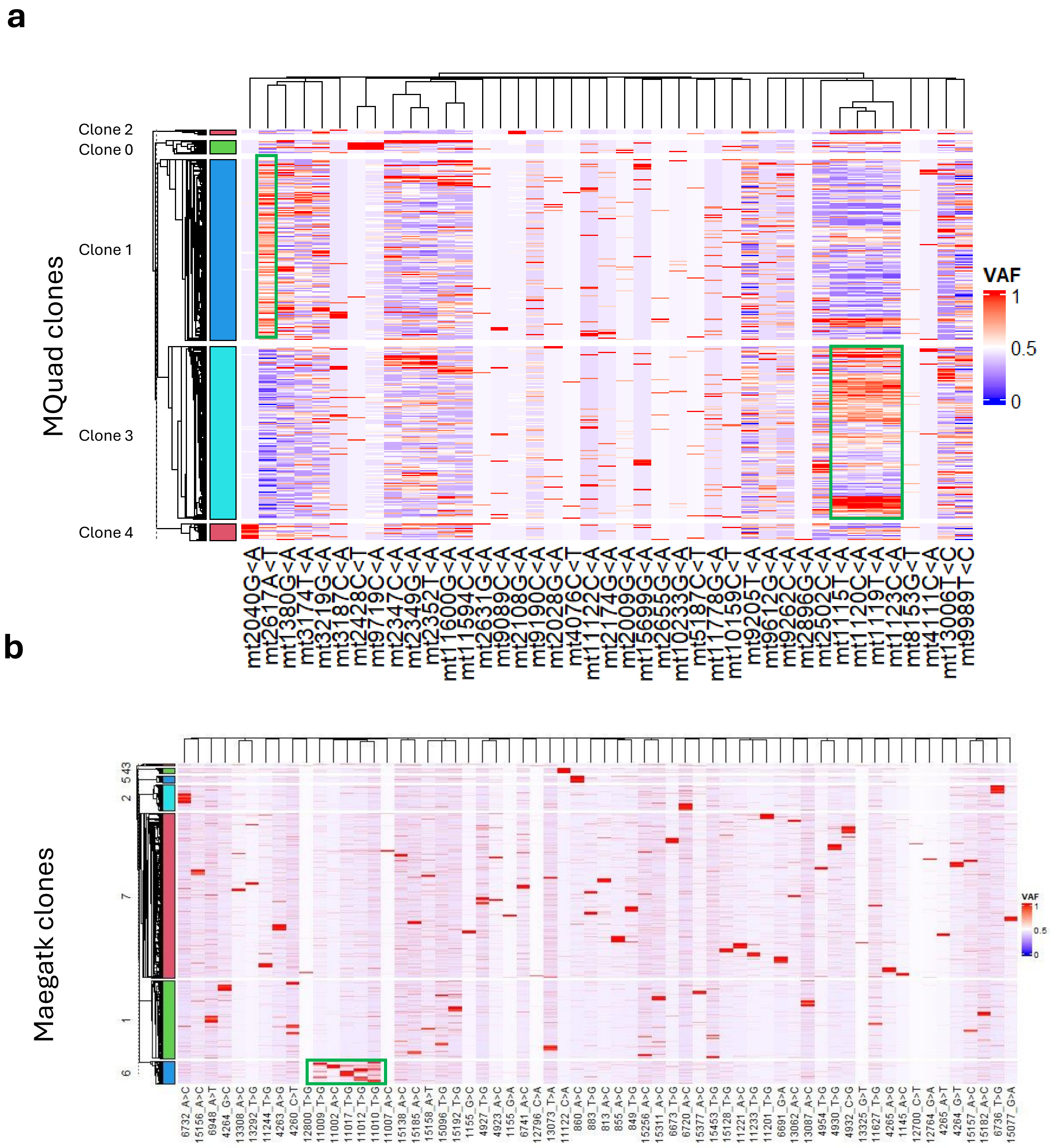


**Fig. S19 Cluster validation of results from both pipelines using the D15 HEMO dataset.**
a, Heatmap constructed using scaled VAF values from MQuad pipeline, green boxes highlight clone-specific variants identified visually.
b, Similar heatmap using variants identified using the maegatk pipeline.

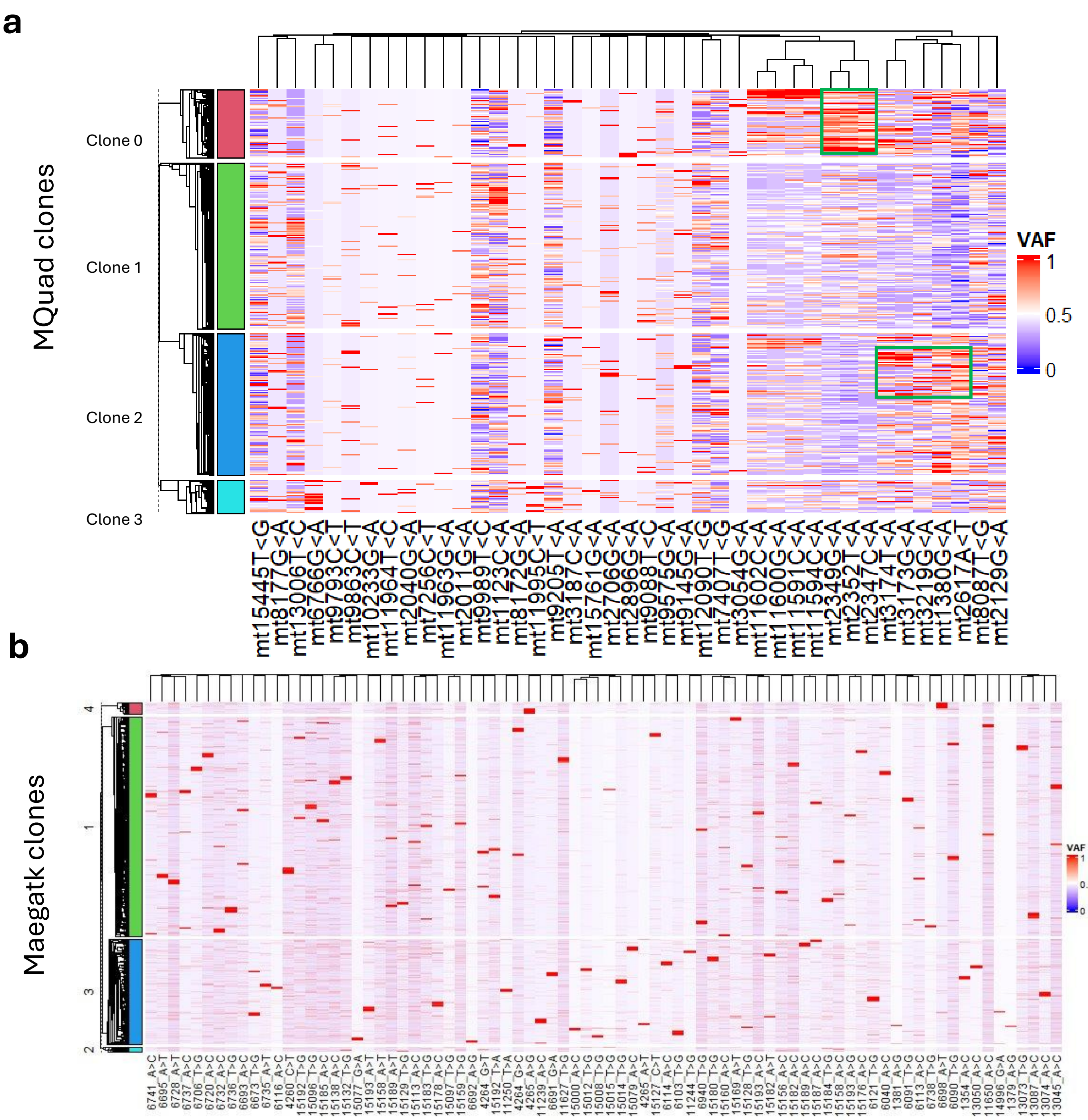


**Fig. S20 Cluster validation of results from both pipelines using the D18 HEMO dataset.**

a, Heatmap constructed using scaled VAF values from MQuad pipeline, green boxes highlight clone-specific variants identified visually.

b, Similar heatmap using variants identified using the maegatk pipeline.

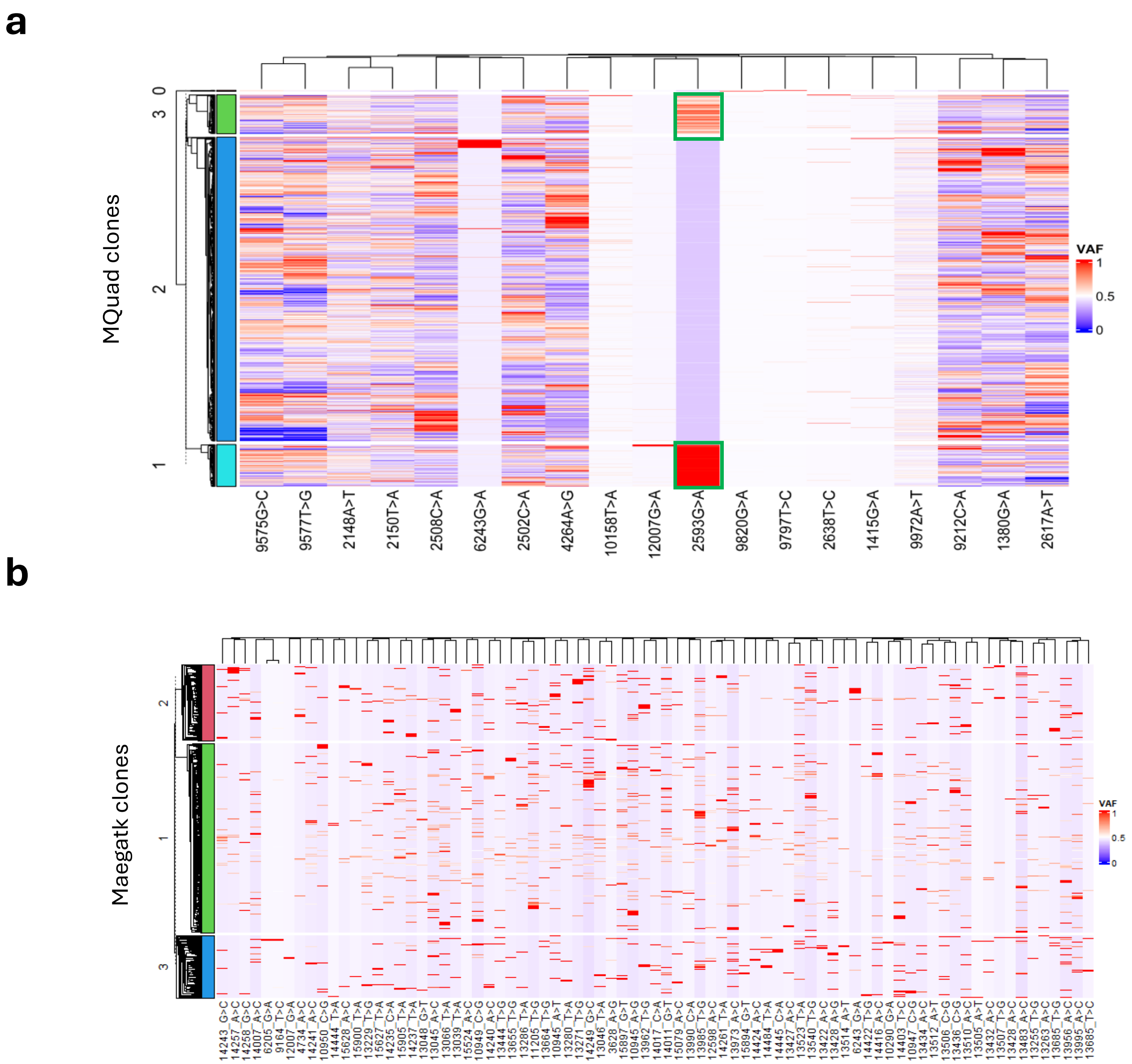


**Fig. S21 Cluster validation of results from both pipelines using the hematopoietic dataset.**
a, Heatmap constructed using scaled VAF values from MQuad pipeline, green boxes highlight clone-specific variants identified visually.
b, Similar heatmap using variants identified using the maegatk pipeline.

### Davies-Bouldin’s (DB) index

| Pipeline | HEMO datasets | | | Hematopoietic dataset |
|---|---|---|---|---|
| | D8 | D15 | D18 | |
| MQuad | 4.38 | 3.59 | 4.58 | 5.60 |
| maegatk | 7.15 | 6.01 | 8.75 | 9.61 |

**Fig. S22 Quantitative comparison of internal clustering metrics from both pipelines using Davies-Bouldin’s (DB) index in four datasets in cluster validation.**

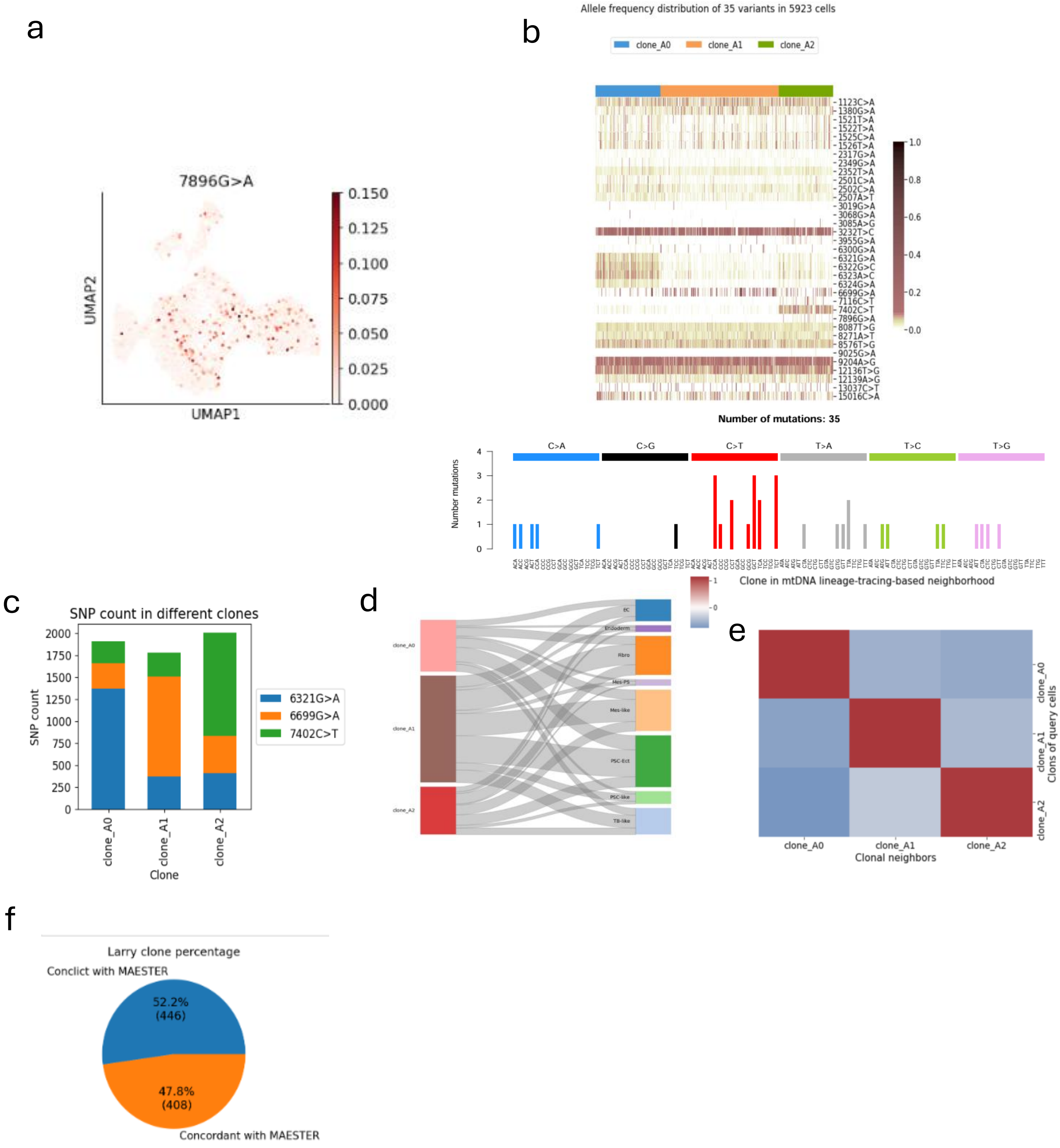
a
7896G>A
UMAP2
UMAP1
0.150
0.125
0.100
0.075
0.050
0.025
0.000
b
Allele frequency distribution of 35 variants in 5923 cells
clone_A0
clone_A1
clone_A2
1123C>A
1380G>A
1521T>A
1522T>A
1525C>A
1526T>A
2317G>A
2349G>A
2352T>A
2501C>A
2502C>A
2507A>T
3019G>A
3068G>A
3085A>G
3232T>C
3955G>A
6300G>A
6321G>A
6322G>C
6323A>C
6324G>A
6699G>A
7116C>T
7402C>T
7896G>A
8087T>G
8271A>T
8576T>G
9025G>A
9204A>G
12136T>G
12139A>G
13037C>T
15016C>A
1.0
0.8
0.6
0.4
0.2
0.0
Number of mutations: 35
Number mutations
C>A
C>G
C>T
T>A
T>C
T>G
c
SNP count in different clones
SNP count
6321G>A
6699G>A
7402C>T
clone_A0
clone_A1
clone_A2
Clone
d
clone_A0
clone_A1
clone_A2
EC
Endoderm
Fibro
Mes-PS
Mes-like
PSC-Ect
PSC-like
TB-like
e
Clone in mtDNA lineage-tracing-based neighborhood
Clones of query cells
Clonal neighbors
clone_A0
clone_A1
clone_A2
f
Larry clone percentage
Conclict with MAESTER
52.2%
(446)
47.8%
(408)
Concordant with MAESTER

**Fig. S23 Clonality barcoded by mitochondrial variants in the early stage HEMOs (D4).**

a, UMAP visualization reveals the uneven distribution of representative mtDNA SNVs across various cell types, suggesting their potential to distinguish clonal lineages with high resolution. These findings underscore the utility of mtDNA variants as stable lineage markers.

b, Allele frequency heatmap displays 35 informative mtDNA SNVs detected by MQuad in three clones from early-stage HEMOs. Each row represents a variant, and each column corresponds to a barcode. Heatmap color indicates allele frequency values, highlighting the heterogeneity of variant distributions.

c, Bar chart illustrates that 6321G>A, 6693G>A, and 7402C>T are the most prevalent variants in Clone_A0, Clone_A1, and Clone_A2, respectively, reflecting clone-specific mutational signatures.

d, Alluvial plot demonstrates that Mesoderm and Ectoderm are the predominant lineages in the three clones traced by mtDNA variant enrichment during early hematopoiesis, indicating lineage commitment at this stage.

e, Neighborhood analysis reveals high intra-clone connectedness, indicating robust clonal identity within each clone and supporting the accuracy of mitochondrial variant-based lineage tracing.

f, Pie chart highlights significant divergence between LARRY-defined clones and mitochondrial variant-based clonality.

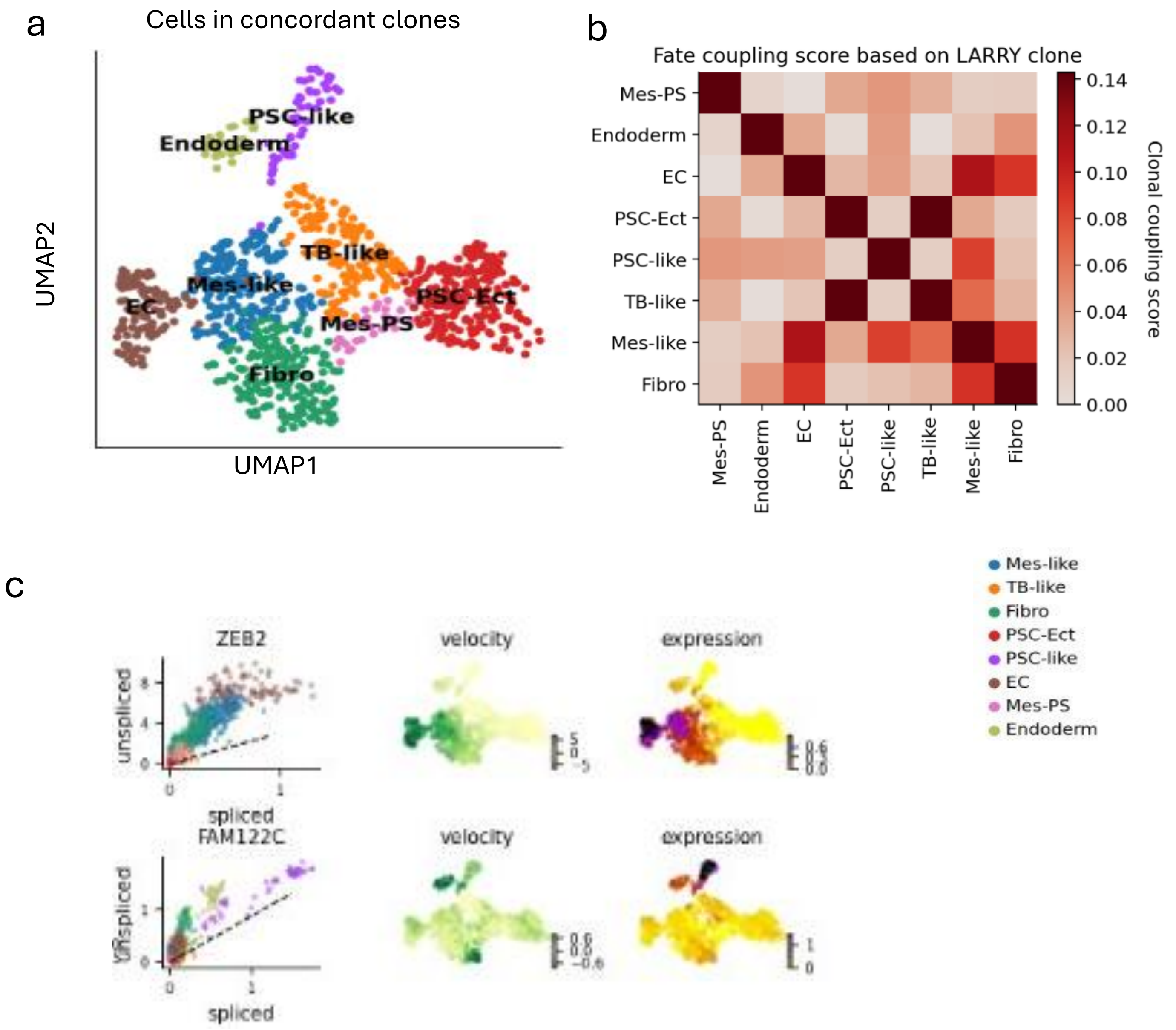


**Fig. S24 Enhanced clonal resolution with mitochondrial variants in early stage HEMO (D4).**
a, The UMAP plot highlights clear cell type boundaries when clonality analysis is aided by mitochondrial variants, demonstrating their ability to resolve ambiguities in lineage tracing.
b, Mitochondrial variants distinguish coupled cell fates. The heatmap demonstrates that mitochondrial variants successfully distinguish between the cell fates of EC and Mes-like cells, which were otherwise coupled together in earlier analyses.
c, Putative driver genes are identified by high likelihoods. Phase portraits, velocity and expression dynamics for these driver genes characterize their activity. ZEB2 explains the directionality in the up-regulated Mes-like (blue) to EC (grey).

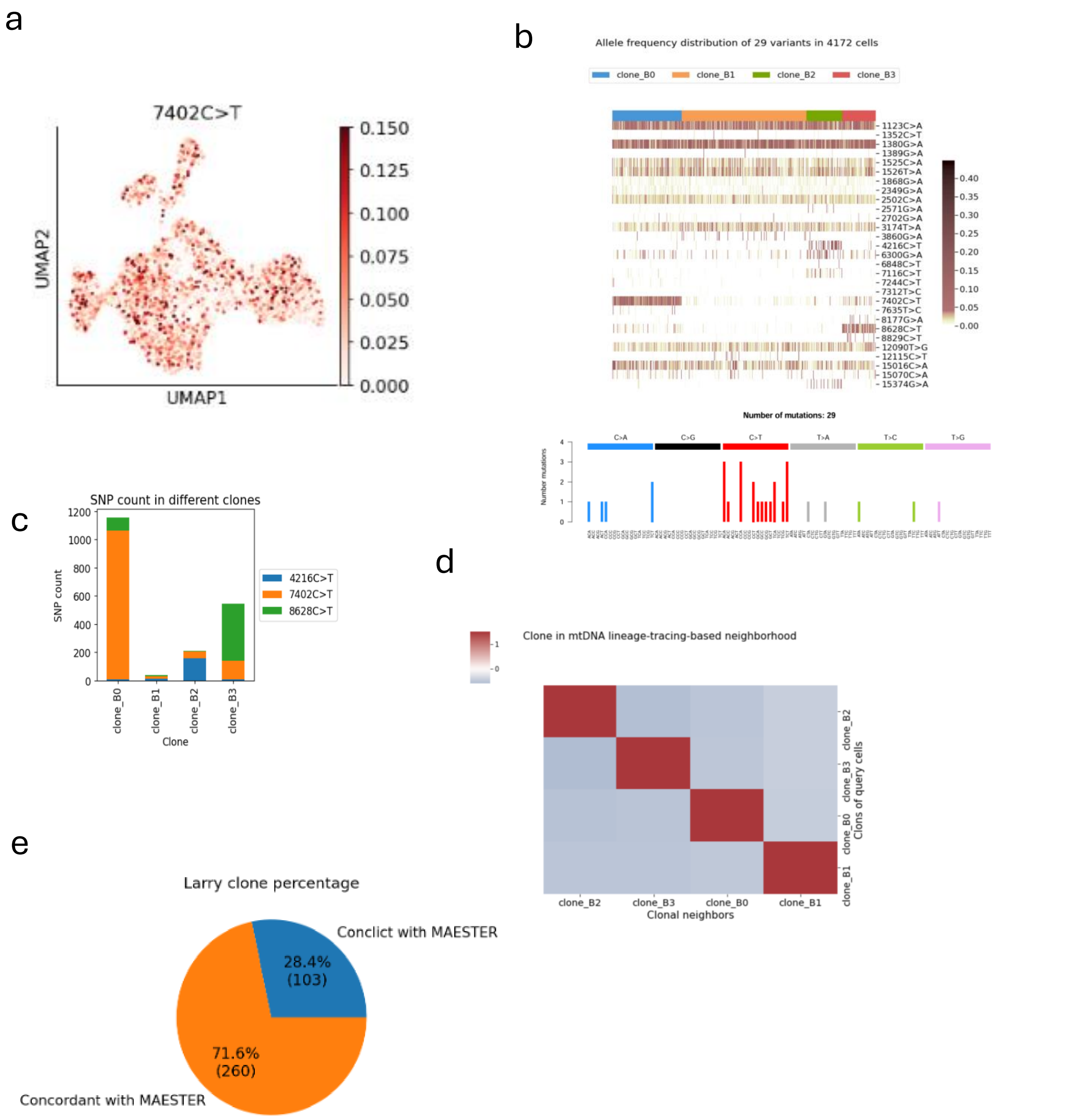


**Fig. S25 Clonality barcoded by mitochondrial variants in the late stage HEMOs (D8).**

a, UMAP visualization reveals the uneven distribution of representative mtDNA SNVs across various cell types, suggesting their potential to distinguish clonal lineages with high resolution. These findings underscore the utility of mtDNA variants as stable lineage markers.

b, Allele frequency heatmap displays 29 informative mtDNA SNVs detected by MQuad in three clones from late stage HEMOs. Each row represents a variant, and each column corresponds to a barcode.

c, Bar chart illustrates that 4216C>T, 7402C>T, and 8628C>T are the most prevalent variants in Clone_B0, Clone_B2 and Clone_B3 respectively, reflecting clone-specific mutational signatures.

d, Neighborhood analysis reveals high intra-clone connectedness, indicating robust clonal identity within each clone and supporting the accuracy of mitochondrial variant-based lineage tracing.

e, Pie chart highlights significant divergence between LARRY-defined clones and mitochondrial variant-based clonality.

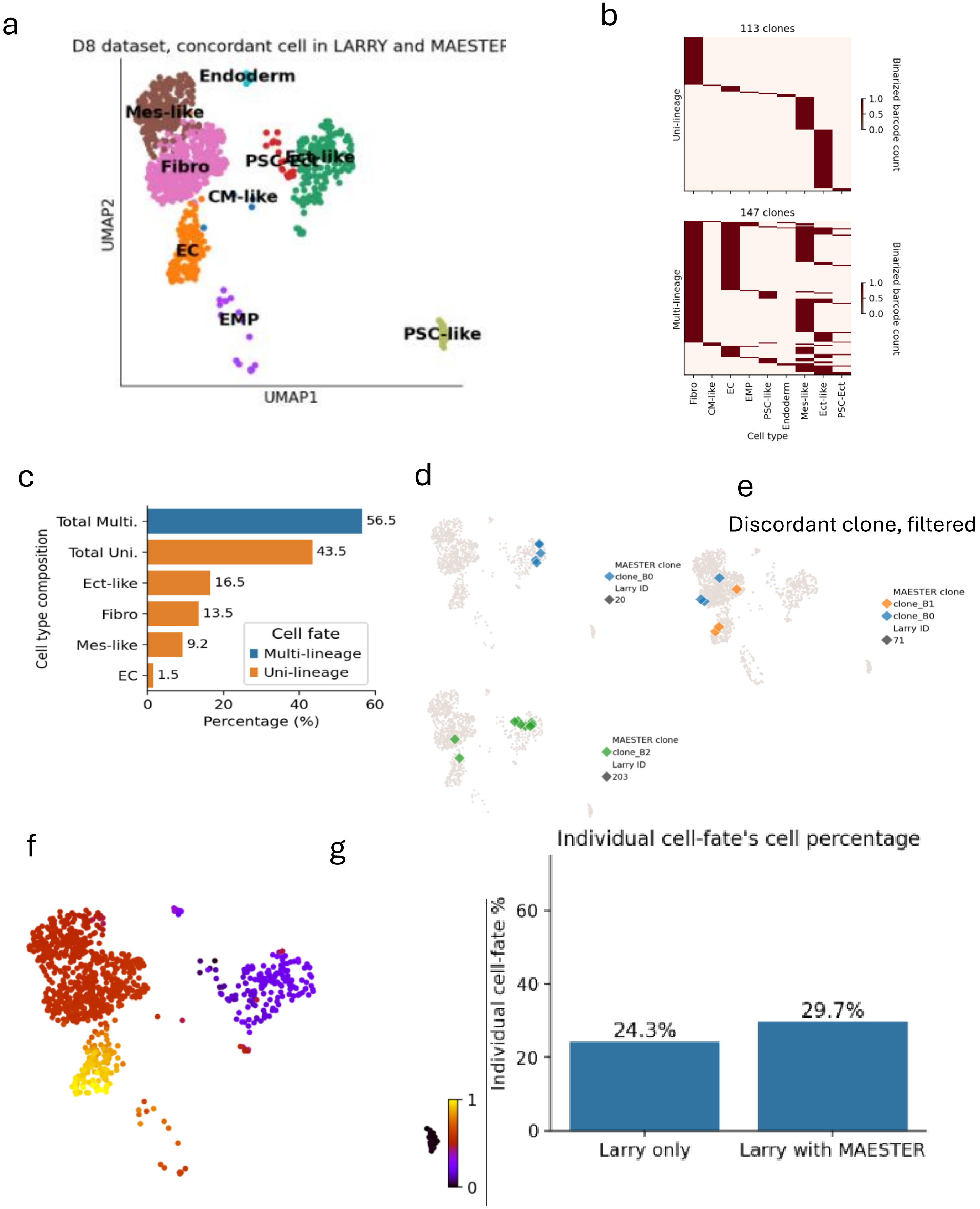
a
D8 dataset, concordant cell in LARRY and MAESTER
Endoderm
Mes-like
Fibro
PSC-Ect
Ect-like
CM-like
EC
EMP
PSC-like
UMAP2
UMAP1
b
113 clones
Uni-lineage
147 clones
Multi-lineage
Binarized barcode count
1.0
0.5
0.0
Fibro
CM-like
EC
EMP
PSC-like
Endoderm
Mes-like
Ect-like
PSC-Ect
Cell type
c
Cell type composition
Total Multi.
Total Uni.
Ect-like
Fibro
Mes-like
EC
56.5
43.5
16.5
13.5
9.2
1.5
Cell fate
Multi-lineage
Uni-lineage
0
20
40
60
Percentage (%)
d
MAESTER clone
clone_B0
Larry ID
20
MAESTER clone
clone_B2
Larry ID
203
e
Discordant clone, filtered
MAESTER clone
clone_B1
clone_B0
Larry ID
71
f
1
0
g
Individual cell-fate's cell percentage
Individual cell-fate %
60
40
20
0
24.3%
29.7%
Larry only
Larry with MAESTER

**Fig. S26 Enhanced clonal resolution with mitochondrial variants in late stage HEMOs (D8).**

a, UMAP plot highlights clear cell type boundaries when clonality analysis is aided by mitochondrial variants, demonstrating their ability to resolve ambiguities in lineage tracing.
b, Heatmaps showing refined clonal fates after correction with mitochondrial variants. Each row represents a LARRY clone, and each column corresponds to a specific lineage.
c, Bar chart showing the proportion of mitochondrial variant refined clones in each lineage.
d, UMAP plots showing the example mitochondrial variant refined clones with uni- (top two) or multi-lineage (bottom two) characteristics.
e, Barcode homoplasy in LARRY clones. The UMAP plot shows that barcode homoplasy results in LARRY clones, such as Clone 71, consisting of multiple subclones defined by distinct mitochondrial variants.
f, Pseudotime analysis reveals distinct trajectories of lineage differentiation during HEMO development. EC and EMP were distributed along two separate late branches of the developmental trajectory.
g, The bar chart demonstrates that mitochondrial variants significantly increase the proportion of cells committed to specific fates.

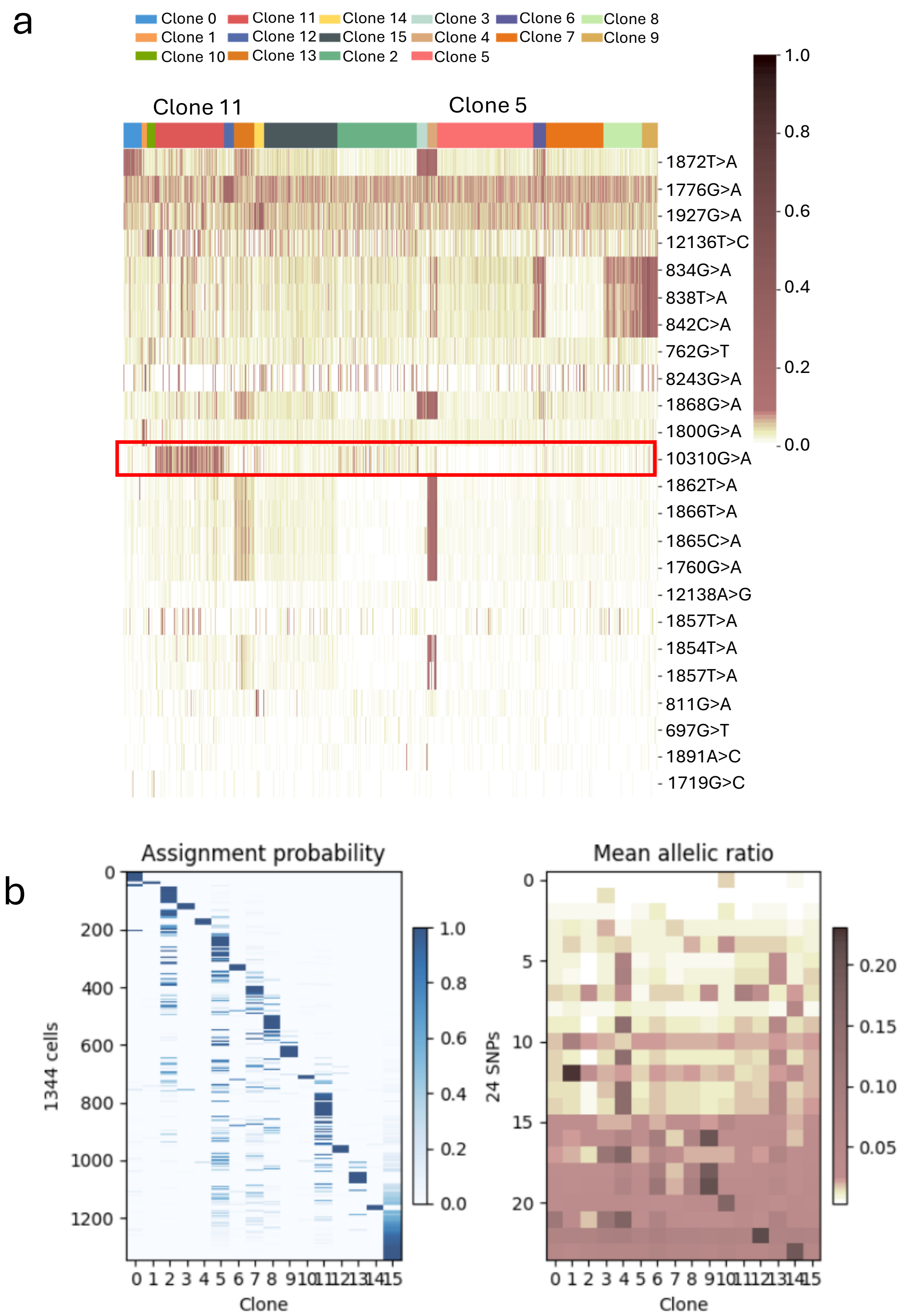


**Fig. S27 Validation of MAESTER-MQuad pipeline in spatial context.**
a, Allele frequency heatmap shows the 24 informative mtDNA SNVs detected by MQuad after MAESTER enrichment in each clone in of chondrosarcoma Visium-MASTER. Mitochondrial SNV 10310 A>G is shown as the informative SNV for identification of big Clone 11 with high allele frequency and Clone 5 with no variation at 10301 position. Each row is a variant, each column is a barcode. Heatmap color indicates the value of the allele frequency.
b, Clones and SNP alleic frequency of chondrosarcoma inferred from Visium-MAESTER.
(Left) Heatmap shows barcode assignment with informative mtDNA variants detected with MQuad. Each row is a cell, each column is a barcode, heatmap color indicates assignment probability.
(Right) Allele frequency heatmap shows the 24 informative mtDNA SNVs detected by MQuad ranked from lowest to highest score of difference in Bayesian Information Criterion (ΔBIC) in each clone. Each row is a variant, each column is a barcode. Heatmap color indicates the value of the ΔBIC which is an indicator for clonal informativeness of each SNP with higher ΔBIC being more informative.

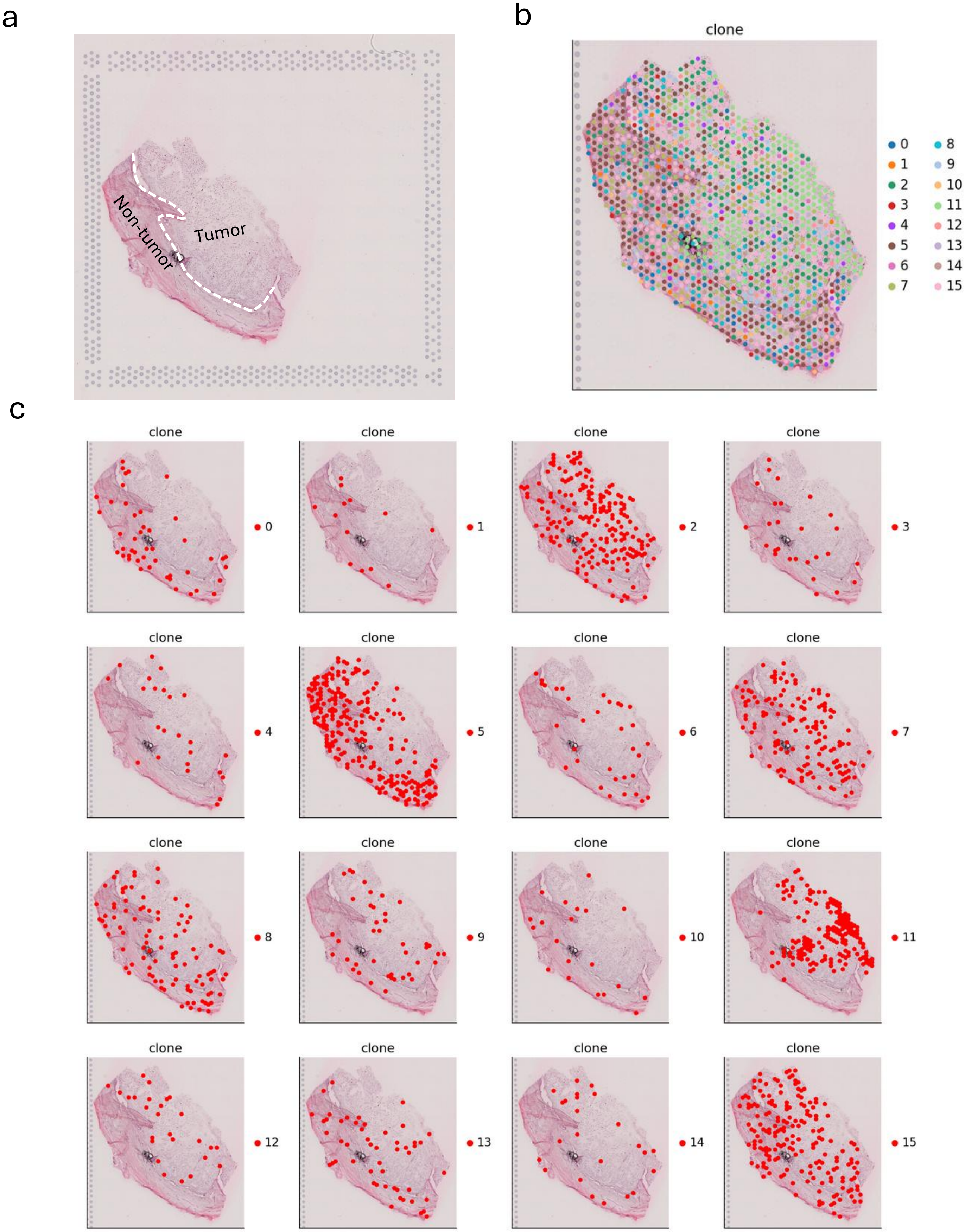


**Fig. S28 Histoclonal relationship of chondrosarcoma inferred from Visium-MAESTER.**
a, Histological image shows the tumor adjacent normal area and neoplastic tumor area.
b, Slide image shows the spatial localization of all 16 clones of chondrosarcoma.
c, Histoclonal mapping demonstrates that Clone 5 distributes in the tumor adjacent normal area and Clone 11 distributes in the neoplastic tumor area.

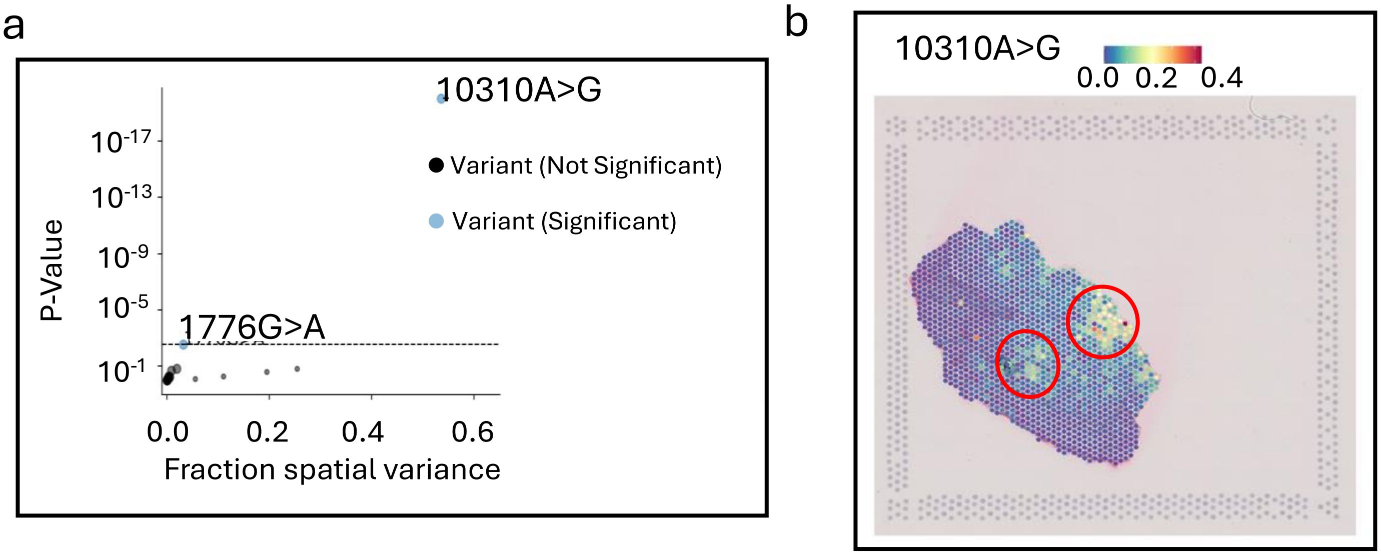


**Fig. S29 The identification of 10310A>G as a significant spatially dependent variable in D15 HEMOs.**

a, Fraction of variance explained by spatial variation versus the adjusted significance of spatial variation (SpatialDE −log P value) for all informative mitochondrial SNVs in chondrosarcoma indicating significant dependance of 10310A>G on spatial location.

b, The spatial allelic frequency of mitochondrial SNV 10310A>G in chondrosarcoma shows its significant dependance on spatial coordinates via SpatialDE.

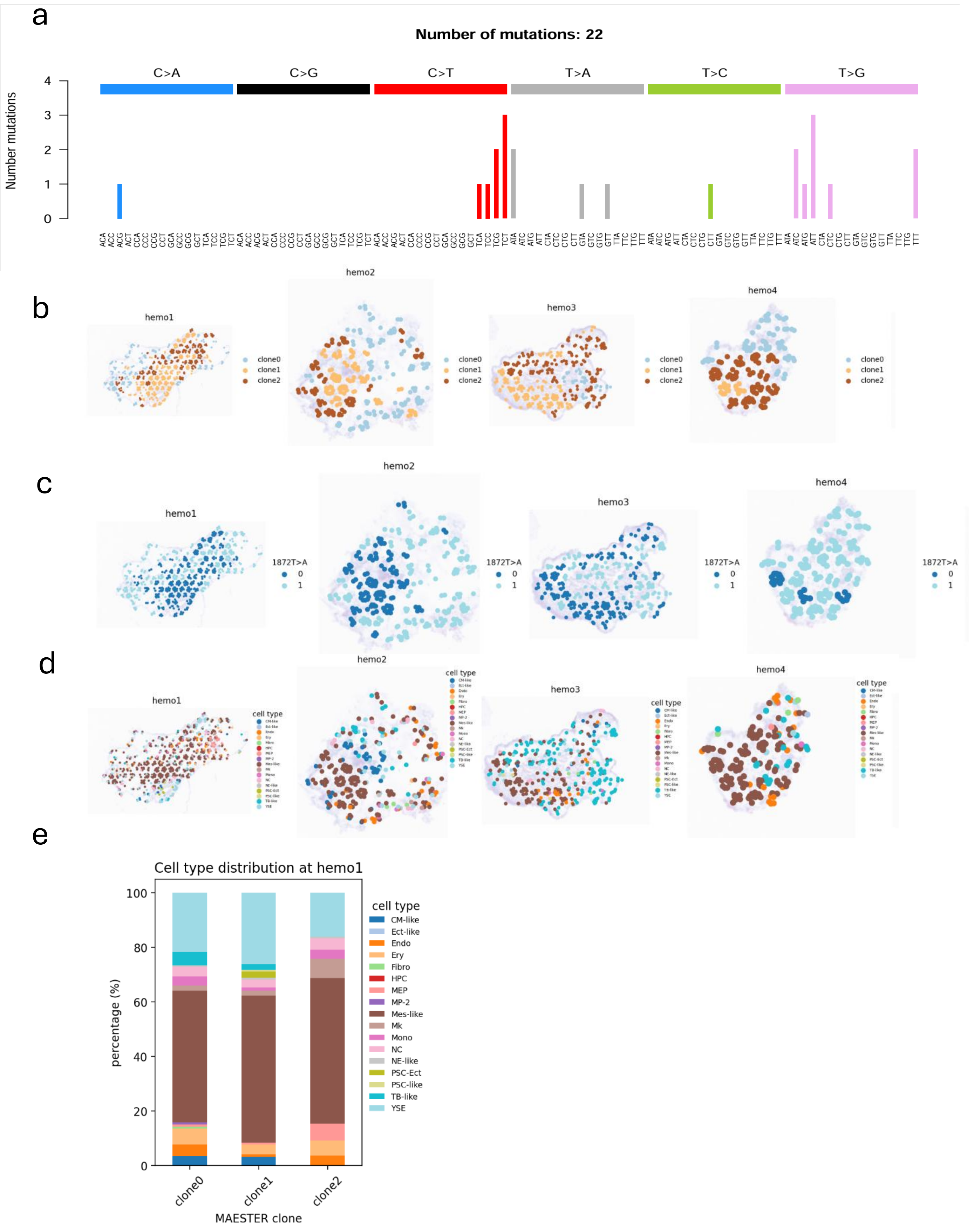

a
Number of mutations: 22
C>A
C>G
C>T
T>A
T>C
T>G
Number mutations
b
hemo1
hemo2
hemo3
hemo4
clone0
clone1
clone2
c
1872T>A
0
1
d
cell type
e
Cell type distribution at hemo1
percentage (%)
MAESTER clone
clone0
clone1
clone2
CM-like
Ect-like
Endo
Ery
Fibro
HPC
MEP
MP-2
Mes-like
Mk
Mono
NC
NE-like
PSC-Ect
PSC-like
TB-like
YSE

**Fig. S30 Spatial clonal dynamics in HEMO #1 to #4  at D15.**
a, Mutational signatures of each mtDNA SNV identified across HEMO #1 to #4.
b, Spatial architecture of 3 clones identified by mtDNA SNVs from MQuad in Visium-MAESTER library in HEMO #1 to #4.
c, Allelic frequency of mtDNA SNV 1872T>A (SpatialDE) aligns with the spatial distribution of Clone 2 across all HEMOs, demonstrating coupling of mitochondrial genetic variation and clonal territoriality.
d, Spatial transcriptomic maps of HEMO #1 to #4, annotated by cluster identity.
e, Lineage dynamics in HEMO #4: Progressive enrichment of erythro-myeloid progenitors (EMP) and megakaryocyte (Mk) populations from Clone 1 to Clone 2, alongside late-stage trophoblast-like (TB-like) differentiation from Clone 2 to Clone 0, reflecting hematopoietic maturation trajectories.

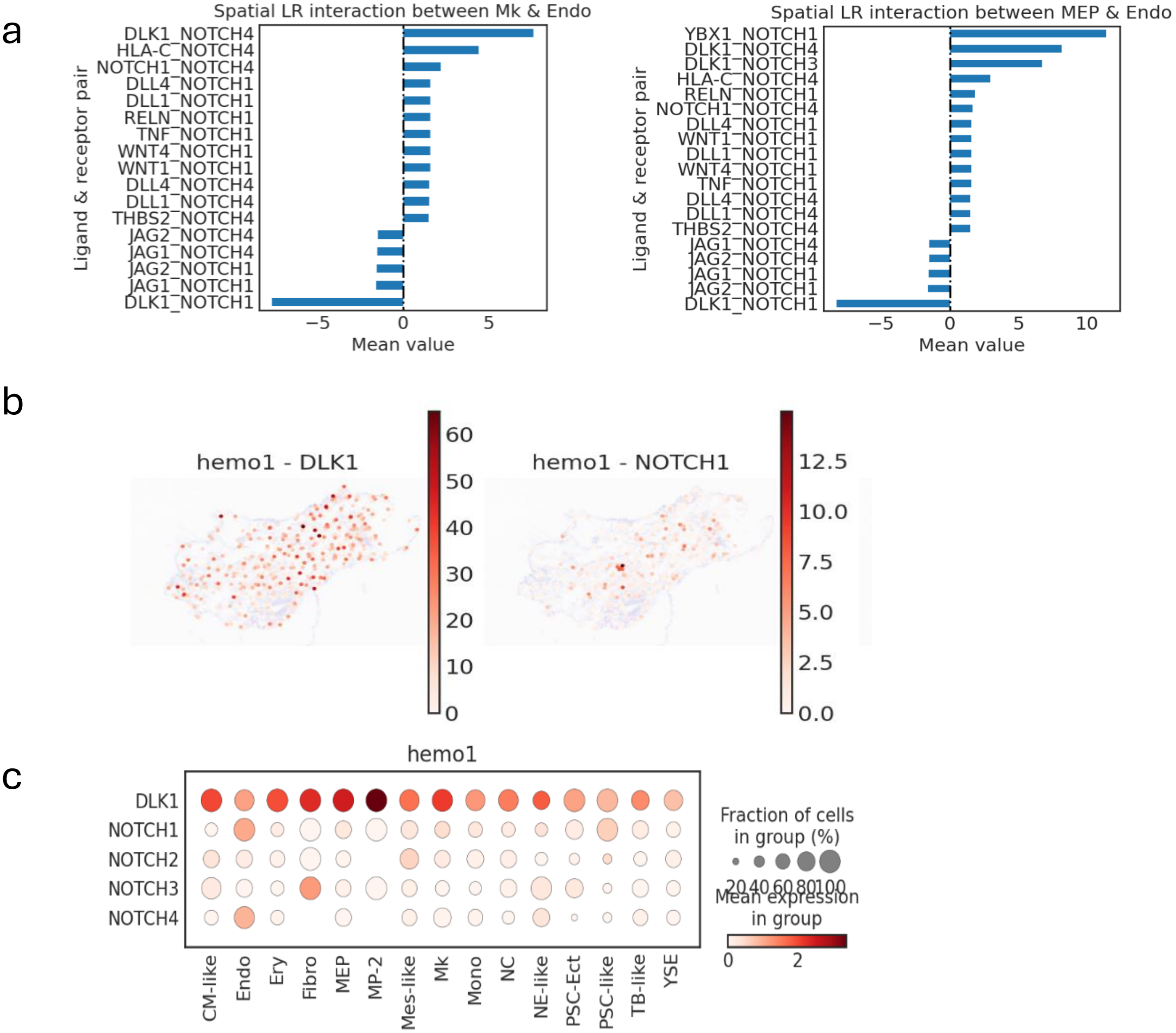


**Fig. S31 Spatially resolved DLK1-NOTCH1 signaling drives lineage differentiation in HEMO clones (D15).**

a, Spatial cell-cell interaction analysis conducted by CellPhoneDB within Clone 2 between EC and Mk, as well as EC and EMP. Pairs with a mean value above 0 indicate the activation of the ligand-receptor pairs. Pairs with a mean value below 0 indicate the inhibition of the pairs. Among these, *DLK1_NOTCH1* exhibit high enriched score between EC and Mk, as well as EC and EMP in Clone 1.

b, Gene expression pattern of *DLK1* and *NOTCH1* within Clone 2 in HEMO #1 in a spatial slice. *DLK1* highly expressed in Mk and EMP within Clone 2. *NOTCH1* highly expressed in EC within Clone 1.

c, Dotplot shows the expression of *DLK1* and *NOTCH1* across various cell types.
